# Induced ubiquitination of the partially disordered Estrogen Receptor alpha protein via a 14-3-3-directed molecular glue-based PROTAC design

**DOI:** 10.1101/2025.11.17.688825

**Authors:** Carlo J. A. Verhoef, Charlotte Crowe, Mark A. Nakasone, Aitana DeLaCuadra-Basté, Tessa Harzing, Naomi A. S. Span, Gajanan Sathe, Kentaro Iso, Christian Ottmann, Luc Brunsveld, Alessio Ciulli, Peter J. Cossar

**Author notes:** These authors contributed equally: Carlo J. A. Verhoef, and Charlotte Crowe. Corresponding authors (A.C) and (P.J.C).

## Abstract

Proteins lacking defined ligandable pockets remain challenging drug targets. Here, we develop a molecular glue-based PROTAC (^MG^PROTACs) approach that chemically conjugates a molecular glue stabilizer to a VHL-recruiting ligand to capture and ubiquitinate the 14-3-3/Estrogen Receptor α (ERα) complex. Our designed ^MG^PROTACs engage a composite interface between 14-3-3 and the disordered F-domain of ERα, promoting cooperative complex formation and target ubiquitination. Biophysical characterization revealed distinct linker-dependent cooperativities across the ^MG^PROTAC series, which influenced both cellular permeability and ubiquitination efficiency. Cryo-EM of the most cooperative ^MG^PROTAC uncovers de novo VHL–14-3-3ζ contacts, while molecular dynamics simulations rationalize the stabilizing interactions underlying cooperativity. Strikingly, fine-tuning linker design enables selective ubiquitination of distinct complex subunits. These findings establish a structural and mechanistic framework for integrating molecular glue and PROTAC principles, expanding the scope of drug discovery to previously intractable protein complexes.

## Main

Induced proximity has enabled precise chemical control over protein–protein interactions (PPIs) to elicit defined biochemical outcomes. Rather than simply inhibiting protein activity, small-molecules that bring proteins together modulate function through processes such as PPI stabilization^1–3^, or signaling modulation.^4^ Conventionally, this is achieved by either bifunctional molecules e.g. proteolysis-targeting chimeras (PROTACs) that are made of two separate protein-binding ligands, or by so-called “molecular glues” that typically bind to either of the two proteins to enhance their complex formation (Fig. 1a).^5^ The most prominently explored modality illustrating this concept is targeted protein degradation (TPD), where the ubiquitin-proteasome system (UPS) is harnessed to chemically induce ubiquitination and subsequent proteolysis of a protein of interest.^6^ Over 30 TPD molecules are currently in clinical evaluation^7–9^, with several TPD molecules demonstrating efficacy against previously intractable targets such as KRAS^10^, STAT5^11^, and LRRK2^12^. Together, these advances highlight the broad potential of protein degradation as a powerful paradigm in drug discovery.

**Figure 1.**
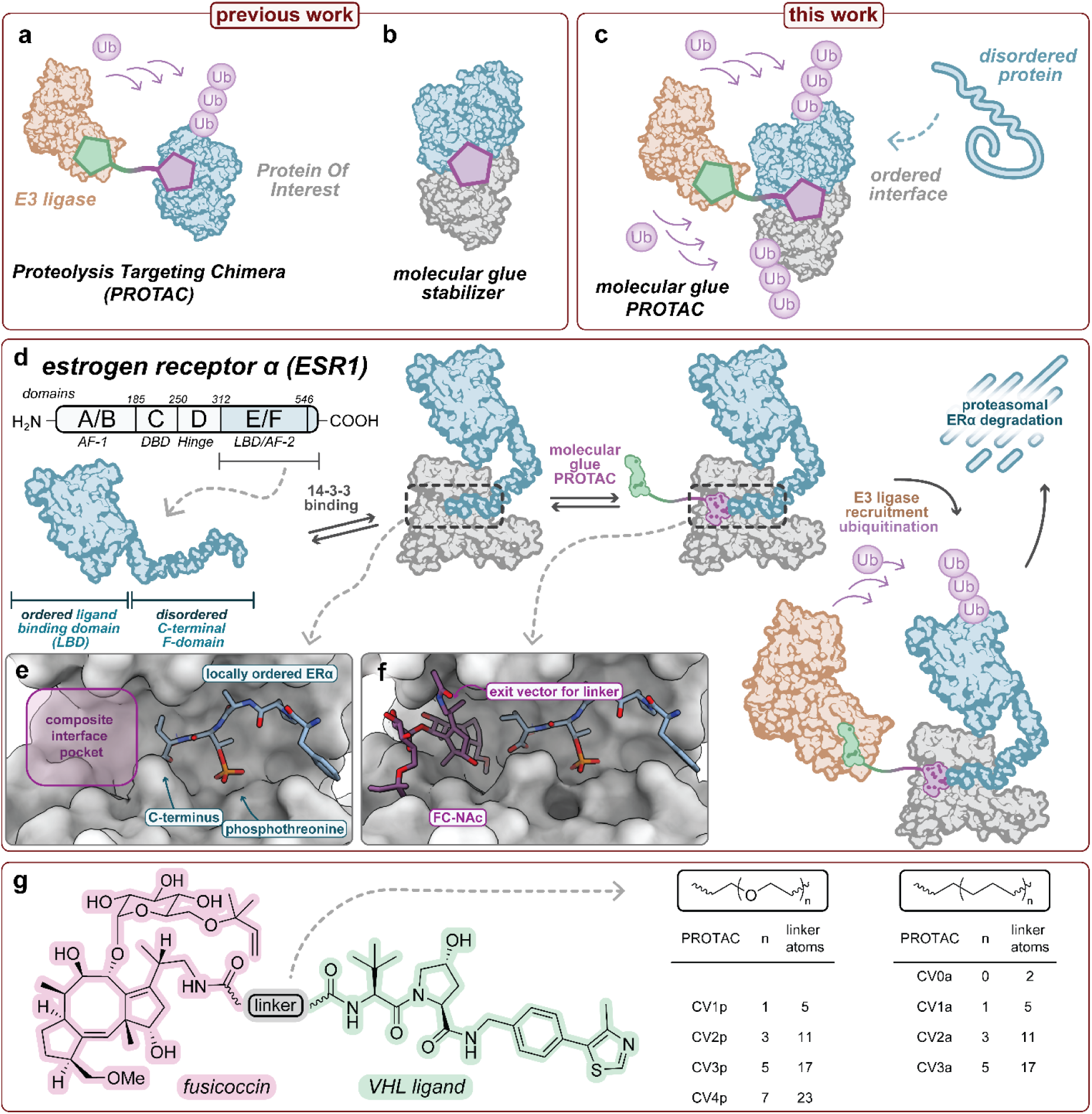
Concept and design of ^MG^PROTACs. (a-b) Schematic illustrates the difference between heterobifunctional PROTACs and monovalent molecular glue induced proximity. (c) Illustration of our approach which replaces a protein of interest ligand with a molecular glue within a PROTAC to produce a molecular glue PROTAC (^MG^PROTAC) that can ubiquitinate a disordered protein complex. (d) ERα was selected as the target protein for ubiquitination, using the FC-NAc ligand as a 14-3-3/ERα molecular glue. (e) Crystal structure of the binary 14-3-3/ERα complex reveals the composite interface and a pocket for small molecule stabilization (PDB: 7NFW). (f) Crystal structure of the ternary 14-3-3/ERα/FC-NAc complex, showing the exit vector for ^MG^PROTAC development (PDB: 8BZH). (g) Overview of the ^MG^PROTAC library, featuring PEG and alkyl linkers.

Beyond degradation, induced proximity has expanded to encompass diverse heterobifunctional strategies that recruit other enzymatic machineries—including kinases, phosphatases, deubiquitinases, and lysosomal components—to alter protein function.^13–16^ Further, molecular glue stabilizers, which act independently of degradation, restore or enhance weak protein–protein interfaces (Fig. 1b.^3,17,18^ The potential of molecular glue stabilization has been demonstrated by Revolution Medicines’ Zoldonrasib, which glues KRAS^G12D^ (ON) to CYPA^3^, and 14-3-3 glues that selectively target specific 14-3-3 complexes.^2,19,20^ Collectively, these strategies vastly expanding the chemical space for modulating protein behavior.

The efficacy of induced proximity often arises from the cooperative assembly of higher-order complexes^21^, in which activity depends on the formation and stability of multicomponent assemblies rather than on high-affinity ligand binding.^22^ Such cooperativity enhances potency and dictates selectivity through the emergent network of PPI contacts within the induced complex, observed structurally and biophysically with both bifunctional^22–26^ and monovalent glue molecules.^2,27–31^ However, most current strategies remain confined to protein targets bearing ligandable surfaces, while many clinically relevant proteins, such as RNA binding proteins, transcription factors and scaffolding proteins, lack ligandable pockets or surfaces.

Here, we establish a framework for chemically controlled ubiquitination of a protein complex by integrating a molecular glue stabilizer as the target-recruiting ligand in a PROTAC design to develop a molecular glue PROTAC (^MG^PROTAC; Fig.1c). In our system, the molecular glue functions as the ligand for a protein complex, stabilizing an interface between a partially disordered target and a scaffold protein, enabling the conditional recruitment of the complex to an E3 ligase. We demonstrate this concept by recruiting the von Hippel–Lindau (VHL) E3 ligase to a complex between the partially disordered Estrogen receptor α (ERα) and the hub protein 14-3-3. We found that the degree of cooperative VHL recruitment is strictly influenced by linker composition and length. Structural analysis using cryogenic electron microscopy (cryo-EM) reveals a de novo VHL/14-3-3 interface formed through induced hydrogen bonds and hydrophobic contacts. Recruitment requires both 14-3-3 and ERα, and promotes selective compound-dependent ubiquitination whose pattern and extent could be precisely modulated by tuning linker properties.

## Results and Discussion

### Design and synthesis of 14-3-3/ERα complex ^MG^PROTACs

To explore small-molecule–induced degradation of a protein complex, we conjugated the 14-3-3/ERα molecular glue 3’-deacetylated fusicoccin N-acetyl (FC-NAc)^32^ to the VHL ligand VH032^33^ through a series of chemical linkers (Fig. 1d–e). ERα comprises both structured and intrinsically disordered regions; with the N-terminal AF-1 domain, hinge region, and C-terminal F-domain existing as unstructured, whereas the DNA-binding (DBD) and ligand-binding (LBD) domains adopt well-defined tertiary structures (Fig 1d). FC-NAc enhances 14-3-3 binding to the disordered C-terminal F-domain of ERα, which undergoes a disorder-to-order transition upon 14-3-3 engagement with phosphorylated T594 (pT594), generating a composite interface stabilized by FC-NAc (Fig. 1 e–f).^32,34^ ERα remains a clinically relevant target in ERα–positive breast cancer, where existing small molecules targeting the ligand-binding domain (LBD) are susceptible to resistance conferred by recurrent LBD mutations (e.g., Y537S, Y537N, D538G).^35^ Taken together, this presented an interesting biomolecular case study to explore VHL recruitment and ubiquitination of a partially disordered protein 14-3-3 complex.

^MG^PROTAC design began with analysis of the crystal structure of the 14-3-3/ERα/FC-NAc complex (Fig. 1f) (PDB: 8BZH)^32^, which identified the amide modification site of FC-NAc as a suitable exit vector for PROTAC extension. As the acetyl group projects into the solvent, we hypothesized that this position would tolerate linker attachment without disrupting 14-3-3/ERα binding. To optimize cooperative E3 ligase recruitment to the 14-3-3/ERα complex an eight-member library of FC-NAc–VH032 chimeras incorporating linker lengths ranging from 2 to 23 atoms and two distinct chemical compositions—polyethylene glycol (PEG) or alkyl were synthesized (Fig 1g). The ^MG^PROTAC libraries were prepared via a six-step convergent synthesis (SI Scheme 1), starting from the fungal extract 3’-deacetylated fusicoccin-A. Notably, the final products contained a partial cleavage (5–43%) of the 2-methylbut-3-en-2-yl ether moiety on the fusicoccane ligand, however, this modification had negligible impact on 14-3-3/ERα stabilization (Fig. S1).

### ^MG^PROTACs stabilize the 14-3-3/ERα PPI and engage VHL in vitro and in cellulo

With the bifunctional library in hand, we examined the biophysical properties of the ^MG^PROTAC libraries, specifically, the influence of linker length and hydrophobicity. ^MG^PROTAC binding and 14-3-3/ERα complex stabilization were assessed using a fluorescence anisotropy (FA) assay. To assess ^MG^PROTAC binding each compound was titrated to a 14-3-3/ ERα protein-peptide complex (Fig. 2a). The monovalent molecular glue FC-NAc served as a positive control (EC₅₀ = 0.34 µM). All ^MG^PROTAC, except for CV3a, displayed dose-dependent binding (EC₅₀ of 0.59*—*1.49 µM; Fig. 2a). CV3a, which features a 17-atom aliphatic linker, exhibited substantially weaker binding (EC₅₀ ≈ 40 µM). Further, the VHL ligand VH032 does not engage the 14-3-3/ERα complex. These findings validated that the N-acetyl exit vector of FC-NAc was compatible with linker derivatization.

**Figure 2.**
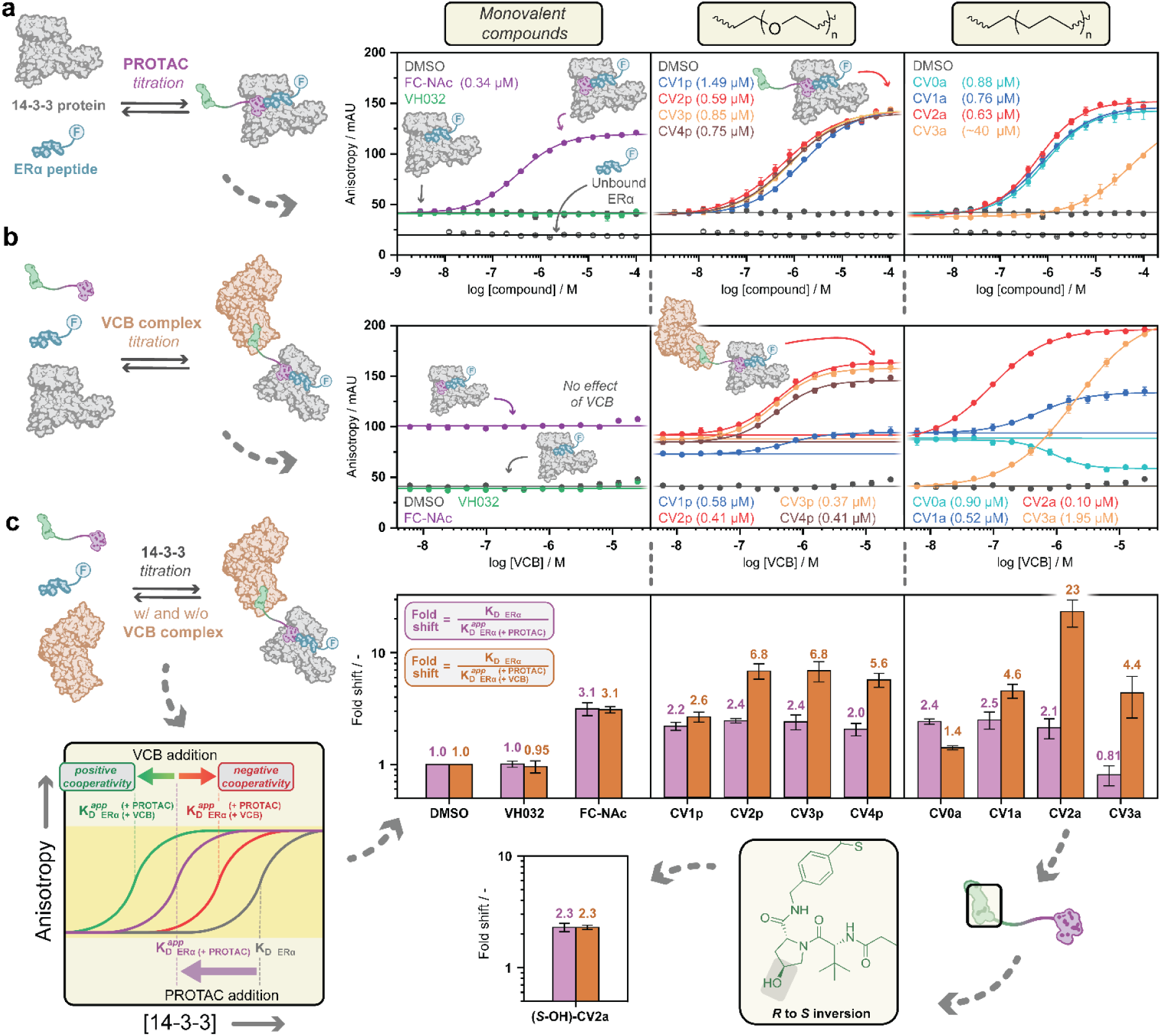
Stabilization of the 14-3-3/ERα protein complex and cooperative binding of VCB. (a) Fluorescence anisotropy assay data of PROTAC (or monovalent compound) titration to 10 nM fluorescently labeled ERα 8-mer phosphopeptide and 1 µM 14-3-3, with DMSO serving as negative control. EC50 values of binding curves are indicated. (b) Titration of the VCB protein complex to 10 nM fluorescently labeled ERα peptide, 500 nM 14-3-3, and 1 µM PROTAC (or monovalent compound). EC50 values of binding curves are indicated. (c) Conceptual overview and results presented in a barchart of 14-3-3 protein titration to 10 nM fluorescently labeled ERα peptide, 1 µM PROTAC (or monovalent compound), and in the presence (5 µM, orange bars) or absence (purple bars) of VCB protein complex. Error bars represent standard deviations of independent scientific replicates (n = 3).

^MG^PROTAC stabilization of the binary 14-3-3/ER*α* complex (K_D_ = 2.32 µM) was then characterized by 14-3-3 titration in the absence or presence of 1 µM ^MG^PROTAC (Fig. S2). All compounds, except CV3a, enhanced ERα binding to 14-3-3, yielding enhanced apparent K_D_ (K_D_^app^) values of 0.96*—*1.14 µM. Both linker series provided stabilization only marginally weaker than FC-NAc (0.75 µM), indicating the ^MG^PROTAC retain the molecular glue stabilization.

Next, we assessed ^MG^PROTAC engagement with the VHL/Elongin C/Elongin B (VCB) complex using two assays; a FA HIF1α displacement assay—the native VHL substrate— and an in cell VHL target engagement assay.^36^ All ^MG^PROTACs displaced the HIF1α peptide in the FA assay (IC₅₀ ranged from 0.43*—*1.41 µM), except CV3a, which showed minimal displacement (IC₅₀ ≥ 100 µM) (Extended Fig 1a).

In cell VHL engagement was evaluated using a bioluminescence resonance energy transfer (BRET) assay in ERα-positive cells (MCF-7). Cells were transiently transfected with NanoLuc–VHL, treated with a VHL fluorescent probe and incubated with compound (Promega, N2931). In permeabilized cells, ^MG^PROTACs elicited an analogous trend to the FA-based HIF1α displacement assay (Extended Fig. 1b). In non-permeabilized MCF-7 cells, differential permeability was observed. Of the two ^MG^PROTAC series, the alkylic-linkers showed improved cell penetration over the polar PEG-linkers, except for CV0a and CV3a (Extended Fig. 1c). The poor engagement of CV0a was likely due to its short linker, which increases overall polarity and limits membrane diffusion. Further, we hypothesize the lowered activity of CV3a originates from the hydrophobic character of the long carbon linker, which potentially affects solubility and ligand engagement. Combined, the results of the FA assays and VHL target engagement experiments demonstrate that the ^MG^PROTAC effectively engage both 14-3-3/ERα and VHL complexes.

### ^MG^PROTACs cooperatively recruit VHL to the 14-3-3/ERα complex

Recruitment of the VCB complex to the 14-3-3/ERα complex was next investigated. A VCB titration was performed at a fixed concentration of compound, and 14-3-3/ERα complex. In the presence of the vehicle (DMSO), FC-NAc, or VH032 no response was elicited (Fig. 2b). This indicates that VCB, whether in an unbound or VH032-bound state, has no natural affinity for the 14-3-3/ERα complex. VCB titration in the presence of the ^MG^PROTACs yielded a direct effect on ERα binding, consistent with favorable quaternary complex formation. Structure-activity relationship (SAR) analysis revealed distinct linker-dependent effects on ^MG^PROTAC-induced VCB recruitment. PEG-based ^MG^PROTACs displayed weak to moderate positive recruitment, with the shortest linker (CV1p) giving the smallest effect. In contrast, the shortest alkyl-linker (2 atoms) based ^MG^PROTAC (CV0a) disrupted 14-3-3/ERα complex formation, indicative of negative cooperativity. This likely reflects steric limitations imposed by the insufficient linker length. Extending the alkyl linker (CV1a–CV3a) restored cooperative recruitment, with CV2a and CV3a showing the strongest responses across both linker series (EC_50_ = 0.10 and 1.95 μM, respectively, Fig. 2b).

Having established that seven of the eight ^MG^PROTACs recruit VCB to the 14-3-3/ERα complex, we assessed glue-like enhancement of 14-3-3/ERα avidity. FA-based 14-3-3 titrations were performed in the presence of a ^MG^PROTAC (1 µM) and VCB (5 µM). At these concentrations >95% of the ^MG^PROTAC is bound to the VCB complex, enabling the detection of cooperative interactions between the VCB/^MG^PROTAC complex and the 14-3-3/ERα complex. All ^MG^PROTACs stabilized the 14-3-3/ERα complex, except negatively cooperative CV0a, consistent with the VCB titration assay (Fig. 2c, Fig. S3). The greatest effects were observed for compounds with 11- and 17-atom linkers (CV2a and CV3a), showing 23- and 4.4-fold stabilization, respectively. A two-dimensional FA titration^21^ varying both CV2a and VCB revealed cooperative interplay between ^MG^PROTAC and VCB concentration (Extended Fig 2a-d). CV2a titration dose-dependently enhanced ERα binding to 14-3-3 (up to 2.5-fold), which was further amplified by VCB, reaching over 23-fold enhancement at high CV2a and VCB concentrations. These findings show that VCB binding can promote an over 11-fold improvement in the apparent affinity (K_D_^app^) of ERα.

To validate the observed K_D_ shifts were due to VCB binding, we synthesized an epimer control of CV2a bearing inverted stereochemistry at the proline 4-position of the VHL ligand, designated as (*S*-OH)-CV2a, which disrupts VCB binding (Extended Fig. 2e-f).^37^ As expected, (*S*-OH)-CV2a stabilized the 14-3-3/ERα complex similarly to CV2a, but failed to recruit VCB (Fig. 2c, Extended Fig 2f). Finally, we examined VCB recruitment by CV2a across all seven human 14-3-3 isoforms using the previously described 14-3-3 titration (Extended Fig. 3). Although 14-3-3 isoforms share a highly conserved binding groove and engage ERα similarly, their peripheral regions differ, potentially influencing VCB interaction. CV2a-mediated 14-3-3/ERα stabilization was relatively consistent across isoforms (1.2–3.8-fold), but VCB recruitment varied substantially, ranging from 5.6-to 176-fold. The σ isoform showed the weakest VCB stabilization (5.6-fold), τ, β, γ, and η displayed moderate effects (29-, 34-, 48-, and 61-fold), while ζ and ε exhibited the strongest stabilization (109- and 176-fold, respectively). These findings show both linker length and chemical composition govern the cooperative stabilization. Specifically, longer apolar linkers promote strong positive cooperativity up to an optimal length (CV2a, 11-atoms). The results also support formation of a *de novo* VCB–14-3-3/ERα interface, which is influenced by isoform-specific residues outside the conserved binding groove.

### Cooperative VHL recruitment extends to the 14-3-3/ERα^(LBD_F)^ protein-protein complex

Using 14-3-3 and a ERα^(LBD_F)^ construct (Fig S4)—which includes the ligand-binding domain (LBD) and disordered F domain containing phosphorylated T594—we assessed ^MG^PROTAC recruitment of the 14-3-3/ ERα^(LBD_F)^ protein-protein complex to VCB using the HIF1α displacement assay.^38^

Cotreatment with 14-3-3/ERα^(LBD_F)^ protein-protein complex and either CV2a or CV3a significantly enhanced VCB displacement by the ^MG^PROTACs, compared to ^MG^PROTAC alone, reducing the IC₅₀ for HIF1α displacement by 2.8-fold and >34-fold, respectively (Fig. 3a and Extended Fig 4a). In contrast, the negatively cooperative ^MG^PROTAC CV0a decreased HIF1α displacement in the presence of the complex, increasing the IC₅₀ by 4.0-fold, demonstrating ^MG^PROTACs engage the full-length 14-3-3/ERα protein complex. This represents one of the first examples of controlled chemical-induced recruitment of an E3 ligase to a protein complex.

**Figure 3.**
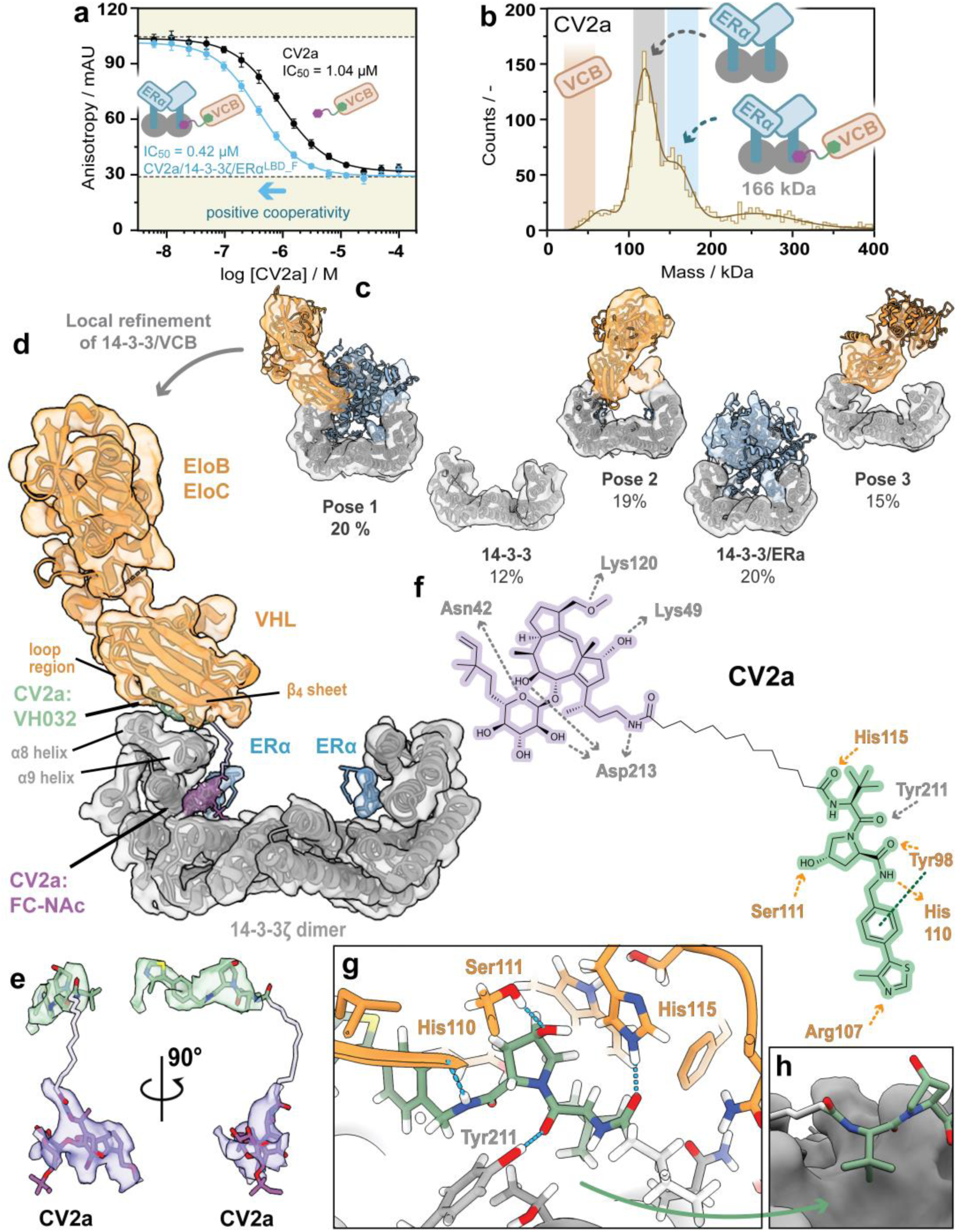
Structure of the VCB/CV2a/14-3-3ζ/ERα^(LBD_F)^ complex. (a) FA assay data of CV2a titration to 25 nM fluorescently labeled HIF1α peptide and 50 nM VCB protein complex in the absence (black) or presence of 10 µM 14-3-3/ERα^(LBD_F)^ complex (blue). Error bars represent standard deviations of independent scientific replicates (n = 3). (b) Mass photometry data of a 1:1 mixture of the 14-3-3/ERα^(LBD_F)^ complex (2:2 stoichiometry, 124 kDa) and the VCB complex (41 kDa) with three equivalents of CV2a. (c) 3D classes obtained displaying the conformational and compositional landscape of the sample. (d) Cryo-EM map and model of the best resolved VCB/CV2a/14-3-3ζ/ERα species showing a 1:1:2:2 stoichiometry. (e) Local volume of the locally refined cryo-EM map corresponding to CV2a. (f) Residues on VHL and 14-3-3 which were shown to form contacts with CV2a in the MD simulation. Hydrogen bonds are shown for VHL (orange) and 14-3-3 (grey). The pi-stacking interaction with VHL Tyr98 is shown in dark green. (g) Selected residues making contact with VH032, including Tyr211 from 14-3-3. (h) The tert-butyl group on the VH032 moiety of CV2a occupies a pocket on the 14-3-3 surface.

### Cooperativity is driven by favourable protein-protein contacts between 14-3-3 and VHL

To gain a structural insight into ^MG^PROTAC cooperativity, we turned to single-particle cryogenic electron microscopy (cryo-EM). Given that the strongest cooperativity and affinity was observed for CV2a, this compound was selected for structural analysis. The multicomponent complex was prepared by incubating VCB, CV2a and the 14-3-3ζ/ERα^(LBD_F)^ complex in a 1.2:1.2:1:1 molar ratio, followed by grid preparation without further purification. Mass photometry analysis revealed a higher-molecular-weight species (∼166 kDa), consistent with asymmetric binding of VCB to the 14-3-3ζ/ERα^(LBD_F)^ dimer-of-dimers assembly with a 2:2:1 14-3-3/ERα/VCB stoichiometry (Fig. 3b and Extended Fig. 4b). Cryo-EM analysis revealed a compositionally and conformationally heterogenous population, with a single VCB complex being recruited to the 14-3-3ζ dimer in multiple orientations (Fig. 3c, Extended Fig. 5). Despite the heterogeneity in the cryo-EM sample, a major species corresponding to VCB positioned ‘in plane’ with the 14-3-3ζ dimer was resolved to ∼4.3 Å resolution, enabling confident model building of VCB and 14-3-3ζ (Fig. 3d, Extended Fig. 6, Fig S5-6). Although volume corresponding to the ERα LBD was present, the LBD was poorly defined (Fig. 3c ‘Pose 1’) likely due to the high degree of flexibility conferred by the ERα F-domain, and only the final four residues of the phosphorylated C-terminal F-domain were modelled.

Strikingly, CV2a induces a neo-protein–protein interface between 14-3-3ζ and VHL (Fig. 3d), with the loop region and β4 sheet of VHL directly engaging the α8 and α9 helices of one 14-3-3ζ monomer—representing the first chemically induced *de novo* interface reported for 14-3-3. Well-defined density was observed for the VH032 and FC-NAc ligands of CV2a (Fig. 3e), which adopted binding modes consistent to the previously reported structures for VH032 bound to VHL and FC-NAc bound to the 14-3-3/ERα complex (Fig 3f, Extended Fig. S7).^32,33^ In contrast, the 11-atom hydrocarbon linker of CV2a has no interpretable volume. We therefore performed energy minimization and 500 ns molecular dynamics (MD) simulation to explore plausible conformations and interactions of the complex. Superimposing the apo-14-3-3ζ AlphaFold structure with our cryo-EM model (Fig. S7) revealed minimal backbone rearrangement. Rotation of Tyr211 of 14-3-3ζ to accommodate the VH032 ligand of CV2a, forming a putative hydrogen bond between the carbonyl oxygen of the CV2a *tert*-leucine moiety was observed (Fig. 3g). Further, the *tert*-leucine moiety of CV2a occupies a pre-existing pocket on 14-3-3ζ (Fig. 3h). Finally, Arg222 of 14-3-3ζ reorients to avoid steric clashes with VHL and engages a hydrogen bond with Asp67 of VHL.

The MD simulation identified an extensive network of stabilizing inter-protein interactions. The 14-3-3ζ Glu208 forms a triad of contacts with the basic sidechains of Arg108 and His110 in the β4 sheet of VHL (Fig. 4a). On the same β4 sheet, VHL Tyr112 appears to hydrogen bond to Tyr211 of 14-3-3ζ (α9 helix), which in turn hydrogen bonds with CV2a. Nearby the 4-methylthiazole moiety of CV2a, Asp204 of 14-3-3 hydrogen-bonds with the carbonyl oxygen of Pro99 located on a loop region on VHL (Fig. 4b). Towards the CV2a alkylic exit point on VH032, we observed VHL Asn67 and 14-3-3ζ Gln219 form a hydrogen bond (Fig. 4c). Further along the α9 helix, 14-3-3ζ Arg222 forms a hydrogen bond interaction with the carboxylic acid group of Asp92 of VHL. Sequence alignment across the seven human 14-3-3 isoforms (β, ε, η, γ, σ, θ, ζ) showed high conservation of the previously described interface residues (Fig. S8). Notably, Asp204 of 14-3-3ζ, which hydrogen bonds with Pro99 of VHL, is substituted by His in 14-3-3σ—introducing potential electrostatic repulsion with nearby VHL Arg107 and providing a structural rationale for the reduced cooperativity observed for 14-3-3σ.

**Figure 4.**
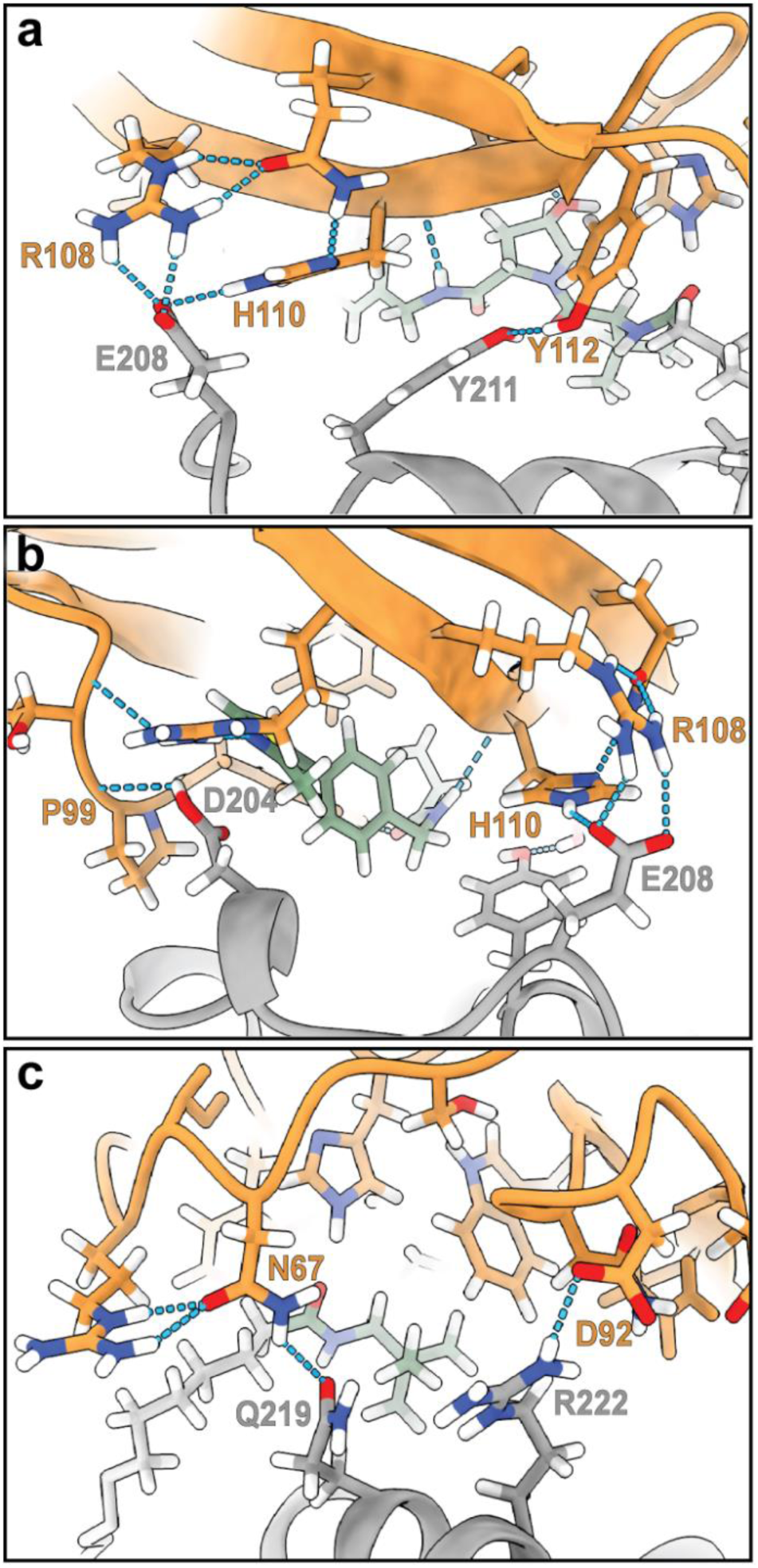
Protein-protein interactions between VHL (orange) and 14-3-3 (grey) based on cryo-EM analysis and MD simulation. Each view is rotated 120 degrees around the vertical axis.

Together, our mass photometry, cryo-EM, and MD analyses reveal a network of favourable interprotein contacts between the VHL β-domain, CV2a, and 14-3-3ζ α8/α9 helices, rationalizing why CV2a displays the strongest cooperativity in the ^MG^PROTAC series. The structural model also suggests that the shorter linkers in CV0a and CV1a (2 and 5 atoms, respectively) are insufficient to span the VH032 and FC-NAc binding sites, consistent with the structure–activity relationships correlating linker length with cooperativity. These findings provide a structural rationale for cooperative binding of CV2a, and potentially CV3a.

### VHL recruitment to 14-3-3 is driven by ERα

Structural analysis of ERα within the complex proved more challenging due to the conformational heterogeneity of ERα, which prevented the reconstruction of a single stable high-resolution structure. Nonetheless, structural analysis revealed that ERα engages with both monomers of the 14-3-3 dimer as volume of the phosphorylated F-domain was observed within both binding grooves of 14-3-3 (Fig. 5a-b, Fig. S9). This observation is consistent with previously published 14-3-3/ERα and 14-3-3/ERα/FC-NAc crystal structures (Fig. 1e-f) and in solution data.^32,38^ Moreover, additional density was detected near helix α8 and α9 of monomer 2 in 14-3-3ζ, likely corresponding to the ERα ligand-binding domain (LBD) (Fig. 5c).

**Figure 5.**
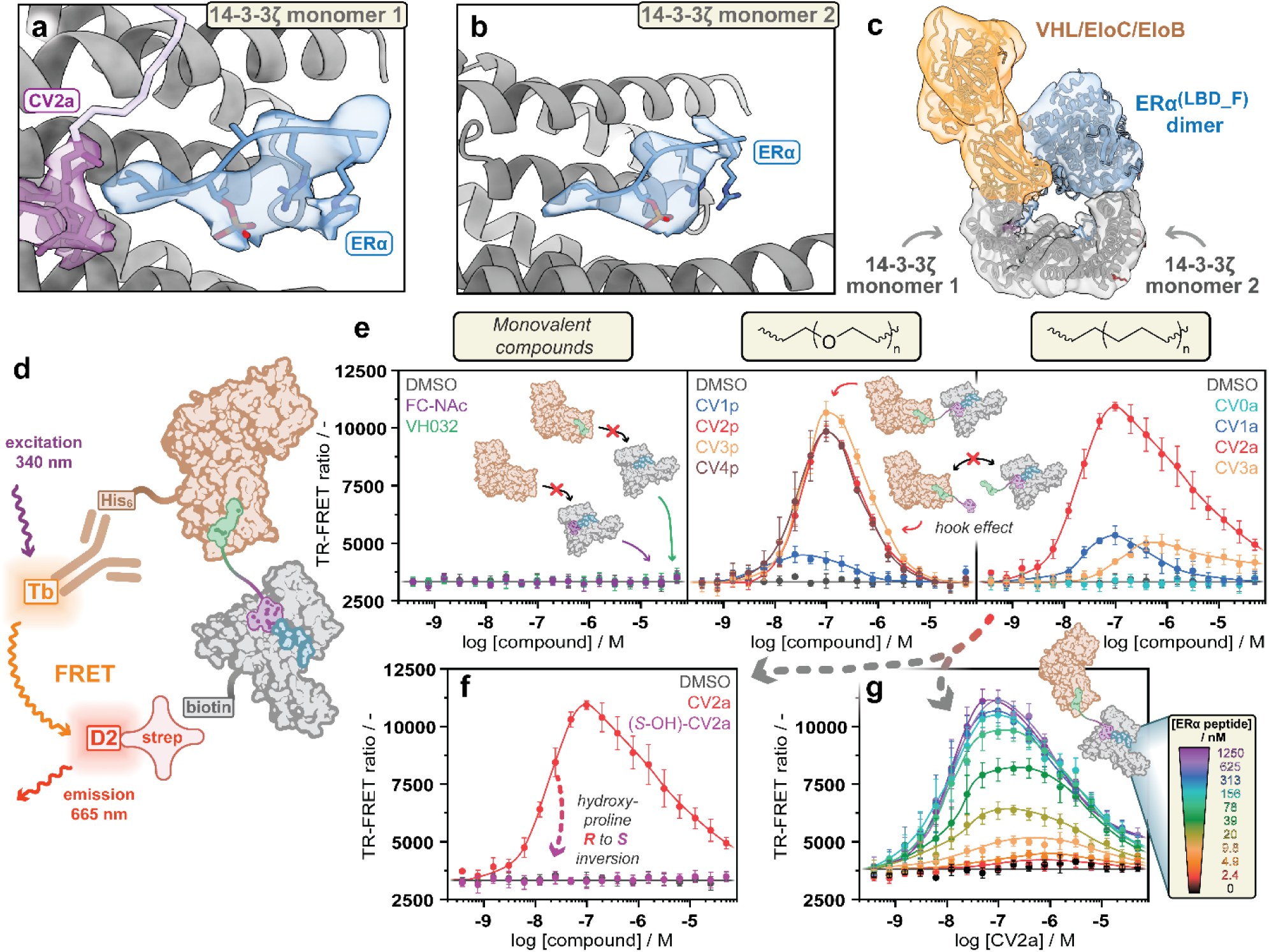
ERα drives quaternary complex formation. (a) Locally refined cryo-EM map from this study showing the volume around the ERα C-terminus associated with 14-3-3ζ monomer 1, and (b) monomer 2. (c) Unsharpened and Gaussian-smoothed cryo-EM map from this study showing volume corresponding to the ER**α** LBD domain. (d) TR-FRET assay setup. (e) TR-FRET assay data of PROTAC (or monovalent compound) titration to 20 µM unlabeled ERα peptide, 50 nM His-tagged VCB complex, 50 nM biotinylated 14-3-3γ, 0.5 nM anti-His Tb-conjugated mAb and 12.5 nM streptavidin D2-conjugate. (f) Comparison between the CV2a and (*S*-OH)-CV2a generated TR-FRET signal. (g) Titration of unlabeled ERα and CV2a. Error bars represent standard deviations of independent scientific replicates (n = 3).

To directly assess recruitment of the VCB complex to 14-3-3, we optimized a time-resolved Förster resonance energy transfer (TR-FRET) assay (Fig. S10), employing unlabeled ERα peptide, His-tagged VCB complex, and biotinylated 14-3-3γ (Fig. 5d). An analogous correlation between linker properties was observed between FA and TR-FRET assay results (Fig. 5). Notably, CV2a exhibited an extended recruitment window that persisted to higher concentrations compared with the CVp series, consistent with its higher cooperativity delaying the onset of the hook effect. In contrast, CV3a—although classified as cooperative—showed markedly reduced recruitment. These lack of induced proximity likely arise from the weaker intrinsic affinity of CV3a for both 14-3-3ζ/ERα and VCB at nanomolar concentrations. Further, control compound (S-OH)-CV2a elicited no TR-FRET signal, indicating induced proximity was directly mediated by the VHL ligand (Fig. 5f). To examine ERα dependence, a two-dimensional titration of ERα and CV2a was performed (Fig. 5g). VCB recruitment was strictly dependent on ERα concentration, with efficient recruitment observed above 10 nM ERα — consistent with the proposed mechanism in which 14-3-3 acts as a scaffold to mediate E3 ligase recruitment to ERα. Together, this data shows that VCB recruitment is ERα-dependent eliciting high selectivity for the 14-3-3/ERα complex over the apo-14-3-3.

### CV2a-dependent VHL recruitment drives ubiquitination of the 14-3-3ζ/ERα^(LBD_F)^ complex

As we could not detect measurable degradation of ERα in ERα-positive MCF7 cells treated with CV2a, we turned our attention to investigating the preceding step in the mechanism, and moved to examine whether the 14-3-3ζ/ERα^(LBD_F)^ complex undergoes ubiquitination in vitro. Briefly, the 14-3-3ζ/ERα^(LBD_F)^ complex was incubated with recombinantly expressed and purified UBE1, UBE2D2, UBE2R1, CRL2^VHL^ and ubiquitin in the presence of ^MG^PROTACs and analysed time-dependently using Coomassie-stained SDS– PAGE (Fig. 6a).

**Figure 6.**
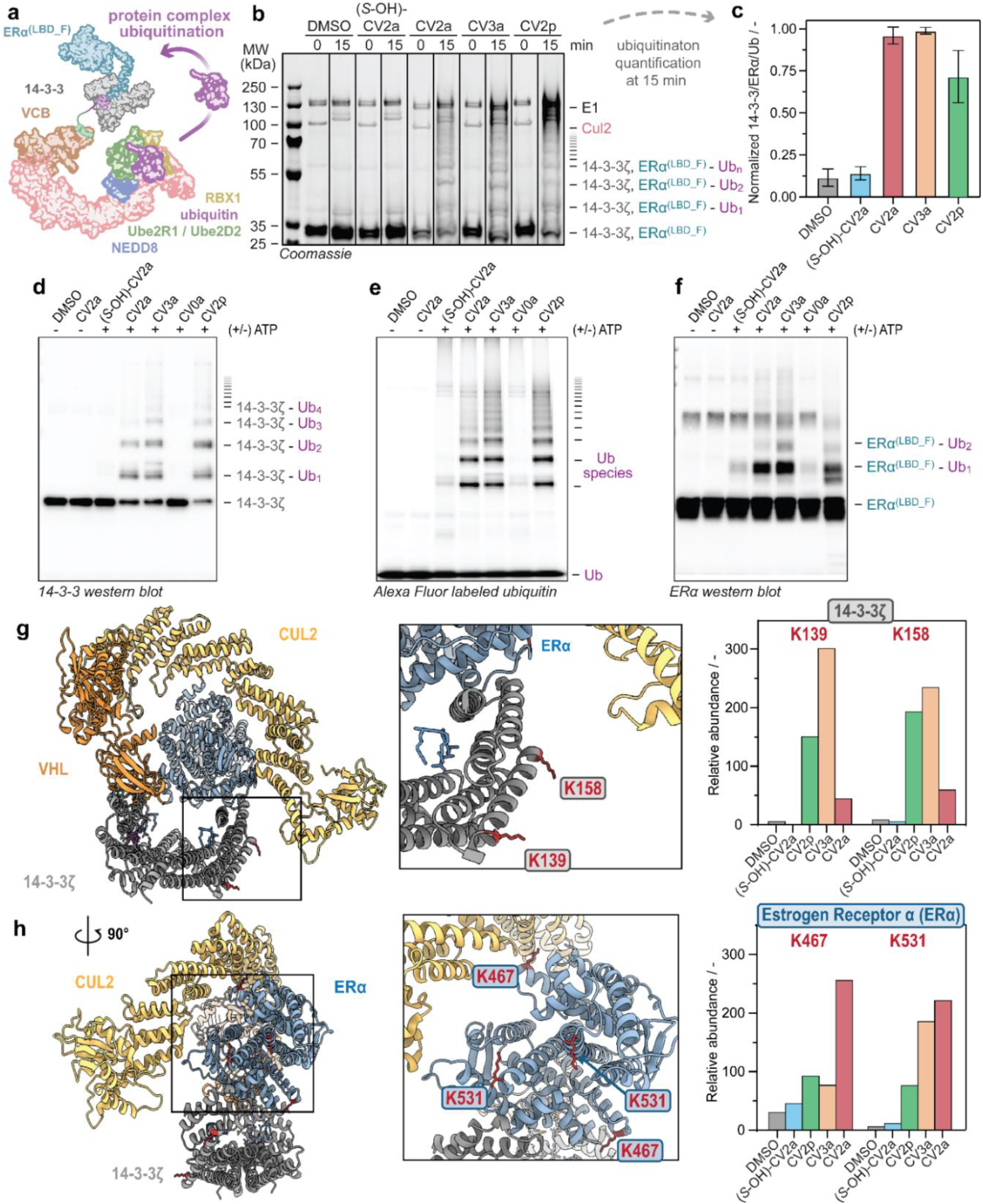
Ubiquitination of the 14-3-3/ERα complex. (a) Schematic of the ubiquitination reaction. (b) Coomassie gel results over time, extended timepoints in Fig. S11. (c) Quantification of ubiquitination at 15 min. Error bars represent standard deviations of independent scientific replicates (n = 3). (d) 14-3-3 western blot, (e) in-gel Alexa Fluor fluorescence of labeled ubiquitin, and (f) ERα western blot of in vitro ubiquitination assay at 24 hours. Mass spectrometry results after 15 min incubation indicate two ubiquitination sites on 14-3-3 (g) and two on ERα (h). Only peptide fragments of the native protein sequence with PSM>10 are shown. An overview of all detected peptides is presented in SI Table S1.

Ubiquitin conjugation was rapid. Higher molecular weight species were evident within 5 minutes (Fig. S11) and were quantified at 15 minutes by Coomassie-stained SDS–PAGE (Fig. 6b, c; Fig. S12). No ubiquitination was seen in the DMSO control, consistent with ubiquitination being contingent on induced-proximity. By contrast, CV2p, CV3a, and CV2a each promoted robust polyubiquitination of the 14-3-3ζ/ERα^(LBD_F)^ complex, with modification levels correlating with the degree of VCB cooperativity measured in biophysical assays. CV2a and CV3a produced the strongest effects, whereas stereochemical inversion of CV2a ((*S*–OH)-CV2a) completely abrogated ubiquitination, confirming dependence on cooperative ^MG^PROTAC–VCB engagement. As 14-3-3ζ and ERα^(LBD_F)^ migrate at similar apparent molecular weights in the SDS-PAGE, ATP-dependent and ^MG^PROTAC-dependent ubiquitination of both proteins was validated by western blot analysis and using an Alexa Fluor 488-labelled ubiquitin (Fig. 6d–f).

To define E2 requirements, we compared UBE2D2 and UBE2R1 using CV2p as a characteristic moderately cooperative ^MG^PROTAC. UBE2D2 was selected as a representative of the promiscuous UBE2D family which are known to work with Cullin RING E3 ligases such as CRL2^VHL^.^39–43^ The UBE2R family are known to be more apt at extending polyubiquitin chains^40–42^, however recent studies have shown that UBE2R1 and UBE2R2 are also capable of installing the first ubiquitin on degrader-targeted substrates with CRL2^VHL^.^44,45^ In this system, efficient ubiquitination of the 14-3-3ζ/ERα^(LBD_F)^ complex was observed only with UBE2D2 (Extended Fig. 8), whereas UBE2R1 failed to promote detectable substrate labelling under the same conditions. Furthermore, CV2p induced preferential ubiquitination of the assembled 14-3-3ζ/ERα^(LBD_F)^ complex over apo-14-3-3, with the latter showing less modification even after 24 h incubation.

### ^MG^PROTAC design controls ubiquitination sites within the protein complex

To define site selectivity, we mapped ubiquitination sites by mass spectrometry after in vitro ubiquitination and proteolytic digestion of the 14-3-3ζ/ERα^(LBD_F)^ assembly. Ubiquitination was detected at Lys139 and Lys158 of 14-3-3ζ and at Lys467 and Lys531 of ERα^(LBD_F)^; a modification at residue 253 of the 14-3-3 construct was excluded because it resides in the cleaved protease cleavage scar and is absent from the native sequence. Only peptides with PSM > 10 were considered; a full peptide list is provided in Table S1. To place these sites in structural context, we aligned VHL–EloC–EloB from our cryo-EM model with a previously reported VHL–EloC–EloB–Cul2–Rbx1–UBE2R1–Ub structure (Fig. 6g–h). In this overlay, the Cullin scaffold projected from the VHL–EloC–EloB module on 14-3-3ζ monomer 1 towards 14-3-3ζ monomer 2, orientating over the top of the 14-3-3ζ/ERα^(LBD_F)^ complex. This places the Cullin C-terminus proximal to 14-3-3ζ Lys139 and Lys158 of 14-3-3 monomer 2 (Fig. 6g, left). Complementary AlphaFold modelling of the ERα^(LBD_F)^ dimer positioned Lys467and Lys531 on the same face of the dimer toward the Cullin C-terminus, supporting a geometry in which these ERα lysines are accessible to ubiquitin transfer (Fig. 6g, right).

Comparison with our biochemical data showed that increased cooperativity correlated with ubiquitination levels and that ^MG^PROTAC linker chemistry directed site selectivity (Extended Fig. 9). CV3a biased modification towards 14-3-3ζ, whereas the most cooperative ^MG^PROTAC, CV2a, preferentially targeted ERα^(LBD_F)^. This increased labeling of ERα may arise from cooperative contacts between ERα and VHL within the VCB module (Fig. 5c); however, that region of the cryo-EM map is poorly resolved, precluding in-depth structural analysis. Together, these observations indicate that tuning linker length and composition orients the ligase–substrate assembly and can favour ubiquitination of one protein within the complex over the other.

Taken together with the cellular and ubiquitination data, these results show that cooperative ^MG^PROTACs confer selective, E2-dependent ubiquitin transfer onto the 14-3-3ζ/ERα^(LBD_F)^ complex. Ubiquitination level is enhanced by cooperative quaternary complex formation and target preference is fine-tuned by the chemical linker of the ^MG^PROTAC.

## Discussion

This study establishes a structural and mechanistic framework for chemically induced ubiquitination of targeted protein complexes. By covalently linking a molecular glue stabilizer ligand—which recruits a disordered region of the partially disordered ERα target protein to 14-3-3—to an E3 ligase ligand (VHL), we demonstrate that induced proximity can be extended beyond single, well-folded proteins. Furthermore, we show that tuning the linker length and composition allows precise control over ^MG^PROTAC cooperativity, E3 ligase recruitment, and site-selective ubiquitination efficiency. Cryo-EM analysis of the most cooperative ^MG^PROTAC, CV2a, revealed a *de novo* interface between VHL and 14-3-3ζ mediated by the α8/α9 helices of 14-3-3ζ—regions previously implicated in endogenous complexes such as BRAF/MEK1/14-3-3^46^, LRRK2/14-3-3^47^, HSPB6/14-3-3^48^, and KRAS/BRAF/MEK1/14-3-3^49^. These helices have also been reported to form secondary binding sites for small-molecule fragments.^50^ Uniquely, the interaction described here represents an unprecedented ligand-induced interface between a 14-3-3:client complex and a neo-substrate, illustrating an additional cooperativity driven by the bifunctional molecule. This cooperative engagement between 14-3-3ζ and VHL, established from our biophysical data, explains the VHL-dependent enhancement of 14-3-3/ERα stabilization observed *in vitro*. Crucially, we find that cooperative formation of stable complexes is conducive to productive and efficient protein ubiquitination at specific lysine residues, including on ERα at its globular domain far from the disordered region directly recruited to the E3 ligase by our ^MG^PROTAC. Lysine ubiquitination preference is fine-tuned depending on the chemical nature of the compound inducing proximity, demonstrating that ^MG^PROTACs can selectively ubiquitinate specific protein species within a biomolecular complex.

Despite efficient ubiquitination of both 14-3-3ζ and ERα^(LBD_F)^ in biochemical assays, no measurable reduction in ERα levels was detected in cells. A possible explanation for this lack of degradation is the limited characterized phosphorylation state of ERα at Thr594, which mediates 14-3-3 binding. Moreover, fusicoccin is known to engage multiple 14-3-3:client protein complexes in cells in addition to the one studied here.^51^ Together these constraints likely restrict cellular degradation activity and explain why total ERα levels were unchanged even with the best compounds. Future focus on highly abundant 14-3-3-interacting phosphoproteins, or protein interactions beyond 14-3-3 complexes, could enhance the probability to observe induced ubiquitination translating into degradation.

These findings expand the conceptual drug discovery scope of induced-proximity approaches by recruiting a protein complex for induced-ubiquitination through a hybrid molecular glue PROTAC strategy. By demonstrating that degrader architecture can tune both the extent and topology of induced ubiquitination through cooperative interface formation, this work provides a blueprint for designing ^MG^PROTAC capable of reprogramming the ubiquitin–proteasome system toward protein assemblies composed of dynamic and disordered regions previously considered undruggable.

## Acknowledgements

We acknowledge Bente A. Somsen and Eline Sijbesma for their foundational work developing the recombinant expression of the 14-3-3/ERα complex; Joost L.J van Dongen and Sebastian A.H. van den Wildenberg for assistance with preparative HPLC and high-resolution LC-MS measurements; Marloes Pennings and Galen Miley for support with TR-FRET assay development (Department of Biomedical Engineering, Eindhoven University of Technology). We acknowledge the technical and research staff at CeTPD for the setup and upkeep of protein purification and computational infrastructure. We thank Diane Cassidy from CeTPD for assisting in cellular experiments; Ramasubramanian Sundaramoorthy for access and support with Dundee’s School of Life Sciences in-house cryo-EM Facility where screening and preliminary data collection was performed; P. da Fonseca and E. Morris (University of Glasgow) and David Owen (Diamond Light Source) for microscopy support at eBIC where the final dataset was collected; Ronald Hay (Dundee’s School of Life Sciences) for access to mass photometry.

## Author contributions

CJAV, CO, and PJC conceived the project, with input from LB and AC. CJAV and PJC designed and planned the project. CJAV and PJC designed the compounds, which were synthesized, purified, and characterized by CJAV and NAS. CJAV, MAN, and CC expressed and purified recombinant proteins. CJAV and TH performed biophysical studies. TH performed mass photometry experiments. CC and MAN designed and conducted the in-vitro ubiquitination experiments assays as well as the cryo-EM sample preparation, imaging, data processing, and modelling. CC and KI performed molecular dynamics simulations. ADB and TH carried out cellular target engagement assays. GS performed in vitro mass spectrometry experiments and data analysis with input from MAN. CJAV, CC, AC, and PJC analyzed and interpreted the data and co-wrote the manuscript with input from all co-authors. LB, AC, and PJC supervised the project and acquired funding. All authors approved the final version of the manuscript.

## Note

MAN is presently at Birkbeck University of London, ISMB EM Facility. Malet Street, London, WC1E 7HX, UK.

## Data and materials availability

All data needed to evaluate the conclusions in the paper are present in the paper and/or the Supplementary Materials. Atomic models have been deposited to the PDB under ID 9SV3. Electron microscopy reconstructions have been deposited to the Electron Microscopy Databank (EMDB) under accession codes EMD-55233, EMD-55234, EMD-55235, EMD-55236 and EMD-55237

## Methods

### Protein expression and purification 14-3-3 proteins

N-terminally His_6_-tagged 14-3-3 proteins (sigma isoform residues 1-248, beta isoform residues 1-246, gamma isoform residues 1-247, epsilon isoform residues 1-255, zeta isoform residues 1-245, eta isoform residues 1-246, sigma isoform residues 1-248, tau isoform residues 1-245) were expressed and purified as previously described.^52^ In short, single colonies of cDNA transformed into *Escherichia coli* BL21(DE3) cells were grown overnight in lysogeny broth (LB) supplemented with ampicillin (100 µg mL^−1^) at 37°C with shaking. The overnight culture was diluted 1:100 in LB supplemented with ampicillin (100 µg mL^−1^) and grown to an optical density (OD_600_) of ∼1.2. Protein expression was induced overnight at 18°C with 0.5 mM isopropyl β-d-1-thiogalactopyranoside (IPTG). The cells were harvested by centrifugation and frozen at −80°C as pellets until further purification. The bacterial pellets were resuspended in buffer (50 mM HEPES, 300 mM NaCl, 12.5 mM imidazole, 2 mM β-mercaptoethanol, pH 8.0) supplemented with 5 mM MgCl_2_, benzonase (5 μL/ 100 mL), and 1× EDTA-free Roche protease inhibitor cocktail, and lysed by cell disruption using a C3 Emulsiflex-C3 homogenizer (Avestin) operating at 4°C. Cellular debris was removed by centrifugation. The His_6_-tagged proteins were purified on a Ni-NTA Superflow cardridge (Qiagen) and eluted with 250 mM imidazole. The proteins were dialyzed into 25 mM HEPES, 100 mM NaCl, 10 mM MgCl_2_, 0.5 mM TCEP, pH 7.5, concentrated, flash frozen with liquid nitrogen, and stored at −80°C.

N-terminally biotinylated 14-3-3 protein (gamma isoform, residues 1-247) was prepared by BirA enzyme co-expression. In short, single colonies of His-AVI-14-3-3 and BirA cDNA transformed into *Escherichia coli* BL21(DE3) cells were grown overnight in lysogeny broth (LB) supplemented with ampicillin (100 µg mL^−1^) at 37°C with shaking. The overnight culture was diluted 1:100 in LB supplemented with ampicillin (100 µg mL^−1^) and biotin (500 µM) and grown to an optical density (OD_600_) of ∼1.2. Protein expression was induced overnight at 18°C with 0.5 mM isopropyl β-d-1-thiogalactopyranoside (IPTG). The cells were harvested by centrifugation and frozen at −80°C as pellets until further purification. The bacterial pellets were resuspended in buffer (50 mM HEPES, 300 mM NaCl, 12.5 mM imidazole, 2 mM β-mercaptoethanol, pH 8.0) supplemented with 5 mM MgCl_2_, benzonase (5 μL/ 100 mL), and 1× EDTA-free Roche protease inhibitor cocktail, and lysed by cell disruption using a C3 Emulsiflex-C3 homogenizer (Avestin) operating at 4°C. Cellular debris was removed by centrifugation. The His_6_-tagged proteins were purified on a Ni-NTA Superflow cardridge (Qiagen) and eluted with 250 mM imidazole. The protein was dialyzed into a low-concentration imidazole buffer, incubated with Tobacco Etch Virus (TEV) protease, and flowed through a Ni-NTA Superflow cardridge (Qiagen). The protein sample was finally purified by size exclusion chromatography on a HiLoad 16/600 Superdex 75 pg column (Cytiva) in 25 mM HEPES, 100 mM NaCl, 10 mM MgCl_2_, 0.5 mM TCEP, pH 7.5, concentrated, flash frozen with liquid nitrogen, and stored at −80°C.

### 14-3-3ζ/ERα(LBD_F) complex

The 14-3-3ζ/ERα^(LBD_F)^ protein complex (14-3-3ζ residues 1-245, ERα residues 302-595 with F591R and P592R mutations to make the 14-3-3 binding epitope PKA responsive and S305A mutation to block phosphorylation of this residue) was expressed and purified as previously described.^38^ In short, single colonies of His_6_-SUMO-ERα, 14-3-3ζ-strep and SUMO-PKA cDNA transformed into *Escherichia coli* BL21(DE3) cells were grown overnight in lysogeny broth (LB) supplemented with ampicillin (100 µg mL^−1^), streptomycin (100 µg mL^−1^), and chloramphenicol (25 µg mL^−1^) at 37°C with shaking. The overnight culture was diluted 1:100 in Zyp-5052 medium supplemented with ampicillin (100 µg mL^−1^), streptomycin (100 µg mL^−1^), and chloramphenicol (25 µg mL^−1^) and grown to an optical density (OD_600_) of ∼2.0. Protein expression was induced overnight at 18°C with 0.5 mM isopropyl β-d-1-thiogalactopyranoside (IPTG). The cells were harvested by centrifugation and frozen at −80°C as pellets until further purification. The bacterial pellets were resuspended in buffer (20 mM Tris, 300 mM NaCl, 12.5 mM imidazole, 5% (v/v) glycerol, 2 mM β-mercaptoethanol, pH 8.0) supplemented with 5 mM MgCl_2_, benzonase (5 μL/ 100 mL), and 1× EDTA-free Roche protease inhibitor cocktail, and lysed by cell disruption using a C3 Emulsiflex-C3 homogenizer (Avestin) operating at 4°C. Cellular debris was removed by centrifugation. The His_6_-tagged protein complex was purified on a Ni-NTA Superflow cartridge (Qiagen) and eluted with 250 mM imidazole. The protein complex was dialyzed into a low-concentration imidazole buffer and incubated with SUMO hydrolase. The complex was further purified on a Strep-Tactin XT 4Flow cartridge (IBA) and eluted with 50 mM biotin. The protein complex was finally purified by size exclusion chromatography on a HiLoad 16/600 Superdex 75 pg column (Cytiva) in 20 mM Tris, 150 mM NaCl, 10 mM MgCl_2_, 0.5 mM TCEP, pH 7.5, concentrated, flash frozen with liquid nitrogen, and stored at −80°C.

### VHL/EloB/EloC complex

VCB (VHL residues 54 to 213, EloB residues 1 to 104, and EloC residues 17 to 112) was expressed and purified as previously described.^22^ In short, single colonies of cDNA transformed into *Escherichia coli* BL21(DE3) cells were grown overnight in lysogeny broth (LB) supplemented with ampicillin (100 µg mL^−1^) and streptomycin (50 µg mL^−1^) at 37°C with shaking. The overnight culture was diluted 1:100 in LB supplemented with ampicillin (100 µg mL^−1^) and streptomycin (50 µg mL^−1^) and grown to an optical density (OD_600_) of ∼0.6. Protein expression was induced overnight at 18°C with 0.5 mM isopropyl β-d-1-thiogalactopyranoside (IPTG). The cells were harvested by centrifugation and frozen at −80°C as pellets until further purification. The bacterial pellets were resuspended in buffer supplemented with 5 mM MgCl_2_, DNase I (1 µg/mL), and 1× EDTA-free Roche protease inhibitor cocktail, and lysed by cell disruption using One Shot Cell Disruptor (Constant Systems) operating at 4°C. Cellular debris was removed by centrifugation. The His_6_-tagged protein complex was purified on a HisTrap FF Ni NTA affinity column (Cytiva) and eluted with an imidazole concentration gradient. The protein was dialyzed into a low-concentration imidazole buffer, incubated with Tobacco Etch Virus (TEV) protease, and flowed through a HisTrap FF Ni NTA affinity column. The His₆-tagged VCB protein complex used in the TR-FRET assays was directly used in the next step without TEV cleavage or Ni-NTA purification. VCB was additionally purified by anion exchange using a HiTrap Q HP column. The protein sample was finally purified by size exclusion chromatography on a HiLoad 16/600 Superdex 75 pg column (Cytiva) in 20 mM HEPES, pH 7.5, 150 mM NaCl, and 1 mM DTT, concentrated, flash frozen with liquid nitrogen, and stored at −80°C.

### APPBP1-UBA3 and UBE2M

The APPBP1-UBA3 and UBE2M proteins were expressed and purified as previously described.^45^ In short, single colonies of cDNA transformed into *Escherichia coli* BL21(DE3) cells were grown overnight in lysogeny broth (LB) supplemented with ampicillin (100 µg mL^−1^) at 37°C with shaking. The overnight culture was diluted 1:100 in LB supplemented with ampicillin (100 µg mL^−1^) and grown to an optical density (OD_600_) of ∼0.6. Protein expression was induced overnight at 16°C with 0.5 mM isopropyl β-d-1-thiogalactopyranoside (IPTG). The cells were harvested by centrifugation and frozen at −80°C as pellets until further purification. The bacterial pellets were resuspended in 50 mM Tris, 200 mM NaCl, and 5 mM DTT, pH 7.5, supplemented with 5 mM MgCl_2_, DNase I (1 µg/mL), and 1× EDTA-free Roche protease inhibitor cocktail, and lysed by cell disruption using One Shot Cell Disruptor (Constant Systems) operating at 4°C. Cellular debris was removed by centrifugation. The lysate was incubated with 10 mL of glutathione Sepharose 4B (Cytiva) resin for 1 hour at room temperature. The resin was washed with buffer and the protein was eluted with 10 mM reduced L-glutathione. One unit of thrombin per milligram of total protein was added and the mixture was incubated overnight at −4°C. APPBP1-UBA3 was further purified by size exclusion chromatography on a HiLoad 16/600 Superdex 200 pg column (Cytiva) in 20 mM HEPES, pH 7.5, 150 mM NaCl, and 0.5 mM TCEP. UBE2M was further purified by size exclusion chromatography on a HiLoad 16/600 Superdex 75 pg column (Cytiva) in 20 mM HEPES, pH 7.5, 150 mM NaCl, and 0.5 mM TCEP. The resulting proteins were flowed over glutathione Sepharose 4B resin to remove any trace amounts of GST-APPBP1-UBA3, GST-UBE2M, and GST. The purified protein samples were each concentrated, flash frozen with liquid nitrogen, and stored at −80°C.

### NEDD8 (residues 1 to 76)

The NEDD8 protein (residues 1-76) was expressed and purified as previously described.^45^ In short, single colonies of cDNA transformed into *Escherichia coli* BL21(DE3) cells were grown overnight in LB supplemented with ampicillin (100 μg mL^−1^) at 37°C with shaking. The overnight culture was diluted 1:100 in LB supplemented with ampicillin (100 μg mL^−1^) and grown to an optical density (OD_600_) of ∼0.6. Protein expression was induced overnight at 16°C with 0.5 mM isopropyl β-d-1-thiogalactopyranoside (IPTG). The cells were harvested by centrifugation and frozen at −80°C as pellets until further purification. The bacterial pellets were resuspended in 30 mM Tris, 200 mM NaCl, and 5 mM dithiothreitol (DTT), pH 7.5, supplemented with 5 mM MgCl_2_, DNase I (1 μg/mL), and 1× EDTA-free Roche protease inhibitor cocktail, and lysed by cell disruption using One Shot Cell Disruptor (Constant Systems) operating at 4°C. Cellular debris was removed by centrifugation. The His_6_-tagged protein was purified on a HisTrap FF Ni NTA affinity column (Cytiva) and eluted with an imidazole concentration gradient (20 to 500 mM). The resulting eluate was incubated with 10 mL of glutathione Sepharose 4B resin (Cytiva). The protein-bound resin was washed with buffer, Tobacco Etch Virus (TEV) protease was added to the resin, and NEDD8 was cleaved at room temperature for 3 hours. NEDD8 was finally purified by size exclusion chromatography on a HiLoad 16/600 Superdex 75 pg column (Cytiva) in 20 mM HEPES, pH 7.5, 150 mM NaCl, and 0.5 mM TCEP, concentrated, flash frozen with liquid nitrogen, and stored at −80°C.

### NEDD8-CRL2^VHL^

The NEDD8-CRL2^VHL^ protein complex was expressed and purified as previously described.^45^ In short, single colonies of Cul2-Rbx1 cDNA^53^ transformed into *Escherichia coli* BL21(DE3) cells were grown overnight in LB supplemented with ampicillin (100 μg mL^−1^) at 37°C with shaking. The overnight culture was diluted 1:100 in LB supplemented with ampicillin (100 μg mL^−1^) and grown to an optical density (OD_600_) of ∼0.6. Protein expression was induced for 16 hours at 16°C with 0.2 mM isopropyl β-d-1-thiogalactopyranoside (IPTG). The cells were harvested by centrifugation and frozen at −80°C as pellets until further purification. The bacterial pellets were resuspended in 30 mM Tris, 200 mM NaCl, and 5 mM DTT, pH 7.5, supplemented with 5 mM MgCl_2_, DNase I (1 μg/mL), and 1× EDTA-free Roche protease inhibitor cocktail, and lysed by cell disruption using One Shot Cell Disruptor (Constant Systems) operating at 4°C. Cellular debris was removed by centrifugation. Cul2-Rbx1 was purified on a HisTrap FF Ni NTA affinity column (Cytiva) and eluted with an imidazole concentration gradient from 0 to 300 mM. The protein was desalted into 20 mM HEPES, 150 mM NaCl, 0.5 mM TCEP, and 5% (v/v) glycerol, pH 7.5, with a HiPrep 26/10 desalting column (Cytiva) and incubated with Tobacco Etch Virus (TEV) protease and an excess of recombinant VHL-EloB-EloC. VHL-EloB-EloC-Cul2-Rbx1 (CRL2^VHL^) was purified on a StrepTrap XT column (IBA), and eluted with 50 mM biotin. Purified CRL2^VHL^ (1.7 μM) was incubated with APPBP1-UBA3 (250 nM), Ube2M (1.2 μM), NEDD8 (20 μM), adenosine triphosphate (ATP) (1 mM), and MgCl_2_ (5 mM) for 10 min at 37°C. NEDD8-CRL2^VHL^ was finally purified by size exclusion chromatography on a HiLoad 16/600 Superdex 200 pg column (Cytiva) in 20 mM HEPES, 150 mM NaCl, 0.5 mM TCEP, and 5% (v/v) glycerol, pH 7.5, concentrated, flash frozen with liquid nitrogen, and stored at −80°C.

### GACG-ubiquitin

The GACG-ubiquitin protein was expressed and purified as previously described.^45^ In short, single colonies of His_6_-TEV-GACG-ubiquitin cDNA^54^ transformed into SoluBL21(DE3) competent cells were grown overnight in LB supplemented kanamycin (50 μg mL^−1^) at 37°C with shaking. The overnight culture was diluted 1:100 in LB supplemented with kanamycin (50 μg mL^−1^) and grown to an optical density (OD_600_) of ∼0.75. Protein expression was induced for 4 hours at 37°C with 1 mM IPTG. The cells were harvested by centrifugation and frozen at −80°C as pellets until further purification. The bacterial pellets were resuspended in 20 mM HEPES, 500 mM NaCl, and 0.5 mM TCEP, pH 7.5, supplemented with 5 mM MgCl_2_, DNase I (1 μg/mL), and 1X EDTA-free Roche protease inhibitor cocktail, and lysed by cell disruption using One Shot Cell Disruptor (Constant Systems) operating at 4°C. Cellular debris was removed by centrifugation. His_6_-TEV-GACG-ubiquitin was purified on a HisTrap FF Ni NTA affinity column (Cytiva) and eluted with an imidazole concentration gradient from 0 to 500 mM. The protein was dialyzed into 50 mM HEPES, 150 mM NaCl, and 0.5 mM TCEP, pH 7.5, incubated with Tobacco Etch Virus (TEV) protease, and flowed through a HisTrap FF Ni NTA affinity column. GACG-ubiquitin was finally purified by size exclusion chromatography on a HiLoad 16/600 Superdex 75 pg column (Cytiva) in 50 mM Tris, 150 mM NaCl, and 0.5 mM TCEP, pH 7.5, concentrated, flash frozen with liquid nitrogen, and stored at −80°C.

### Ubiquitin-activating enzyme 1

The ubiquitin-activating enzyme 1 (UBA1) was expressed and purified as previously described.^45^ In short, single colonies of UBA1-His_6_ cDNA^43^ transformed into *Escherichia coli* Rosetta 2(DE3) competent cells were grown overnight in LB supplemented with kanamycin (50 μg mL^−1^) at 37°C with shaking. The overnight culture was diluted 1:100 in LB supplemented with kanamycin (50 μg mL^−1^) and 1 mM MgSO_4_ and grown to an optical density (OD_600_) of ∼0.8. Protein expression was induced for 16 hours at 16°C with 0.2 mM isopropyl β-d-1-thiogalactopyranoside (IPTG). The cells were harvested by centrifugation and frozen at −80°C as pellets until further purification. The bacterial pellets were resuspended in IMAC buffer A (20 mM sodium phosphate, 500 mM NaCl, 50 mM imidazole, and 1 mM TCEP, pH 7.4), supplemented with 5 mM MgCl_2_ DNase I (1 μg/mL), and 1X EDTA-free Roche protease inhibitor cocktail, and lysed by cell disruption using One Shot Cell Disruptor (Constant Systems) operating at 4°C. Cellular debris was removed by centrifugation, and UBA1 was purified on a HisTrap FF Ni NTA affinity column (Cytiva) and eluted with 100% IMAC Buffer B (20 mM sodium phosphate, 500 mM NaCl, 350 mM imidazole, and 1 mM TCEP, pH 7.4). The elution was concentrated to ∼2 mL and diluted 1:100 in Anion Buffer A (50 mM Tris and 5 mM β-mercaptoethanol, pH 8.0) and loaded on an equilibrated Q-HP column (Cytiva). A 50% gradient over 40 CV with Anion Buffer B (50 mM Tris, 1 M NaCl, and 5 mM β-mercaptoethanol, pH 8.0) was applied and UBA1 containing fractions were concentrated and finally purified by size exclusion chromatography on a 16/600 Superdex 200 column (Cytiva) equilibrated in 20 mM HEPES, 150 mM NaCl, and 1 mM TCEP, pH 8.0, concentrated, flash frozen with liquid nitrogen, and stored at −80°C.

### UBE2D2 and UBE2R1

The UBE2D2 and UBE2R1 proteins were expressed and purified as previously described.^45^ In short, single colonies of cDNA encoding for the His_6_-TEV-E2 constructs transformed into *Escherichia coli* BL21(DE3) cells were grown overnight in lysogeny broth (LB) supplemented with kanamycin (50 µg mL^−1^) at 37°C with shaking. The overnight culture was diluted 1:100 in LB supplemented with kanamycin (100 µg mL^−1^) and grown to an optical density (OD_600_) of ∼0.6. Protein expression was induced overnight at 18°C with 0.3 mM isopropyl β-d-1-thiogalactopyranoside (IPTG). The cells were harvested by centrifugation and frozen at −80°C as pellets until further purification. The bacterial pellets were resuspended in 20 mM HEPES, 500 mM NaCl, and 0.5 mM TCEP, pH 7.5, supplemented with 5 mM MgCl_2_, DNase I (1 μg/mL), and 1X EDTA-free Roche protease inhibitor cocktail, and lysed by cell disruption using One Shot Cell Disruptor (Constant Systems) operating at 4°C. Cellular debris was removed by centrifugation. The His_6_-tagged protein was purified on a HisTrap FF Ni NTA affinity column (Cytiva) and eluted with an imidazole concentration gradient of 0 to 500 mM. The protein was dialysed into a low concentration imidazole buffer, incubated with Tobacco Etch Virus (TEV) protease, and flowed through a HisTrap FF Ni NTA affinity column. The proteins were finally purified by size exclusion chromatography on a HiLoad^TM^ 16/600 Superdex^TM^ 75 pg column (Cytiva) in 20 mM HEPES, pH 7.5, 150 mM NaCl and 0.5 mM TCEP, concentrated, flash frozen with liquid nitrogen and stored at −80 °C.

### Ubiquitin labelling with maleimide Alexa Fluor 488

Alexa Fluor 488 C5 maleimide dye was dissolved to 5 mM in anhydrous DMSO. Ubiquitin cysteine mutant recombinant protein (GACG-ubiquitin) was buffer exchanged into 20 mM HEPES and 150 mM NaCl, pH 7.0, using a 10/300 GL Superdex 75 Increase prepacked column (Cytiva). The fractions containing the protein were combined, concentrated to 100 µM and labelled with a five times molar excess of fluorescent dye for 2 hours at room temperature. The excess dye was removed by gel filtration on a 10/300 GL Superdex 75 Increase prepacked column (Cytiva), eluting in 20 mM HEPES, 150 mM NaCl, and 0.5 mM TCEP, pH 7.5. The dye and subsequently dye-labelled protein were protected from light throughout, flash frozen, and stored at −80°C.

### Fluorescence anisotropy assays

FA assays were conducted using FAM labeled ERα 8mer peptide (10 nM, FAM-AEGFPA phosphoT V-COOH, Genscript)^55^, or FAM labeled HIF1α 10mer peptide (25 nM, FAM-DEALA hydroxyPYIPD-COOH, Tocris).^56^ All assays were conducted using freshly prepared buffer (20 mM HEPES, 100 mM NaCl, 100 µM TCEP, 0.1 % (v/v) Tween20, 1 mg/mL BSA, pH 7.5, 1 % (v/v) DMSO) in 384-well plates (black, round bottom, low binding surface, 10 μL sample per well, Corning). Titrations were performed using a Thermo Scientific E1-ClipTip autopipette (1:1 dilution series). Measurements were performed directly after plate preparation, using a Tecan Spark plate reader (25 °C; λ_ex_ 485 ± 20 nm; λ_em_ 535 ± 25 nm; dichroic 510 mirror; 30 flashes; 40 µs integration time; 0 µs lag time, 1 ms settle time; gain 66 for ERα peptide and 55 for HIF1α peptide; Z-position calculated from well). Wells containing solely labeled peptide in buffer were used to calculate the G-factor. The data was plotted using Origin software. Using the same software, sigmoidal functions were fitted to the data using the following formula: anisotropy = start + (end - start)·(x^n^) / (k^n^ + x^n^); start = bottom asymptote; end = top asymptote; x = titrant concentration; k = K_D_; n = Hill coefficient.

### Cellular VHL engagement assays

VHL NanoBRET target engagement assays (N2931, Promega) were conducted following the manufacturer’s guidelines. The day before transfection, MCF7 cells were seeded in a 6-well plate at 4 x 10^5 cells per well. Transfection complexes were prepared by mixing 100 μL of Opti-MEM (Thermo Fisher Scientific #11058021), 6 µL of FuGENE® HD (Promega #E2311), 1 µg Transfection Carrier DNA, and 1 µg of the VHL-NanoLuc fusion vector. This was incubated for 20 minutes at room temperature, then added to the cells. The following day, 75 µL of transfected cells (at 2.2 x 10^5 cells/mL) were plated in White, flat-bottom, Non-Binding Surface 96-well plates (Corning #3600). PROTACs and vehicle controls were prepared at 10x their final concentrations in Opti-MEM. For the live-cell assay, 10 µL of Opti-MEM was added to each well, followed by 10 µL of each 10x PROTAC and vehicle control dilution. A 100x solution of NanoBRET tracer (50 µM for VHL live-cell assay, from kit N2931, Promega) was prepared in 100% DMSO and then diluted to a 20x solution using tracer dilution buffer. A final volume of 5 µL of the tracer was added to each well for a final concentration of 0.5 µM. Plates were mixed on an orbital shaker at 300 RPM for 15 seconds and then incubated at 37 °C with 5% CO₂ for 4 hours. Following incubation, 50 µL of a substrate solution (1/400 NanoBRET Nano-Glo substrate, 1/500 Extracellular NanoLuc inhibitor in Opti-MEM) was added to each well. After 2-3 minutes of incubation at room temperature, donor (450 nm) and acceptor (610 nm) emissions were measured on a BMG Labtech PHERAstar luminescence plate reader. For the permeabilized-cell assay, 10 µL of digitonin was added instead of Opti-MEM. In each well, 10 µL of the 10x PROTAC and vehicle control dilutions were dispensed. A 100x NanoBRET tracer solution (25 µM for the VHL permeabilized-cell assay, from kit N2931, Promega) was prepared with 100% DMSO and then diluted to a 20x solution with tracer dilution buffer. A 5 µL volume of tracer was dispensed into each well at a final concentration of 0.25 µM. This was incubated for 10 minutes, and the plate was mixed at 300 RPM for 15 seconds. Finally, 50 µL of a substrate solution (1/400 NanoBRET Nano-Glo substrate in Opti-MEM) was added, and plates were incubated for 2-3 minutes at room temperature before reading on a BMG Labtech PHERAstar as described above.

### Mass photometry

Interferometric scattering microscopy was carried out using the oneMP (Refeyn). The buffer (20 mM HEPES, 150 mM NaCl, and 0.5 mM TCEP, pH 8.0) was filtered through a 0.22-μm syringe filter, and gasket wells (Grace Bio-labs CW-50R-1.0) along with high-precision 24 × 50 mm coverslips (Marienfeld) were prepared. Buffer (10 μL) was used for focusing in regular mode (128 × 34 binned pixels 18.0 μm^2^ detection area) and all data were recorded for 60 s using AcquireMP software (Refeyn). Calibration standards for mass calibration consisting of conalbumin (Mr = 75 000), aldolase (Mr = 158 000), ferritin (Mr = 440 000), and thyroglobulin (Mr = 669 000) were prepared from the gel filtration HMW calibration kit (Cytiva). All data were processed and analyzed in DiscoverMP (Refeyn).

### Cryo-EM

#### Sample preparation

The purified 1:1 complex of 14-3-3ζ/ERα^(LBD_F)^ was incubated with VCB and CV2a at a molar ratio of 14-3-3ζ:ERα^(LBD_F)^:VCB:CV2a = 1:1:1.2:1.2 on ice for 1 hour in cryo-EM buffer (20 mM HEPES, 100 mM NaCl, 0.5 mM TCEP, pH 7.5) at a concentration of 50 μM. Right before plunge freezing, the complex was diluted 20-fold in cryo-EM buffer and 3.5 μL was applied to the carbon side of 400 mesh copper (Cu) R1.2/1.3 holey carbon Quantifoil grids. TEM grids were glow discharged for 60 sec under vacuum at 30 mA using a Quorum SC7620 just before use. A Vitrobot Mark IV (Thermo Fisher Scientific) operating at 4°C and 100% humidity was set to blot force = 4 and blot time = 4 sec for plunging grids into liquid ethane.

#### Data acquisition

All grids were clipped and screened at the University of Dundee Electron Microscopy Facility using the Glacios (Thermo Fisher Scientific) operating at 200 kV equipped with a Falcon4i direct electron detector. The dataset for the 14-3-3ζ:ERα^(LBD_F)^:VCB:CV2a complex was collected at eBIC (Diamond Light Source, UK) under BAG access BI-31827-16 using Krios-4 (m07) operating a Schottky X-FEG at 300 kV and Ametek-Gatan BioQuantum K3 Imaging Filter with slit width of 20 eV. Data were recorded using single particle EPU v2.1 (Thermo Fisher Scientific) at a nominal magnification of 105,000x, calibrated pixel size of 0.829 Å, C2 aperture 70 μm, and objective aperture of 100 μm. All movies were collected using a Gatan K3 direct electron detector (5,760×4092 pixels) in counting mode over 40 dose fractions, recorded as Tiff LZW non-gain normalised, with total measured dose of 32.59 e⁻/Å², exposure time of 1.75 sec, and dose rate of = 15.7e/px/s. A total of 14,091 movies were acquired using aberration-free image shift (AFIS). The data acquisition parameters are summarised in **Table S2**.

#### Image processing, validation and model building

Image processing pipelines are described in **Extended Data Fig. 5**. Cryo-EM movies were imported into CryoSPARC v.4.5^57^ for patch motion correction, patch CTF estimation and manual curation. Particle picking was performed with crYOLO^58^ using a general model for low-pass filtered images. Particles were extracted with a 300 pixel box size and 4 rounds of 2D classification yielded a particle stack of 745,250 particles. These particles were separated into 8 classes using ab initio classification. Particles corresponding to Pose 1 were used for TOPAZ training, the resulting particles were 2D classified and the best particles were taken forward for non-uniform refinement, resulting in the 4.6 Å Pose 1 structure. Local refinement using a mask around EloB/EloC/VHL/CV2a/14-3-3ζ(monomer) resulted in the 4.3 Å locally refined Pose 1 structure. DeepEMhancer^59^ was used for Pose 1 figure generation and model building. Particles corresponding to Pose 1 were used for TOPAZ^57^ training, the resulting particles were 2D classified and the best particles were taken forward for ab initio classification into 3 classes. The full particle stack was taken forwards and refined against the two best volumes from ab initio classification, resulting in Pose 2.1 and Pose 2.2 structures. The particles corresponding to Pose 3 were refined using non-uniform refinement resulting in the Pose 3 structure. All classification was achieved by 2D classification, and 3D reconstruction was performed with *ab initio* reconstruction, heterogeneous refinement and non-uniform refinement in cryoSPARC.^57^ Representative 2D classes, orientation diagnostics, local resolution estimations and gold-standard Fourier shell correlation (GSFSC) curves were generated with cryoSPARC and are shown in **Extended Data Fig. 6**, and **Supplementary Figures S5-6**. Model building was achieved using atomic models from AlphaFold and PDB entries 4W9H, 8C43 and 8BZH, which were docked into the cryo-EM maps using rigid-body fitting with UCSF ChimeraX.^60–62^ The models were refined using ISOLDE^63^ and Phenix^64^ until reasonable agreement between the model and data were achieved. Structural figures were generated using the UCSF ChimeraX programme.^60–62^ The data image analysis, atomic modelling, refinement and validation statistics are summarised in **Tables S3-S4**.

#### Molecular dynamics simulations

Modelling of CV2a into the cryo-EM structure was performed using Maestro (Schrödinger). The docked structures of VH032 in complex with VHL and of fusicoccin A in complex with 14-3-3 were used for molecular dynamics simulations. Modifications to the chemical structures and modelling of the aliphatic linker region were performed using the build tool in Maestro (Schrödinger). The resulting ternary VHL/CV2a/14-3-3 complex was bounded with a predefined TIP3P water model in an orthorhombic box. The pH of the system was set to 7.5 and the charge of the system was neutralised by adding Na+ and Cl-atoms at 100 mM. The temperature and the pressure were kept constant throughout the simulation. The system was energy minimised using Desmond^65^, and then subjected to a 500 ns MD simulation using Desmond^65^. The three-dimensional structures and trajectories were inspected using Maestro (Schrödinger).

#### Time-resolved FRET assays

TR-FRET assays were conducted using His-tagged VCB, biotinylated 14-3-3γ, ERα 8mer peptide (Ac-AEGFPA phosphoT V-COOH, Genscript)^55^, anti-His mAb Tb-conjugate (61HISTLF, Revvity), and streptavidin D2-conjugate (610SADLF, Revvity). All assays were conducted using freshly prepared buffer (10 mM HEPES, 150 mM NaCl, 5 mM DTT, 100 μM CHAPS, 1 mg/mL BSA, pH 7.5, 1 % (v/v) DMSO) in 384-well plates (white, round bottom, low binding surface, 5 μL sample per well, Corning). Titrations were performed using a Thermo Scientific E1-ClipTip autopipette (1:1 dilution series). Measurements were performed directly after plate preparation, using a Tecan Spark plate reader (25 °C; λ_ex_ 340 ± 35 nm; λ_em_ 620 ± 10 nm (gain 159) and 665 ± 8 nm (gain 178); dichroic 510 mirror; 30 flashes; 500 µs integration time; 150 µs lag time, 1 ms settle time; Z-position calculated from well). The TR-FRET ratio was calculated by dividing the emission signal at 665 nm by the emission signal at 620 nm of the respective well. The data was plotted using Origin software and fitted using a freehand fit to approximate the trend, with no underlying model applied. All datapoints are based on three independent experiments from which the average and standard deviation were calculated.

#### In vitro ubiquitination assays

For the time-course ubiquitination assay and mass spectrometry analysis, the E1 Ube1 (2 μM), the E2 UBE2D2 (5 μM), the E2 UBE2R1 (1 μM), 14-3-3zeta/ERa^LBD_F^ (12 μM), CRL2^VHL^ (500 nM), and PROTAC (5 μM) were mixed with ubiquitin (300 μM) and incubated in 30 mM Tris, 150 mM NaCl, 5% glycerol, and 15 mM MgCl_2_, pH 8.0, for 15 min at room temperature. ATP (10 mM) was added, and the mixture was incubated at 37 °C. The reaction was quenched with SDS sample buffer. The protein species were either taken forwards for mass spectrometry analysis or resolved by SDS-PAGE on a 4-12% bis-tris NuPAGE gel using MES running buffer, run at 200 V for 35 min. The SDS-PAGE gel was stained with Coomassie Instant Blue.

For all other ubiquitination assays, the E1 Ube1 (150 nM), the E2 UBE2D2 or UBE2R1 (5 μM), 14-3-3zeta/ERa^LBD_F^ (5 μM), NEDD8-CRL2^VHL^ (150 nM), and PROTAC (1 μM) were mixed with ubiquitin (98 μM) and Alexa Fluor 488-labelled ubiquitin (2 μM) and incubated in 20 mM HEPES, 150 mM NaCl, 0.5 mM TCEP, and 5 mM MgCl_2_, pH 7.5, for 5 min at room temperature. ATP (3 mM) was added, and the mixture was incubated at room temperature. The reaction was quenched with reducing SDS sample buffer. The protein species were resolved by SDS-PAGE on a 10% bis-tris NuPAGE gel using MOPS running buffer, run at 175 V for 50 min. The SDS-PAGE gel was stained with Coomassie Instant Blue, or taken forward for Western Blot. Proteins were transferred to a nitrocellulose membrane either by wet transfer at 90 V for 90 minutes, or using an iBlot 2 Gel Transfer Device (Invitrogen). Membranes were blocked for 1 hour in 5% milk in TBS-T and incubated overnight with primary antibodies (1:1000, for ERα rabbit mAb CST #8644, for 14-3-3 CST #8312), washed with TBS-T, then incubated for 1 hour with secondary antibodies (1:5000) before imaging on the ChemiDoc (Biorad).

#### Mass spectrometry to identify ubiquitination sites

The in vitro ubiquitination assay samples were used for mass spectrometry analysis. Reduction and alkylation of cysteine residues were carried out using a final concentration of 10 mM dithiothreitol (DTT) at 60 °C for 20 min; and 20 mM iodoacetamide (IAA) at room temperature for 30 min in the dark, respectively. In-solution digestion was carried out with 50 µg of protein from each sample. Digestion of the samples was carried out using trypsin (sequencing grade) in the ratio of 1:20 (enzyme:protein) at 37 °C overnight at 500 rpm in a thermomixer. Samples were then acidified with an aqueous solution of 1% formic acid to stop the reaction and were desalted using Sep-Pak C_18_ tip. Eluted samples were dried in a speed vacuum. The dried samples were reconstituted in 1% formic acid and analyzed on an Orbitrap Ascend mass spectrometer coupled to a Thermo Fisher Scientific Vanquish Neo UHPLC. The peptides were enriched on a trap column and resolved on an analytical column (Easy-Spray PepMap Neo 2 μm C18 75 μm × 150 mm) with 300 nl/min. The gradient for separation was used as 35% B at 70 min. Total run time used was 100 min. The MS data acquisition was carried out from 350– 1600 m/z range using an Orbitrap mass analyzer. The automatic gain control (AGC) target was set to 400,000 with an ion injection time of auto and a dynamic exclusion of 45 s. The orbitrap Ascend mass spectrometer was operated in the data-dependent acquisition (DDA) mode. Precursor ions were fragmented using 27 high collision dissociation (HCD) and were analyzed using an ion trap mass analyzer. The mass spectrometry raw data were searched using SEQUEST HT search engines with Proteome Discoverer 3.0 (Thermo Fisher Scientific). The following parameters were used for searches: Precursor mass tolerance 10 ppm, fragment mass tolerance 0.6, enzyme: trypsin, Mis-cleavage: −2, fixed modification: carbamidomethylation of cysteine residues and dynamic modification: oxidation of methionine, DiGly of lysine. The data were filtered for 1% PSM, peptide and protein level FDR. For quantitation label free workflow based on precursor area was used.

## Extended data Figures

**Extended Data Fig. 1.**
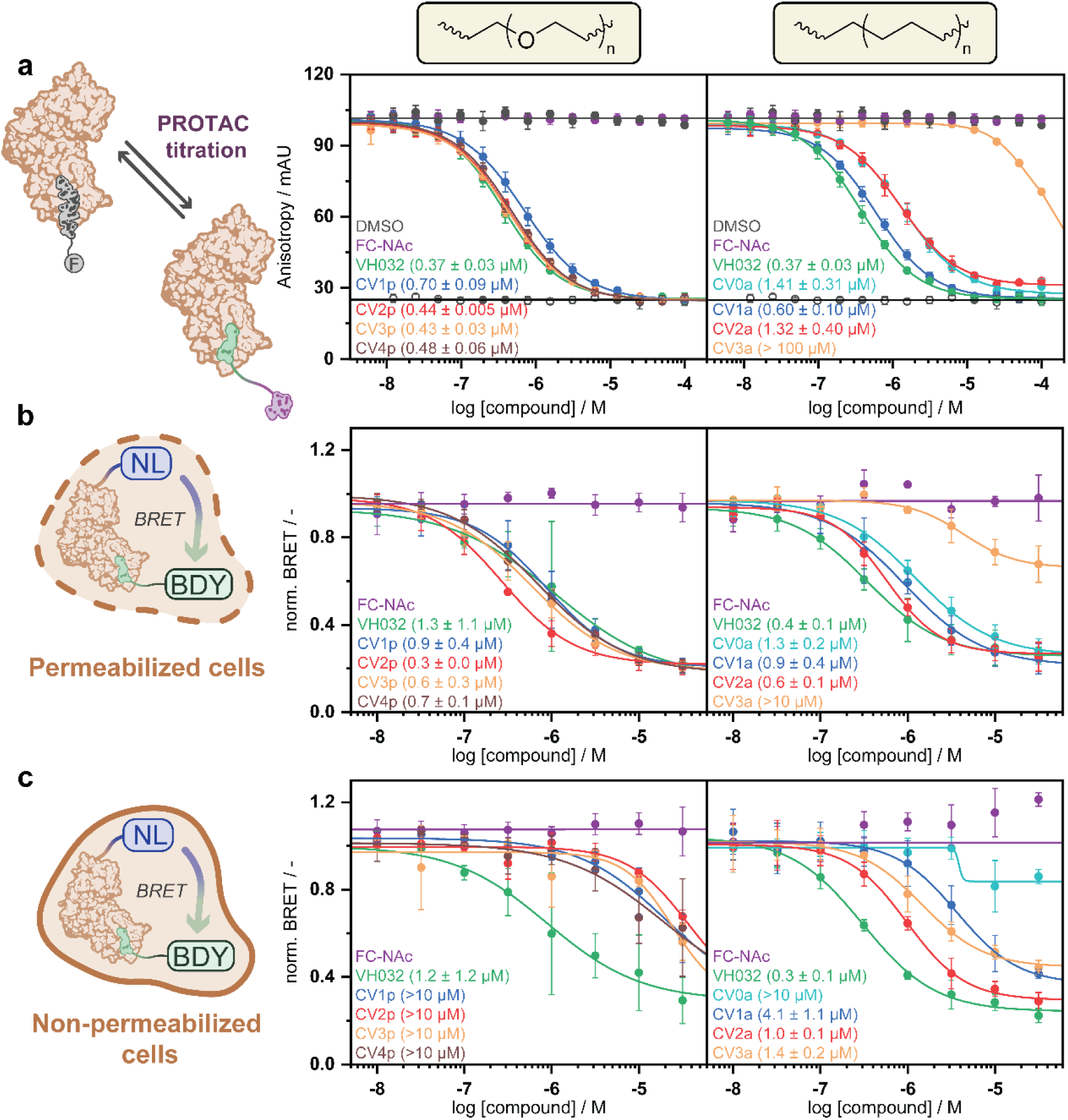
Biophysical and in cell VHL engagement. (a) FA displacement assay titrating monovalent control compound, PEG linker-based ^MG^PROTAC, or alkyl linker-based ^MG^PROTAC to 50 nM VCB and 25 nM fluorescently labeled HIF1α peptide. (b) MCF7 cells were transiently transfected with NanoLuc-VHL to from a BRET pair with the VHL tracer (0.25 µM, Promega). Normalized BRET signal of permeabilized cells after 10 min treatment with the ^MG^PROTAC. (c) Normalized BRET signal of non-permeabilized cells after 4 h treatment with the ^MG^PROTAC.

**Extended Data Fig. 2.**
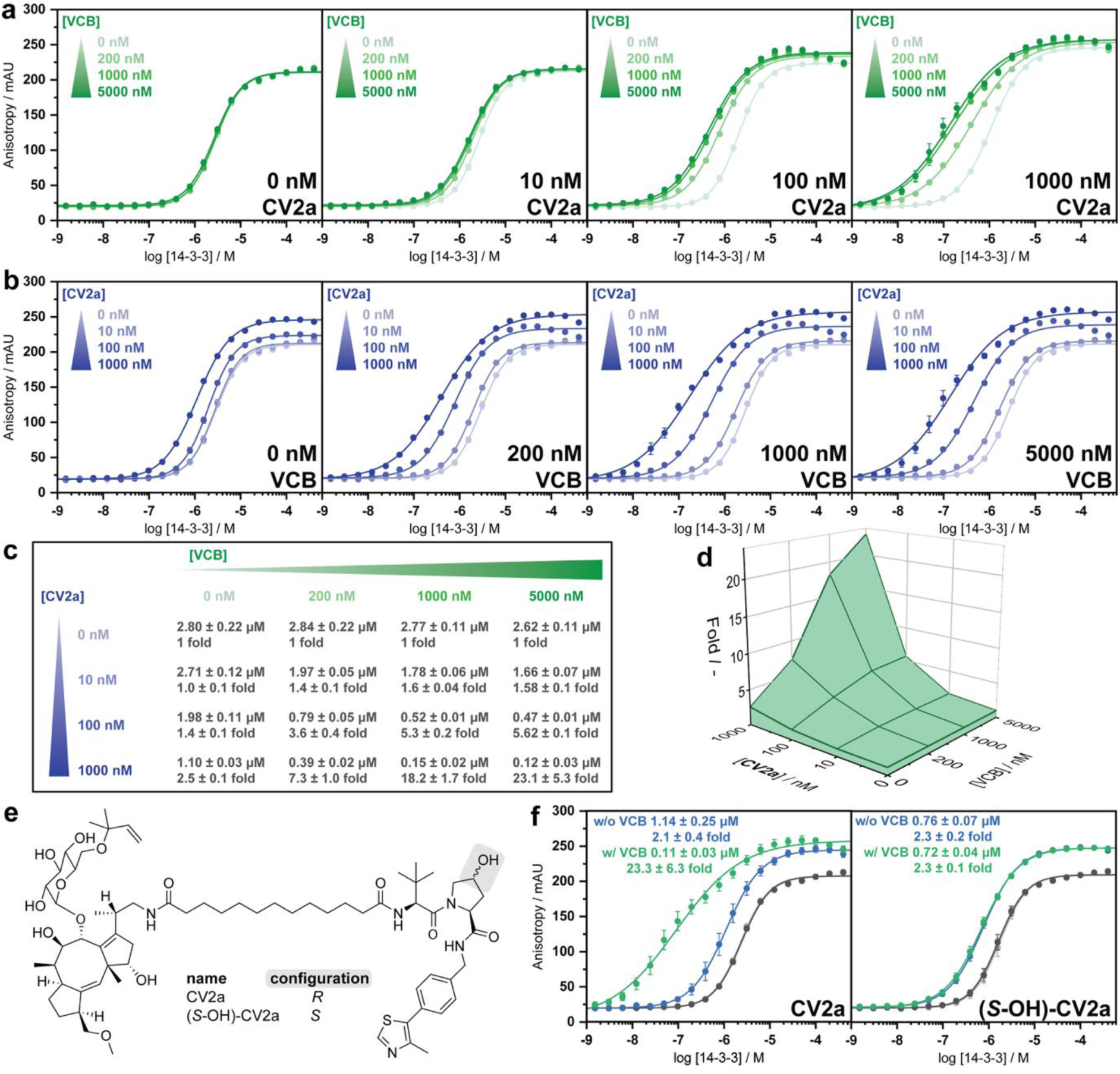
ERα protein stabilization is driven by cooperative protein interactions between VHL and the 14-3-3/ERα complex. FA 14-3-3 titrations in the presence of various CV2a concentrations (0, 10, 100, 1000 nM) and various VCB concentrations (0, 200, 1000, 5000 nM), and 10 nM fluorescently labeled ERα peptide. (a) Titration data grouped per CV2a concentration, and (b) per VCB concentration. (c) Overview of the fitted KDapp values and fold left shift relative to the 14-3-3 titration w/o CV2a and a similar VCB concentration. (d) 3-dimensional plot of the obtained fold left shifts. (e) Chemical structure of CV2a and (*S*-OH)-CV2a. (f) FA 14-3-3 titrations in the presence of 1 µM CV2a or its diastereoisomer (S-OH)-CV2a, and 10 nM fluorescently labeled ERα peptide in the absence of VCB (w/o VCB, blue dots) and presence of 5 µM VCB (w/ VCB, green dots). The DMSO control is indicated in grey (w/o VCB light grey dots, w/ VCB dark grey dots, data overlaps).

**Extended Data Fig. 3.**
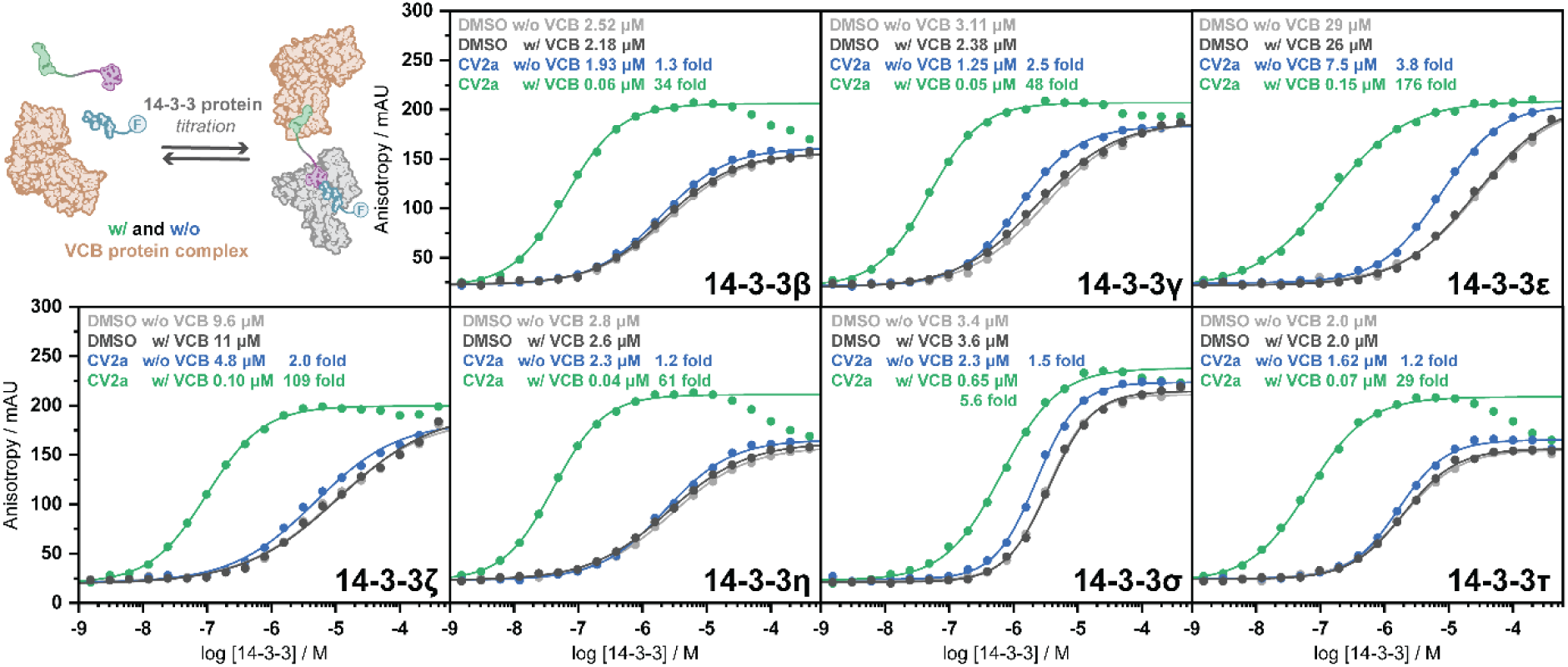
VHL selective recruitment to different 14-3-3 isoforms. FA titration of all 7 human 14-3-3 isoforms in the presence of 500 nM monovalent compound or ^MG^PROTAC, and 10 nM fluorescently labeled ERα peptide in the absence of VCB (w/o VCB, blue dots) and presence of 1 µM VCB (w/ VCB, green dots). The DMSO control is indicated in grey (w/o VCB light grey dots, w/ VCB dark grey dots, data overlaps).

**Extended Data Fig. 4.**
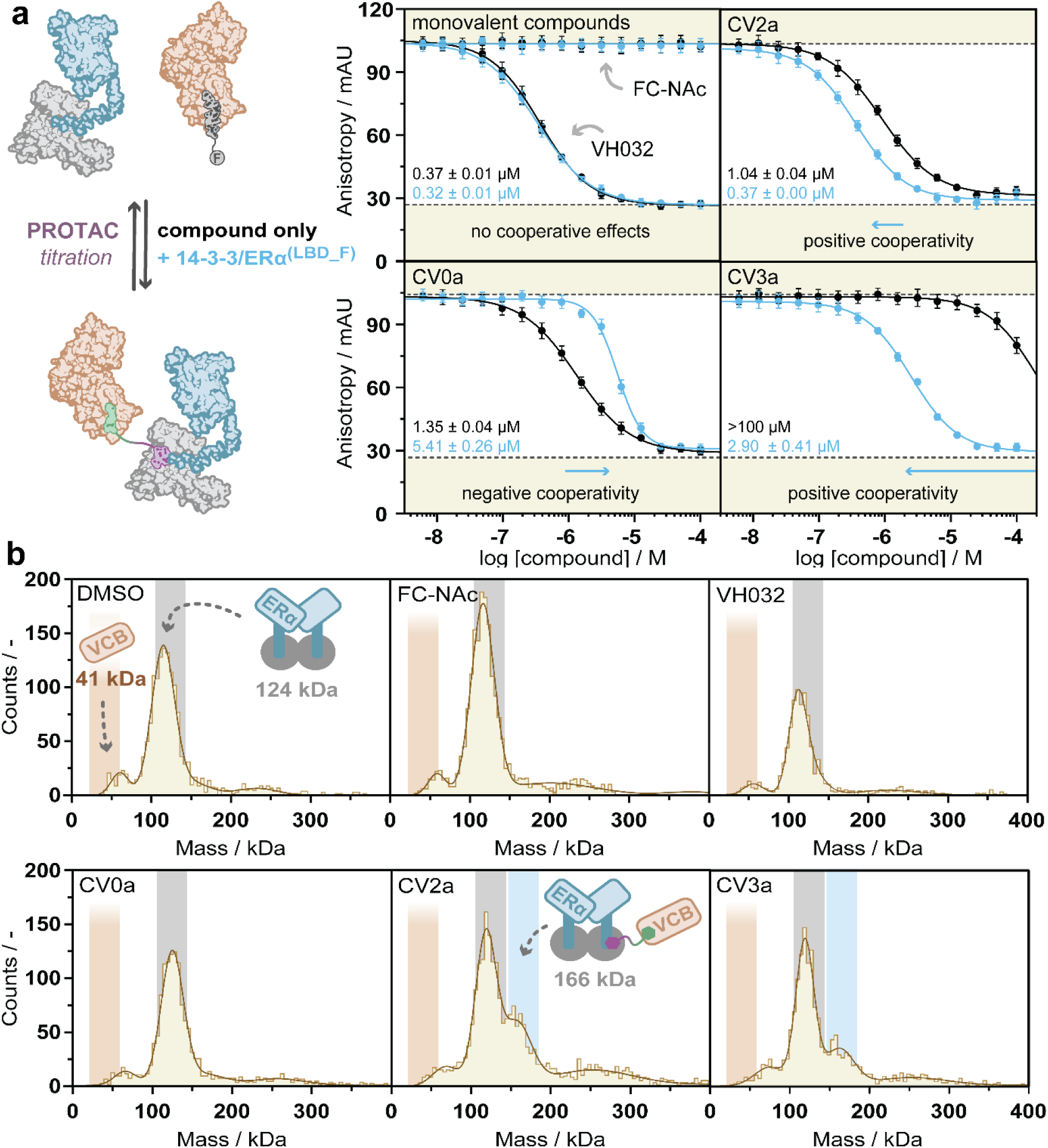
Cooperative VHL recruitment extends to the 14-3-3/ERα^(LBD_F)^ protein-protein complex. (a) FA assay data of ^MG^PROTAC (or monovalent compound) titration to 25 nM fluorescently labeled HIF1α peptide and 50 nM VCB protein complex in the absence (black) or presence of 10 µM 14-3-3/ERα^(LBD_F)^ complex (blue). IC_50_ values of displacement curves are indicated Mass photometry shows a single molecule of VCB binding the 14-3-3/ERα^(LBD_F)^ complex. A 1:1 mixture of 14-3-3/ERα^(LBD_F)^ complex (2:2 stoichiometry, 124 kDa) and the VCB complex (41 kDa) was treated with three equivalents of monovalent binder, bivalent compound or DMSO. Analysis of the histograms upon treatment of the protein mixture with DMSO, FC-NAc, VH032, or negatively cooperative CV0a showed two distinct peaks, corresponding to the VCB complex and the 14-3-3/ERα^(LBD_F)^ complex. In contrast, treatment with CV2a and CV3a promoted the formation of a larger complex consistent with a molecular weight of ∼166 kDa, corresponding to a 2:2:1 14-3-3/ ERα^(LBD_F)^ /VCB stoichiometry, suggesting an asymmetric binding of VCB to the complex.

**Extended Data Fig. 5.**
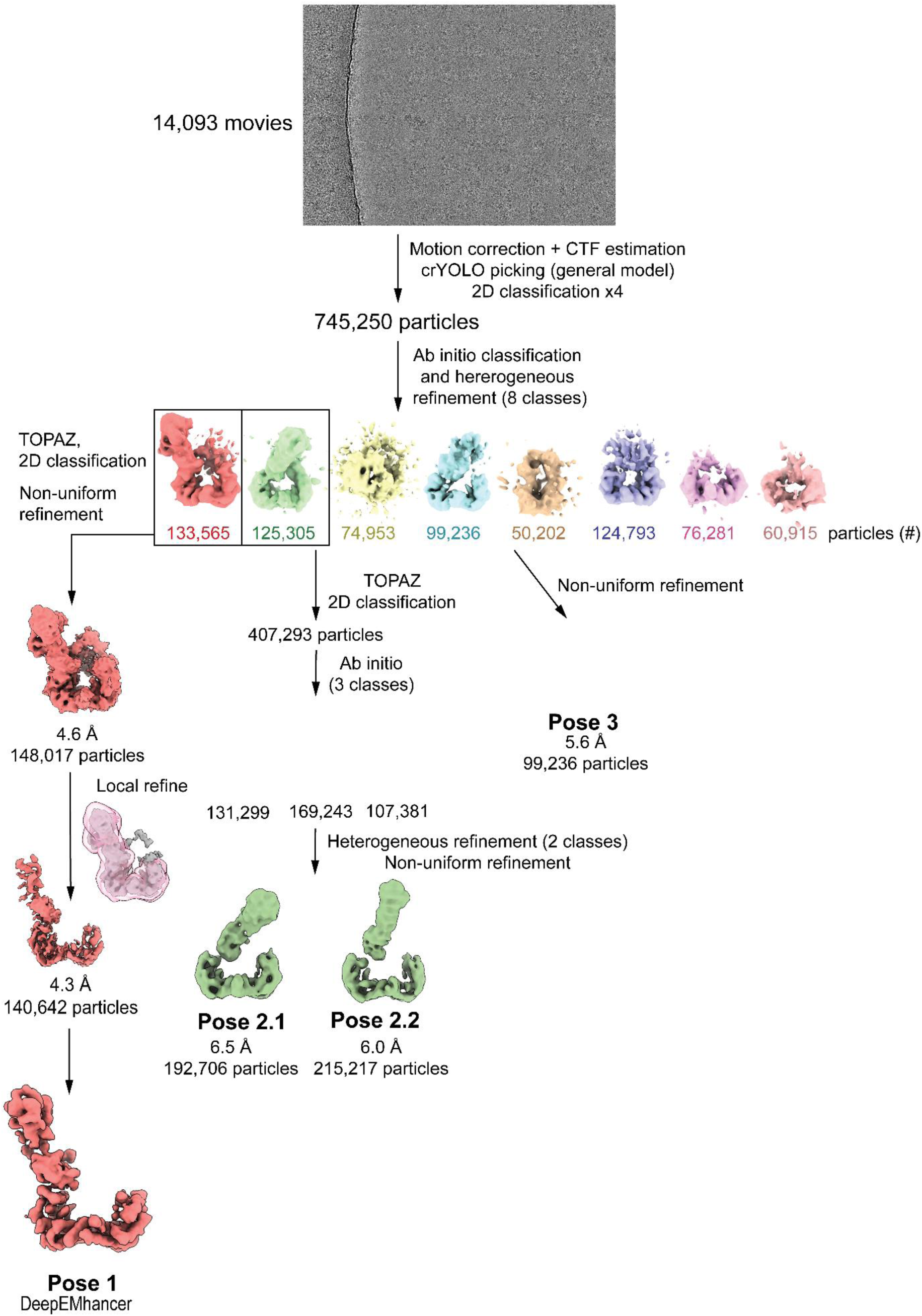
Cryo-EM analysis of the 14-3-3/ERα^(LBD_F)^/CV2a/VCB assembly.

**Extended Data Fig. 6.**
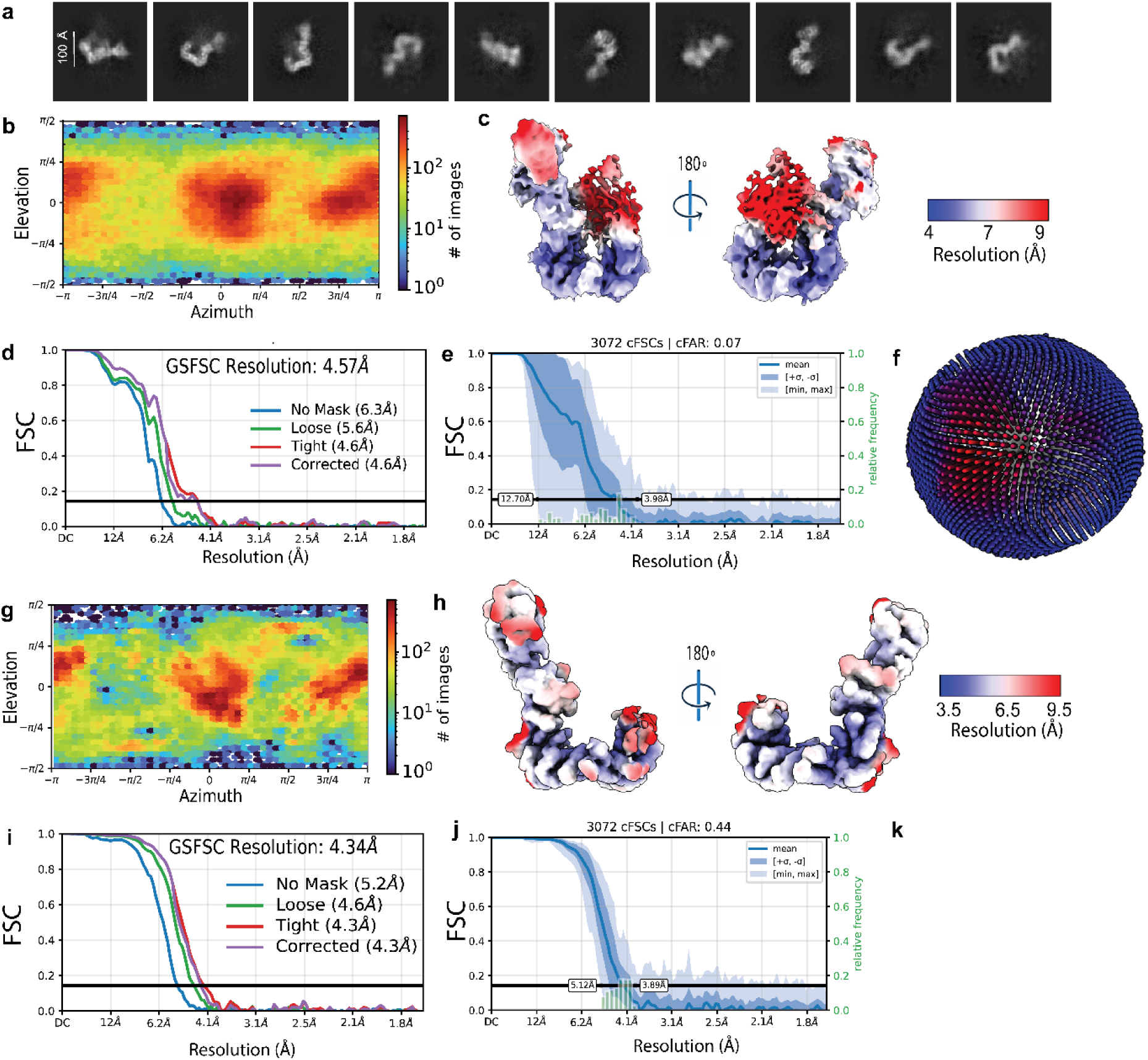
(a) Representative 2D classes associated with particles from Pose 1. (b) 2D viewing distribution for the Pose 1 volume before local refinement. (c) Local resolution estimation for the Pose 1 volume before local refinement. (d) Gold-standard Fourier shell correlation plot at 0.143 for the Pose 1 volume before local refinement. (e) Conical FSC distribution range for the Pose 1 volume before local refinement. (f) 3D viewing distribution for the Pose 1 volume before local refinement. (g) 3D viewing distribution for the Pose 1 volume after local refinement. (h) Local resolution estimation for the Pose 1 volume after local refinement. (i) Gold-standard Fourier shell correlation plot at 0.143 for the Pose 1 volume after local refinement. (j) Conical FSC distribution range for the Pose 1 volume after local refinement. (k) 3D viewing distribution for the Pose 1 volume after local refinement.

**Extended Data Fig. 7.**
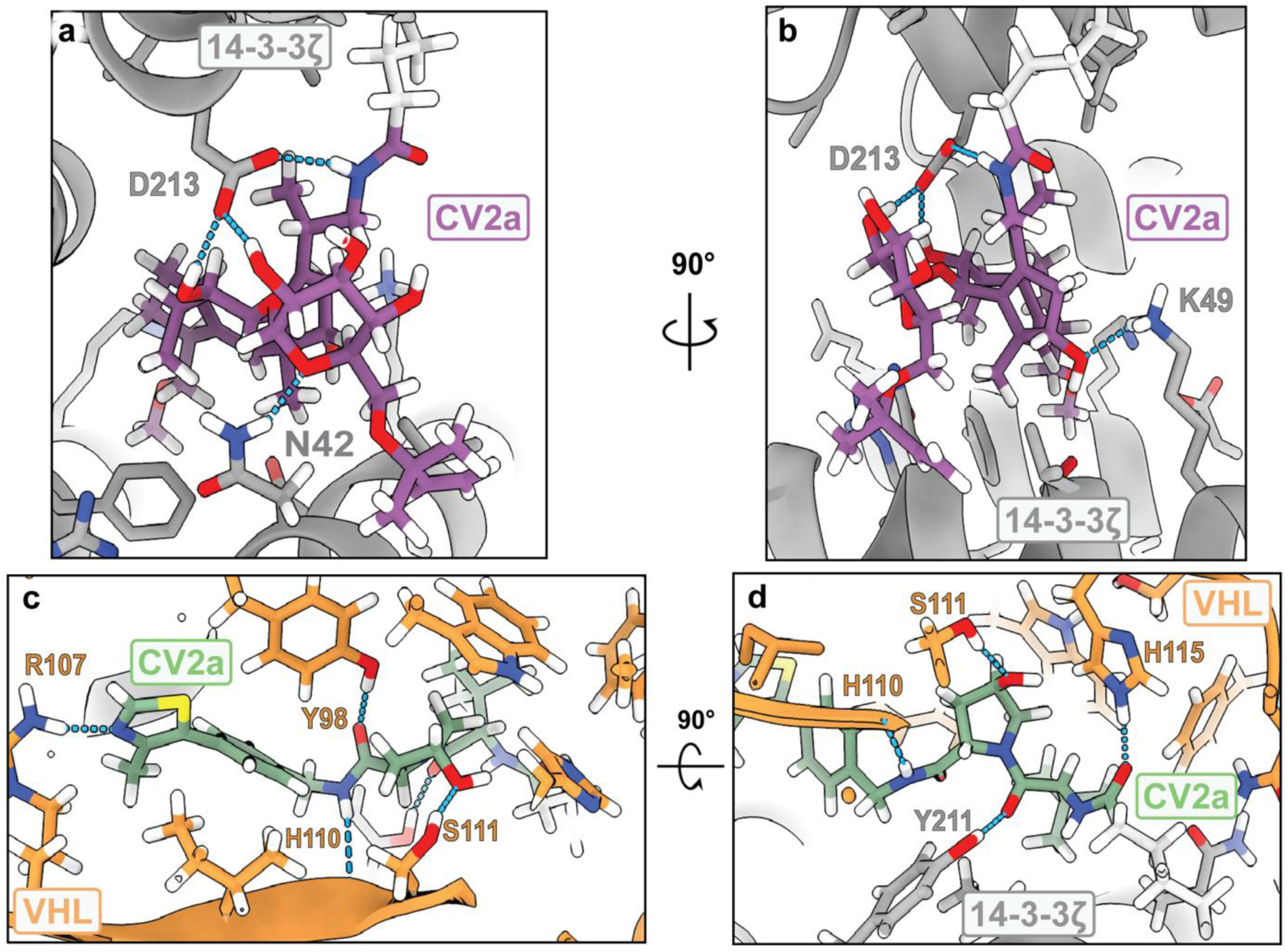
(a-b) Highlighted protein-ligand interactions between the fusicoccin ligand of CV2a and 14-3-3. The FC-NAc ligand of CV2a binds within the composite binding interface formed between binding groove of 14-3-3 and the C-terminal phosphorylated 14-3-3 binding motif of ERα. (c-d) Highlighted protein-ligand interactions between the VH032 ligand of CV2a, VHL, and 14-3-3.

**Extended Data Fig. 8.**
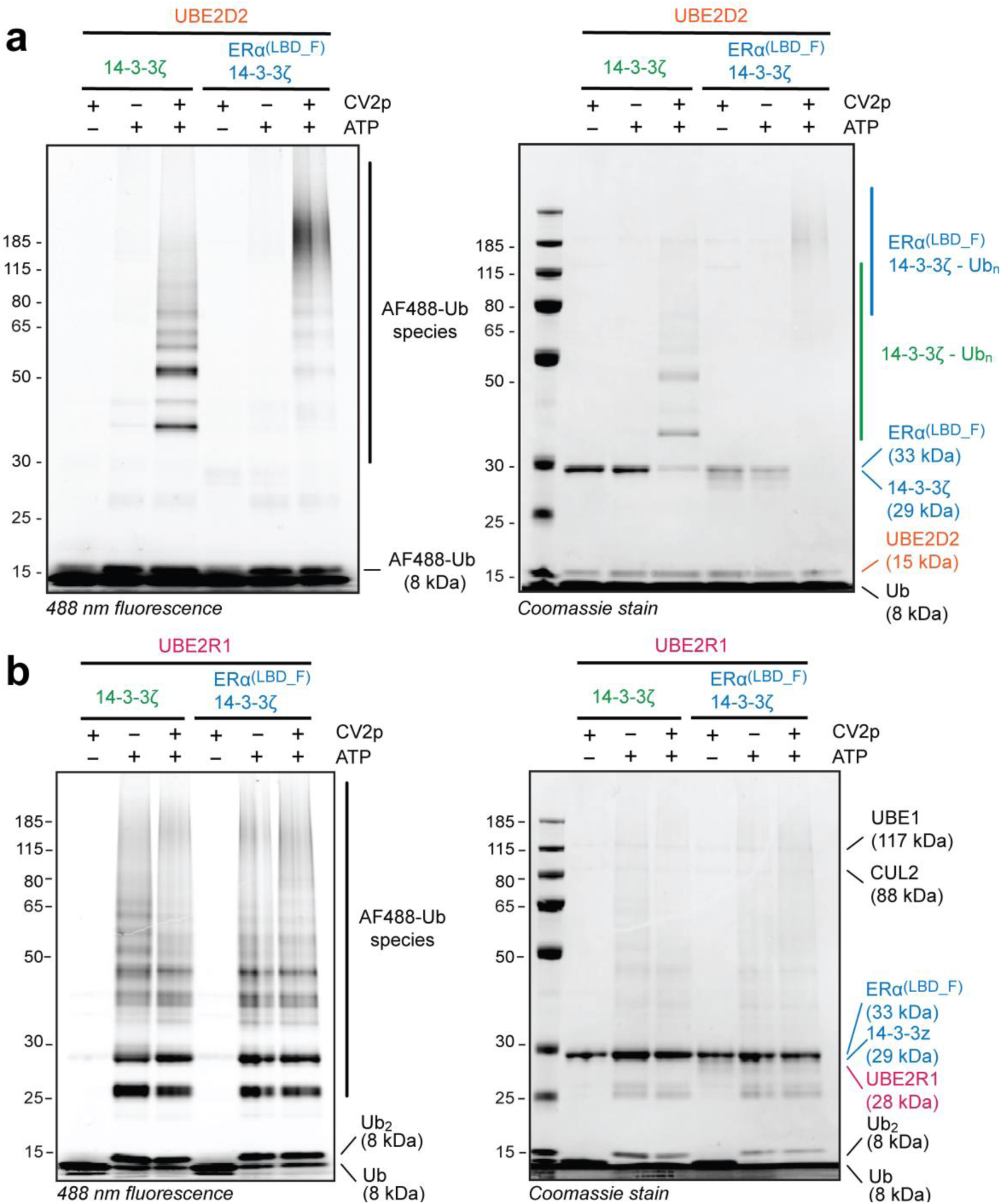
Ubiquitination assays with (a) UBE2D2 or (b) UBE2R1, and 14-3-3ζ or the 14-3-3ζ/ERα^(LBD_F)^ complex, visualized at 488 nm (Alexa Fluor 488-labelled ubiquitin) or Coomassie stain.

**Extended Data Fig. 9.**
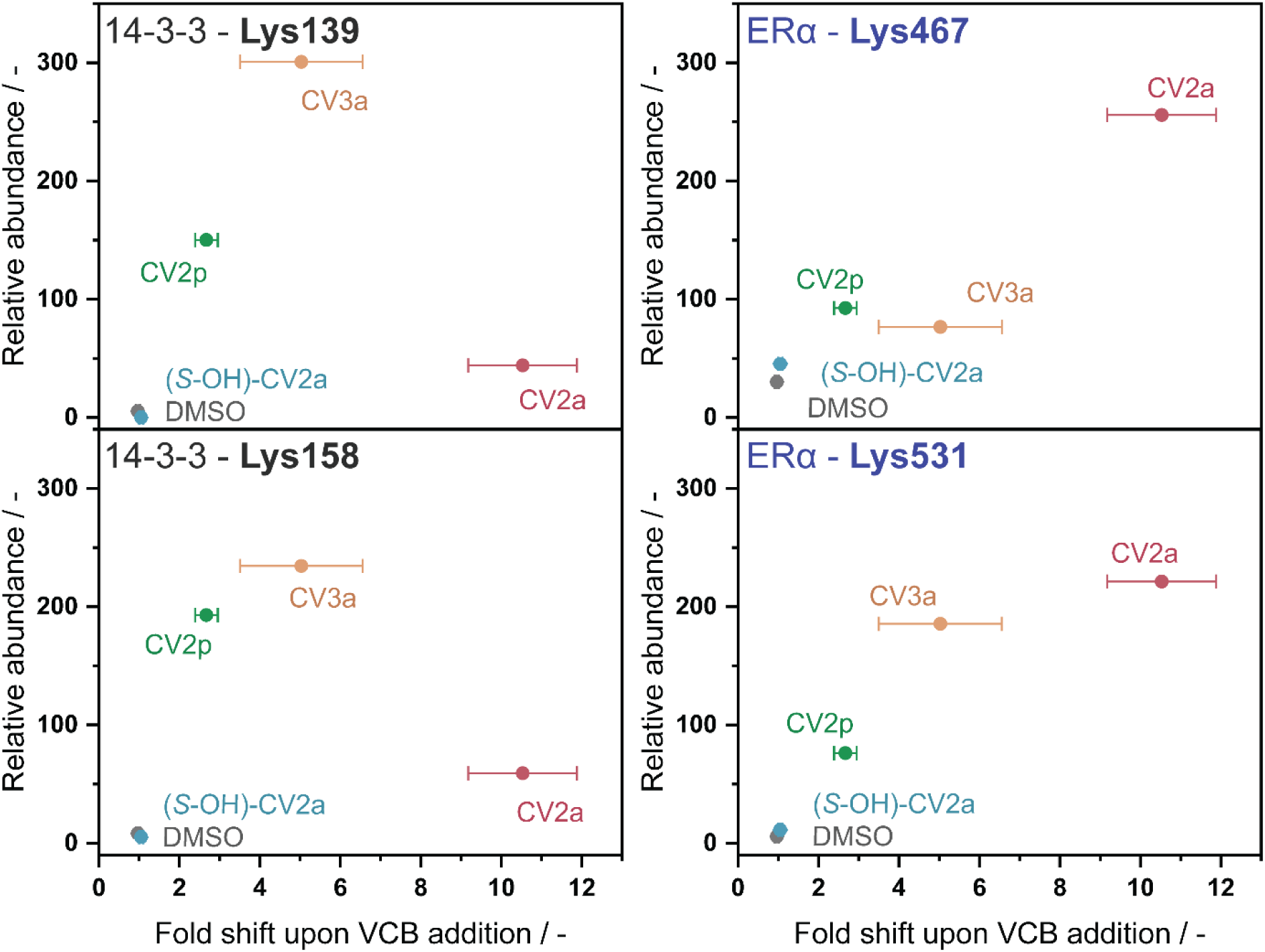
Comparison between ubiquitination in the in-vitro ubiquitination mass spectrometry experiment (vertical) and the cooperativity observed in the FA experiment (horizontal), expressed as fold 14-3-3 binding curve shift upon addition of VCB.

