## Supporting Information for "Induced ubiquitination of the partially disordered Estrogen Receptor alpha protein via a 14-3-3-directed molecular glue-based PROTAC design"

### Table of Contents

1. Supplementary Figures and Tables 3
2. Additional methods - chemistry 13

    Experimental 13
    NMR Spectra 25
    HPLC chromatograms 46

3. References 49

#### 1. Supplementary Figures and Tables

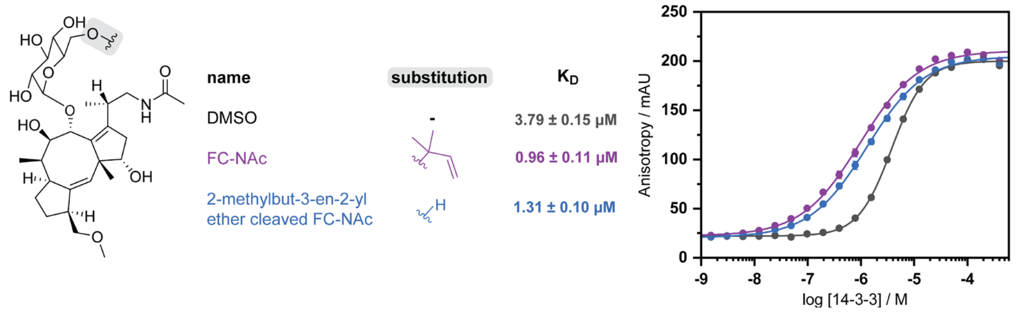

**Figure S1.** The 2-methylbut-3-en-2-yl ether cleaved FC-NAc stabilizes the 14-3-3/ERα interaction. FA 14-3-3 titrations in the presence of 1 µM FC-NAc or 1 µM 2-methylbut-3-en-2-yl ether cleaved FC-NAc, and 10 nM fluorescently labeled ERα peptide.

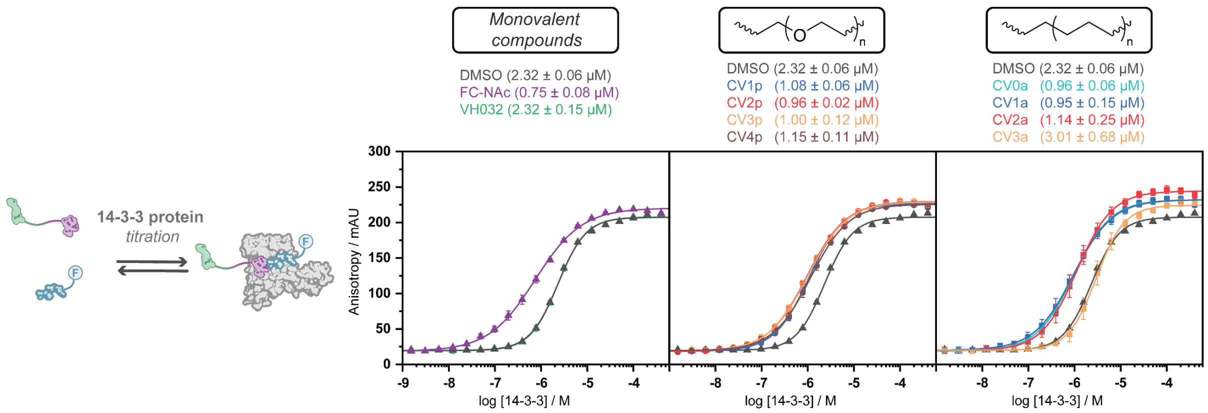

**Figure S2. All synthesized PROTACs, except CV3a, stabilize the 14-3-3/ERα interaction**. FA 14-3-3 titrations in the presence of 1 µM monovalent control compound, PEG linker based $^{MG}$PROTAC, or alkyl linker based $^{MG}$PROTAC, and 10 nM fluorescently labeled ERα peptide.

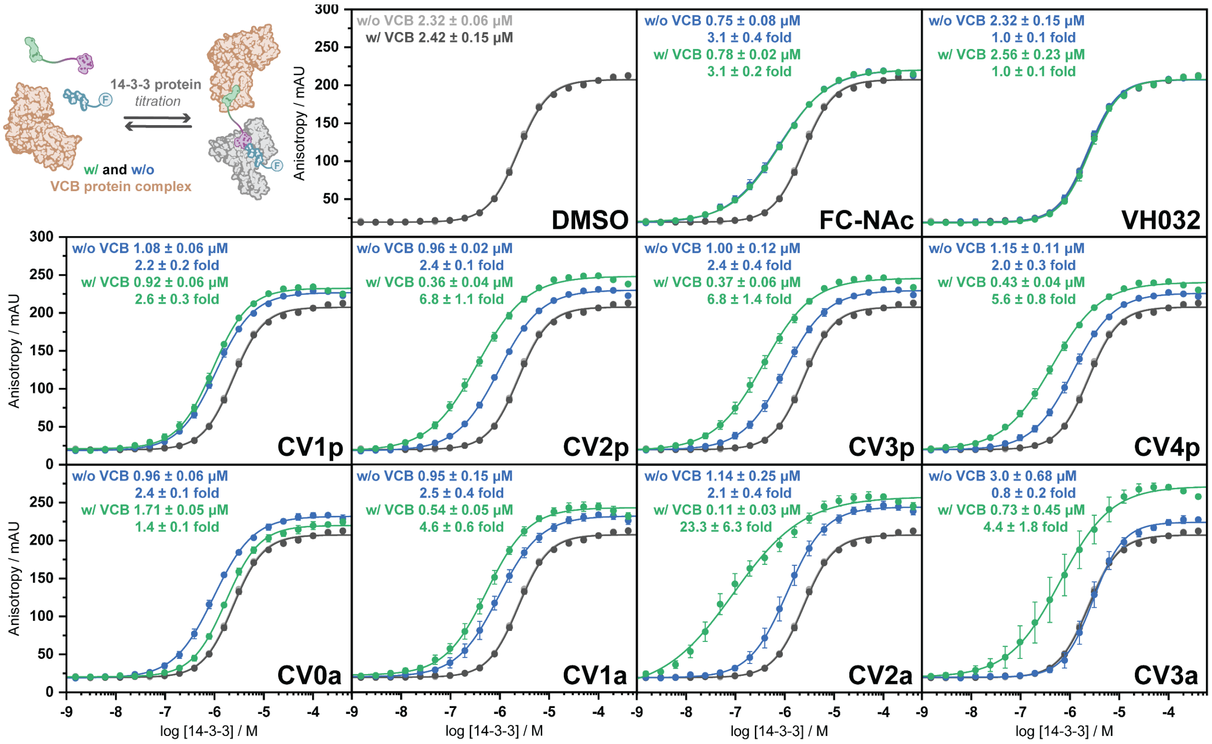

**Figure S3.** FA 14-3-3 titrations in the presence of 1 µM monovalent compound or $^{MG}$PROTAC, and 10 nM fluorescently labeled ERα peptide in the absence of VCB (w/o VCB, blue dots) and presence of 5 µM VCB (w/ VCB, green dots). The DMSO control is indicated in grey (w/o VCB light grey dots, w/ VCB dark grey dots, data overlaps). Error bars represent standard deviations of independent scientific replicates (n = 3).

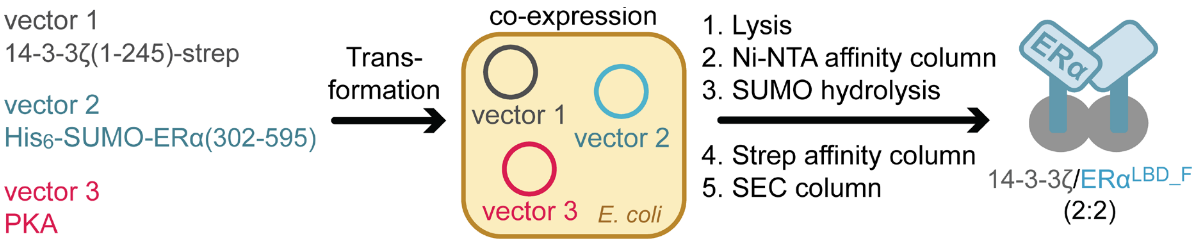

**Figure S4.** Overview of recombinant expression of 14-3-3ζ/ERα$^{(LBD_F)}$ complex. Adapted from Somsen *et al.* [1]

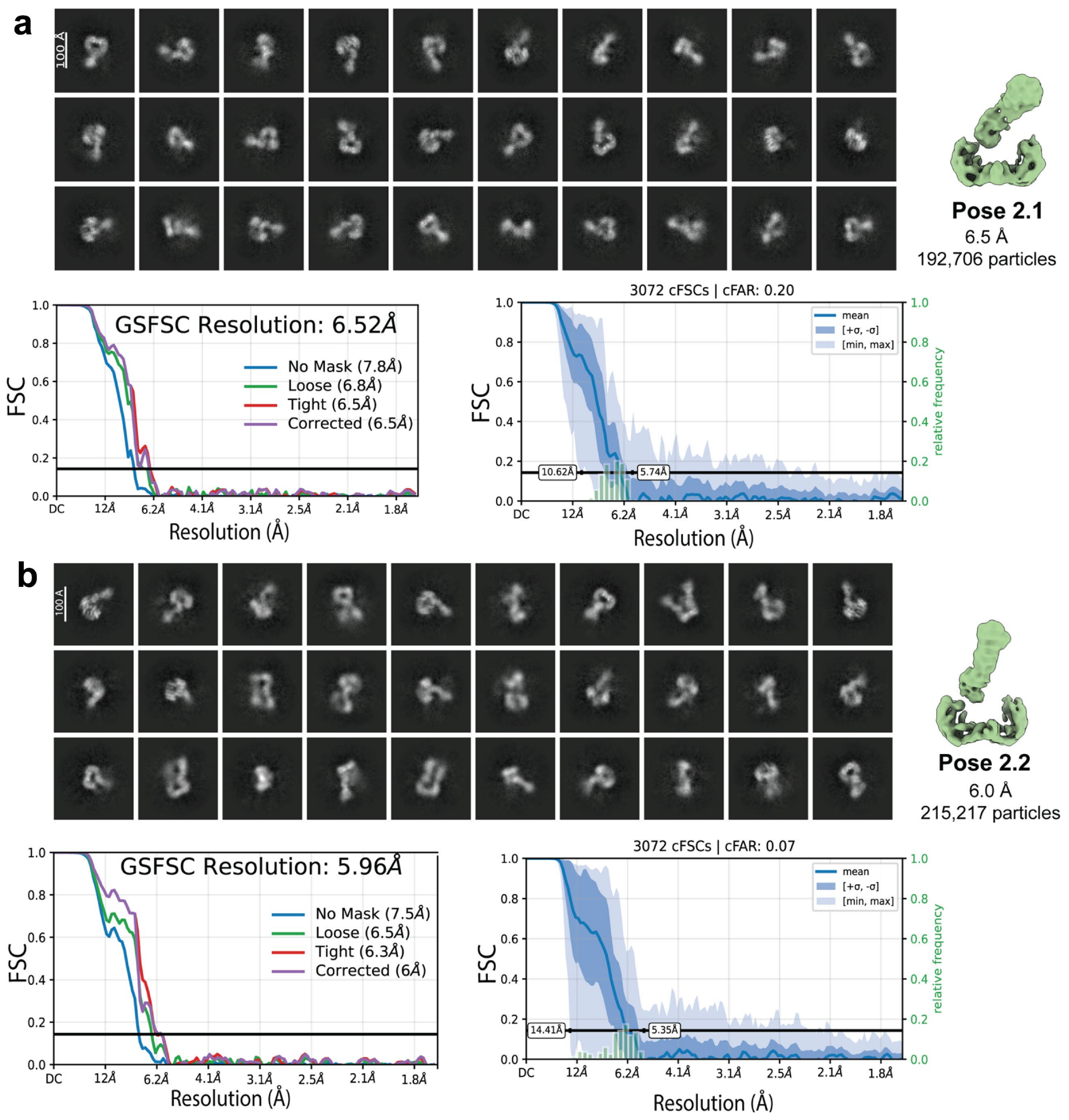

**Figure S5.** (a) Pose 2.1 Representative 2D classes, Gold-standard Fourier shell correlation plot at 0.143 and conical FSC distribution range. (b) Pose 2.2 Representative 2D classes, Gold-standard Fourier shell correlation plot at 0.143 and conical FSC distribution range.

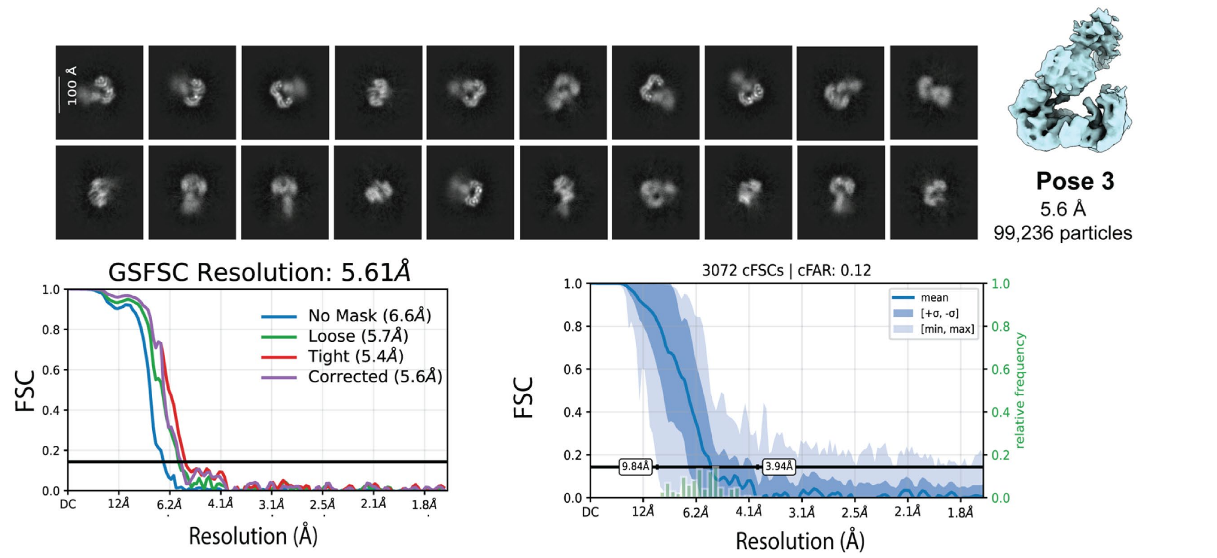

**Figure S6.** Pose 3 Representative 2D classes, Gold-standard Fourier shell correlation plot at 0.143 and conical FSC distribution range.

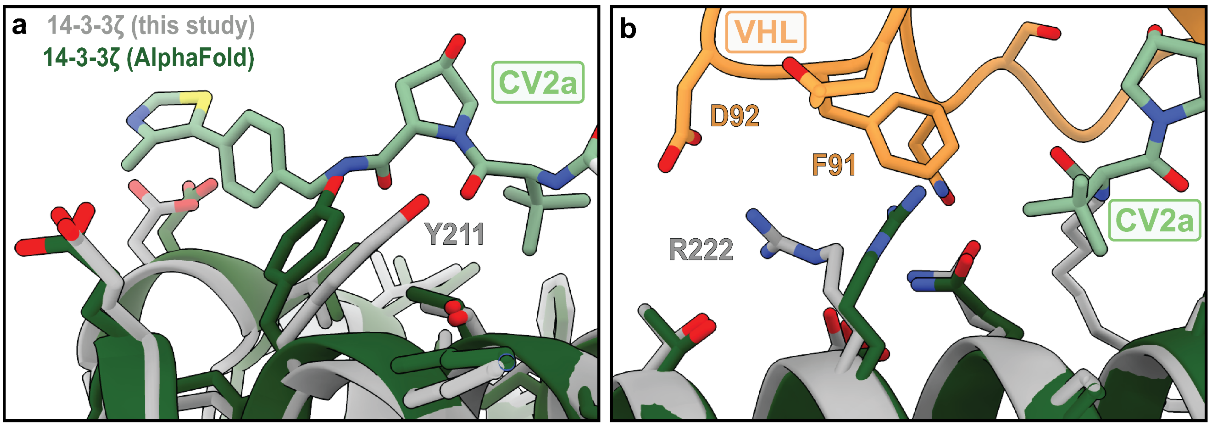

**Figure S7.** (a) Superposition of apo 14-3-3 zeta (AlphaFold, dark green) with 14-3-3 zeta from the model in this study (silver). A Shows Tyr211 rotate to accommodate the VH032 moiety of CV2a. (b) Shows R222 flip to avoid clashing with Phe91 and to form contacts with Asp92.

```
14-3-3β (P31946)  3    MDKSELVQKAKLAEQAERYDDMAAAMKAVTEQGHELSNEERNLLSVAYKNVVGARRSSWR  62
14-3-3ε (P62258)  3     DREDLVYQAKLAEQAERYDEMVESMKKVAGMDVELTVEERNLLSVAYKNVIGARRASWR  61
14-3-3η (Q04917)  3     DREQLLQRARLAEQAERYDDMASAMKAVTELNEPLSNEDRNLLSVAYKNVVGARRSSWR  61
14-3-3γ (P61981)  2    VDREQLVQKARLAEQAERYDDMAAAMKNVTELNEPLSNEERNLLSVAYKNVVGARRSSWR  61
14-3-3σ (P31947)  1    MERASLIQKAKLAEQAERYEDMAAFMKGAVEKGEELSCEERNLLSVAYKNVVGGQRAAWR  60
14-3-3θ (P27348)  1    MEKTELIQKAKLAEQAERYDDMATCMKAVTEQGAELSNEERNLLSVAYKNVVGGRRSAWR  60
14-3-3ζ (P63104)  1    MDKNELVQKAKLAEQAERYDDMAACMKSVTEQGAELSNEERNLLSVAYKNVVGARRSSWR  60
                         helix 1           helix 2              helix 3

14-3-3β (P31946)  63   VISSIEQKT--ER--NEKKQQMGKEYREKIEAELQDICNDVLELLDKYLIPNA--T--QP  114
14-3-3ε (P62258)  62   IISSIEQKE--ENKGGEDKLKMIREYRQMVETELKLICCDILDVLDKHLIPAA--N--TG  115
14-3-3η (Q04917)  62   VISSIEQKTMADG--NEKKLEKVKAYREKIEKELETVCNDVLSLLDKFLIKNC--NDFQY  117
14-3-3γ (P61981)  62   VISSIEQKTSADG--NEKKIEMVRAYREKIEKELEAVCQDVLSLLDNYLIKNCSET--QY  117
14-3-3σ (P31947)  61   VLSSIEQKSNEEG--SEEKGPEVREYREKVETELQGVCDTVLGLLDSHLIKEA--G--DA  114
14-3-3θ (P27348)  61   VISSIEQKT--DT--SDKKLQLIKDYREKVESELRSICTTVLELLDKYLIANA--T--NP  112
14-3-3ζ (P63104)  61   VVSSIEQKT--EG--AEKKQQMAREYREKIETELRDICNDVLSLLEKFLIPNA--S--QA  112
                                             helix 4

14-3-3β (P31946)  115  ESKVFYLKMKGDYFRYLSEVASGDNKQTTVSNSQQAYQEAFEISKKEMQPTHPIRLGLAL  174
14-3-3ε (P62258)  116  ESKVFYYKMKGDYHRYLAEFATGNDRKEAAENSLVAYKAASDIAMTELPPTHPIRLGLAL  175
14-3-3η (Q04917)  118  ESKVFYLKMKGDYYRYLAEVASGEKKNSVVEASEAAYKEAFEISKEQMQPTHPIRLGLAL  177
14-3-3γ (P61981)  118  ESKVFYLKMKGDYYRYLAEVATGEKRATVVESSEKAYSEAHEISKEHMQPTHPIRLGLAL  177
14-3-3σ (P31947)  115  ESRVFYLKMKGDYYRYLAEVATGDDKKRIIDSARSAYQEAMDISKKEMPPTNPIRLGLAL  174
14-3-3θ (P27348)  113  ESKVFYLKMKGDYFRYLAEVACGDDRKQTIDNSQGAYQEAFDISKKEMQPTHPIRLGLAL  172
14-3-3ζ (P63104)  113  ESKVFYLKMKGDYYRYLAEVAAGDDKKGIVDQSQQAYQEAFEISKKEMQPTHPIRLGLAL  172
                          helix 5                   helix 6            helix 7
                                                          204  208 211     219 222
14-3-3β (P31946)  175  NFSVFYYEILNSPEKACSLAKTAFDEAIAELDTLNEESYKDSTLIMQLLRDNLTLWTSEN  234
14-3-3ε (P62258)  176  NFSVFYYEILNSPDRACRLAKAAFDDAIAELDTLSEESYKDSTLIMQLLRDNLTLWTSDM  235
14-3-3η (Q04917)  178  NFSVFYYEIQNAPEQACLLAKQAFDDAIAELDTLNEDSYKDSTLIMQLLRDNLTLWTSDQ  237
14-3-3γ (P61981)  178  NYSVFYYEIQNAPEQACHLAKTAFDDAIAELDTLNEDSYKDSTLIMQLLRDNLTLWTSDQ  237
14-3-3σ (P31947)  175  NFSVFHYEIANSPEEAISLAKTTFDEAMADLHTLSEDSYKDSTLIMQLLRDNLTLWTADN  234
14-3-3θ (P27348)  173  NFSVFYYEILNNPELACTLAKTAFDEAIAELDTLNEDSYKDSTLIMQLLRDNLTLWTSDS  232
14-3-3ζ (P63104)  173  NFSVFYYEILNSPEKACSLAKTAFDEAIAELDTLSEESYKDSTLIMQLLRDNLTLWTSDT  232
                                   helix 8                   helix 9

14-3-3β (P31946)  235  QGDEGDAGEGEN   246
14-3-3ε (P62258)  236  QGD----GEEQN   243
14-3-3η (Q04917)  238  QDEE--AGEG     245
14-3-3γ (P61981)  238  QDDDG--GEGNN   247
14-3-3σ (P31947)  235  AGEEG          239
14-3-3θ (P27348)  233  AGEECDAAEG     242
14-3-3ζ (P63104)  233  QGDEAEAGEG     242
```

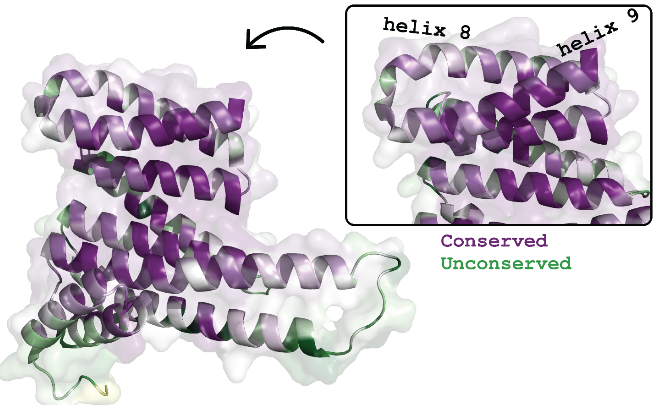

**Figure S8:** Sequence alignments of 14-3-3 isoforms (β, ε, η, γ, σ, θ, and ζ ). The residues forming contacts with VHL in the cryo-EM model in this study are highlighted (purple). Alignment performed by BLAST.[2]

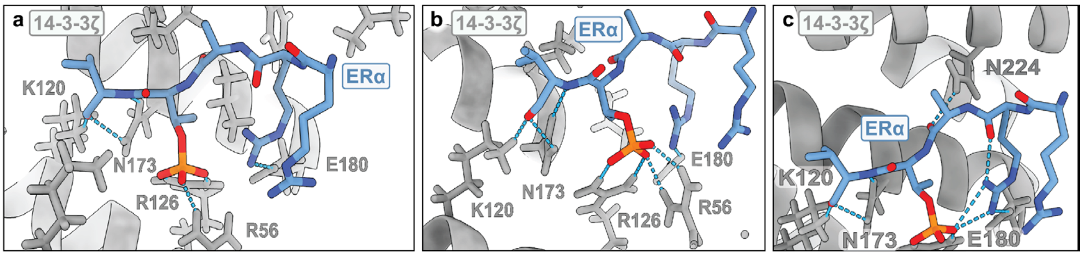

**Figure S9.** Highlighted protein-protein interactions between 14-3-3 and the ERα C-terminus.

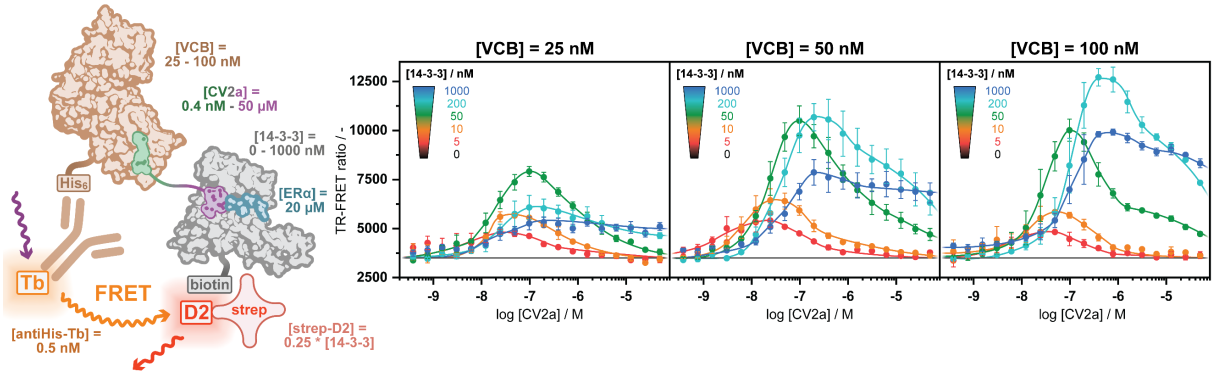

**Figure S10.** TR-FRET assay optimization. Concentrations of His$_6$-tagged VCB protein complex (25 nM – 100 nM), biotinylated 14-3-3 (0 nM – 1 µM; each concentration with 0.25 eq. Strep-D2), and CV2a (0.4 nM – 50 µM) were varied, and the concentrations of Anti-His-Tb (0.5 nM), and ERα peptide (20 µM) were kept constant. 50 nM His$_6$-tagged VCB, 50 nM biotinylated 14-3-3 (with 12.5 nM Strep-D2) were selected as optimal concentrations due to the large TR-FRET signal at low CV2a concentrations.

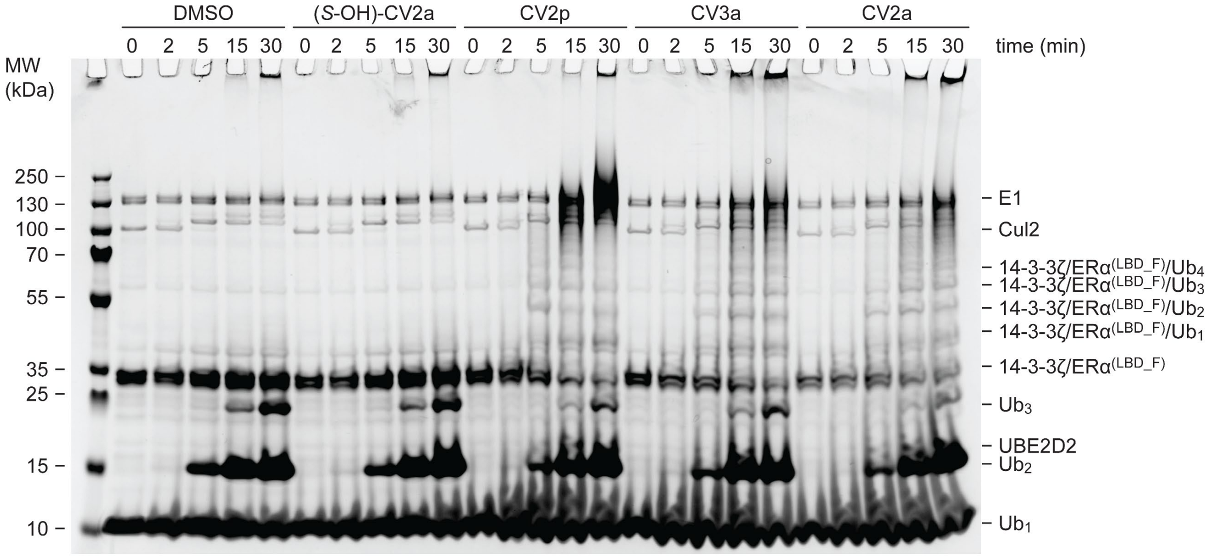

**Figure S11.** Full SDS-PAGE gel of in-vitro time course ubiquitination experiment from time point 0 to 30 mins, Coomassie staining.

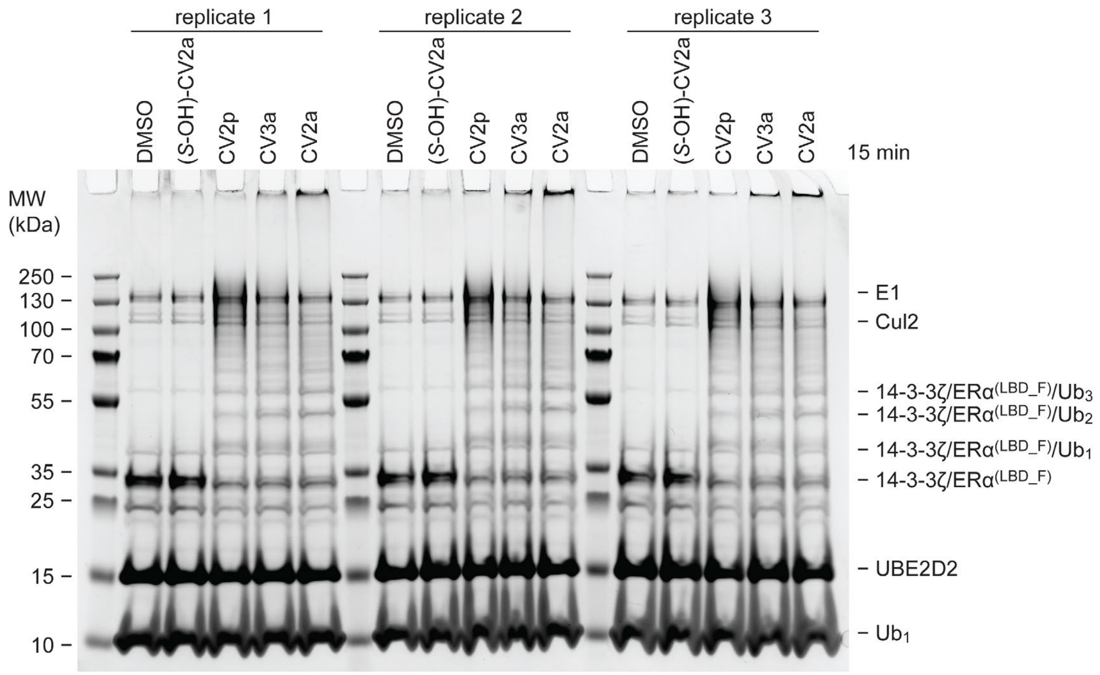

**Figure S12.** Full SDS-PAGE gel of in-vitro ubiquitination experiment comparing control molecules and $^{MG}$PROTAC replicates, Coomassie staining. Overnight incubation.

**Table S1.** Overview of tryptic peptides measured by mass spectrometry in the in-vitro ubiquitination experiment.

| Protein | Annotated Sequence | Modifications | PSMs | Missed cleavages | Compound | | | | |
|---|---|---|---|---|---|---|---|---|---|
| | | | | | DMSO | CV2p | CV2a | (*S*-OH)-CV2a | CV3a |
| ERα | [K].CKNVVPLYDLLLEMLDAHR.[L] | 1x Carbamidomethyl [C1]; 1x GG [K2] | 29 | 1 | 30 | 92.5 | 255.7 | 45.3 | 76.5 |
| ERα | [K].SIILLNSGVYTFLSSTLK.[S] | 1x GG [K18] | 3 | 0 | | | | | 500 |
| ERα | [K].SIILLNSGVYTFLSSTLKSLEEK.[D] | 1x GG [K18] | 56 | 1 | 5.7 | 76.2 | 221.3 | 11.3 | 185.5 |
| ERα | [R].GGASVEETDQSHLATAGSTSSHSLQKYYITGEAEGR.[R] | 1x GG [K26] | 1 | 1 | | | | | |
| ERα | [R].GGASVEETDQSHLATAGSTSSHSLQK.[Y] | 1x Phospho [S/T] | 5 | 0 | 143.1 | 110.4 | 115.8 | 130.7 | |
| ERα | [R].GGASVEETDQSHLATAGSTSSHSLQK.[Y] | 1x Phospho [S23]; 1x GG [K26] | 4 | 0 | 58.2 | 168.7 | 77.3 | 195.8 | |
| ERα | [K].GMEHLYSMK.[C] | 2x Oxidation [M2; M8] | 1 | 0 | 40.5 | 130.9 | 121 | 103.2 | 104.5 |
| 14-3-3ζ | [K].KGIVDQSQQAYQEAFEISKK.[E] | 1x GG [K1] | 4 | 2 | | | 500 | | |
| 14-3-3ζ | [K].KEMQPTHPIR.[L] | 1x GG [K1] | 27 | 1 | 8.3 | 192.9 | 59.2 | 5.2 | 234.4 |
| 14-3-3ζ | [K].KGIVDQSQQAYQEAFEISK.[K] | 1x GG [K1] | 16 | 1 | 5.3 | 150.1 | 44 | | 300.7 |
| 14-3-3ζ | [R].YLAEVAAGDDKK.[G] | 1x GG [K11] | 3 | 1 | | 125.5 | 91.4 | | 283.1 |
| 14-3-3ζ | [K].GIVDQSQQAYQEAFEISKK.[E] | 1x GG [K18] | 1 | 1 | | | | | |
| 14-3-3ζ | [K].TAFDEAIAELDTLSEESYKDSTLIMQLLR.[D] | 1x GG [K19] | 10 | 1 | 56.8 | | 68.2 | | 375 |
| 14-3-3ζ | [K].AKLAEQAER.[Y] | 1x GG [K2] | 1 | 1 | | 371.6 | | | 128.4 |
| 14-3-3ζ | [-].MDKNELVQK.[A] | 1x GG [K3] | 5 | 1 | | 343.5 | 78 | | 78.5 |
| 14-3-3ζ | [R].DNLTLWTSDTQGDEAEAGEGGENWSHPQFEK.[-] | 1x GG [K31] | 15 | 0 | 18.6 | 171.9 | 141.2 | 18.4 | 149.9 |
| 14-3-3ζ | [K].DSTLIMQLLRDNLTLWTSDTQGDEAEAGEGGENWSHPQFEK.[-] | 1x GG [K41] | 16 | 1 | 0.3 | 134 | 349.1 | 4.4 | 12.3 |
| 14-3-3ζ | [K].NELVQKAK.[L] | 1x GG [K6] | 4 | 1 | | | | | |
| 14-3-3ζ | [R].VVSSIEQKTEGAEK.[K] | 1x GG [K8] | 1 | 1 | | | | | |
| 14-3-3ζ | [K].KEMQPTHPIR.[L] | 1x Oxidation [M3]; 1x GG [K1] | 10 | 1 | | 166.7 | 18.2 | | 315.1 |
| 14-3-3ζ | [K].DSTLIMQLLRDNLTLWTSDTQGDEAEAGEGGENWSHPQFEK.[-] | 1x Oxidation [M6]; 1x GG [K41] | 2 | 1 | | | 500 | | |
| 14-3-3ζ | [K].SVTEQGAELSNEER.[N] | 1x Phospho [S1] | 5 | 0 | 80.9 | 133.9 | 85.7 | 90.7 | 108.7 |
| 14-3-3ζ | [K].FLIPNASQAESK.[V] | 1x Phospho [S11]; 1x GG [K12] | 2 | 0 | 117.5 | 16.2 | 71.7 | 278.8 | 15.7 |
| 14-3-3ζ | [K].TAFDEAIAELDTLSEESYK.[D] | 1x Phospho [S17] | 1 | 0 | | | | | |
| 14-3-3ζ | [K].GIVDQSQQAYQEAFEISKK.[E] | 1x Phospho [S17]; 1x GG [K] | 3 | 1 | 57.2 | 72.5 | 121.7 | 248.7 | |
| 14-3-3ζ | [K].TAFDEAIAELDTLSEESYK.[D] | 1x Phospho [Y18]; 1x GG [K19] | 8 | 0 | 100.6 | 114.6 | 111.2 | 161 | 12.5 |
| 14-3-3ζ | [R].YLAEVAAGDDKKGIVDQSQQAYQEAFEISK.[K] | 2x GG [K11; K12] | 1 | 2 | | | | | |

| *Data collection* | *Dataset which yielded volumes: EMD-55233, EMD-55234, EMD-55235, EMD-55236, EMD-55237* |
|---:|:---:|
| *Microscope* | Krios |
| *Detector* | Gatan K3 (counting mode) |
| *Voltage (kV)* | 300 |
| *Magnification (nominal)* | 105,000x |
| *Number of fractions* | 40 |
| *Total electron exposure ($e^{-}/Å^{2}$)* | 33 |
| *Defocus range (µm)* | -1.2 to -2.7 |
| *Pixel size at detector (Å/pixel)* | 0.829 |
| *Total exposure (s)* | 1.75 |
| *Dose rate ($e^{-}$/px/s)* | 15.7 |
| *Automation software* | EPU v2.1 |
| *Energy filter slit width (eV)* | 20 |
| *C2 aperture (µm)* | 70 |
| *Movies collected (#)* | 14,093 |

**Table S2.** Cryo-EM data collection parameters.

| ***Reconstruction*** | *Pose 1 EMDB-55234* | *Local refinement of 'Pose 1' EMDB-55233 PDB-9SV3* | *Pose 2.1 EMDB-55235* | *Pose 2.2 EMDB-55236* | *Pose 3 EMDB-55237* |
|---:|:---:|:---:|:---:|:---:|:---:|
| *Image processing package* | CryoSPARC | CryoSPARC | CryoSPARC | CryoSPARC | CryoSPARC |
| *Symmetry imposed* | C1 | C1 | C1 | C1 | C1 |
| *Initial particles (#)* | 148,017 | 148,017 | 192,706 | 215,217 | 99,236 |
| *Final particles (#)* | 148,017 | 140,642 | 192,706 | 215,217 | 99,236 |
| *Map resolution (Å) at FSC 0.143* | 4.6 | 4.3 | 6.5 | 6.0 | 5.6 |
| *B Factor ($Å^{2}$)* | 280 | 382 | 489 | 475 | 436 |
| *Map resolution range (Å)* | 4.0 to 12.7 | 3.9 to 5.1 | 5.6 to 10.6 | 5.3 to 14.4 | 3.9 to 9.8 |
| *SCF** | 0.892 | 0.814 | 0.971 | 0.943 | 0.944 |
| *cFAR* | 0.07 | 0.44 | 0.20 | 0.07 | 0.12 |

**Table S3.** Cryo-EM data image analysis.

| ***Model composition*** | *Local refinement of 'Pose 1' EMDB-55233 PDB-9SV3* |
|---:|:---:|
| *Proteins* | VHL, EloC, EloB, 14-3-3zeta, estrogen receptor alpha |
| *Ligands* | CV2a |
| ***Model refinement*** | |
| *Atomic modelling packages* | iSOLDE, Phenix |
| *Initial model(s) used* | 4W9H, 8C43, 8BZH, AlphaFold |
| *Cross-correlation (map/model)* | 0.35 |
| *R.m.s deviations from ideal values:* | |
| *- Bond lengths (Å)* | 0.014 |
| *- Bond angles (°)* | 1.854 |
| *Protein residues B Factor(Å)* | 12.68 |
| *Ligand B Factor (Å)* | 16.52 |
| ***Validation*** | |
| *MolProbity score* | 1.84 |
| *Clashscore (all atoms)* | 13.42 |
| *Poor rotamers (%)* | 0.83 |
| *Ramachandran - favoured (%)* | 96.77 |
| *Ramachandran outliers (%)* | 0.00 |
| *EMRinger score* | 0.39 |
| *CaBLAM outliers (%)* | 0.63 |

**Table S4.** Cryo-EM data atomic modelling, refinement, and validation statistics.

### 2. Additonal methods - chemistry

**General chemistry information**

All reagents were purchased from TCI, Sigma-Aldrich, or BLDPharm and were used without further purification. All solvents were of analytical grade and supplied by Biosolve. Reaction glassware was dried at 130 °C for more than 24 hours prior to use. TLC was carried out on aluminum-backed silica (Merck silica gel 60 $F_{254}$) plates supplied by Merck. Visualization of the plates was achieved using an ultraviolet lamp ($\lambda_{max}$ = 254 nm) or by phosphomolybdic acid staining. Reversed-phase preparative HPLC was performed using a C18 Atlantis T3 OBD 150 x 19 mm column using miliQ water with 0.1% formic acid (FA) and acetonitrile (ACN) with 0.1% FA. Analytical (LR) LC-MS (ESI) analysis was performed on a system using comprising a C18 Jupiter SuC4300A 150 x 2.0 mm column using water with 0.1% FA and acetonitrile with 0.1% FA, using a gradient of 5% to 100% acetonitrile over 8 minutes, connected to a Thermo Fisher LTQ XL Linear Ion Trap Mass Spectrometer. High resolution mass spectra (HRMS) were recorded using a Waters ACQUITY UPLC I-Class LC system coupled to a Xevo G2 Quadrupole Time of Flight (Q-TOF) mass spectrometer equipped with a Phenomex kinetex® 2.6 µm EVO C18 100 x 2.1 mm column. Proton ($^1$H) and carbon ($^{13}$C) NMR spectra were collected on a 400 MHz Bruker Cryomagnet. Chemical shifts (δ) are referenced to the residual solvent peak. Splitting patterns are reported in an abbreviated manner: s (singlet), d (doublet), t (triplet), q (quartet), and m (multiplet). The percentages of 2-methylbut-3-en-2-yl ether cleavage of the final PROTAC compounds are estimated by HPLC and $^1$H NMR.

**Synthesis overview**

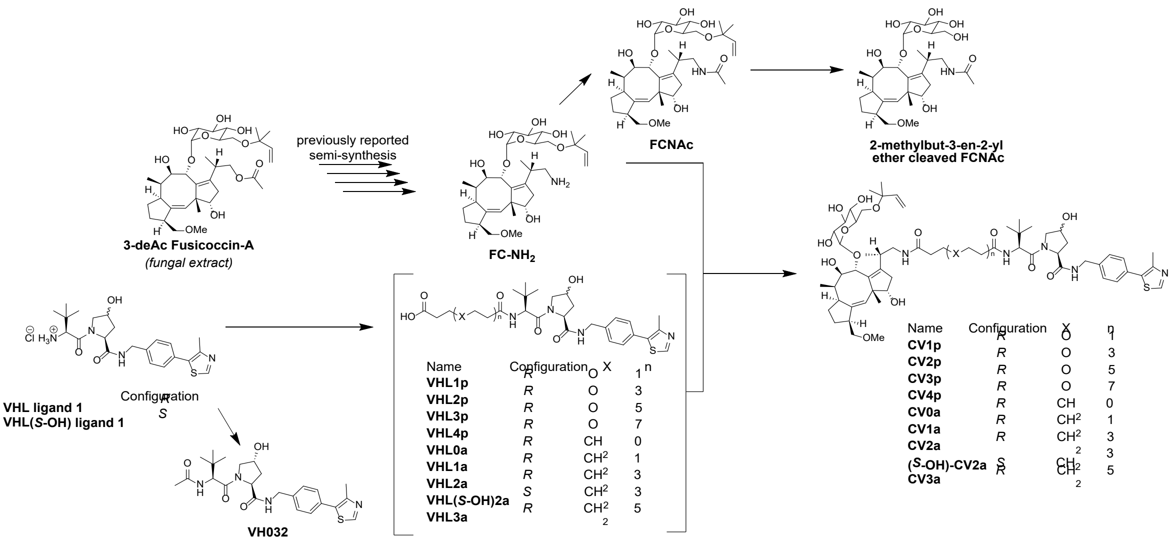

**General procedure A: acetylation**

To a solution of amine or suspension of amine HCl salt (0.0588-0.214 mmol, 1 eq) in dry chloroform (3-5 mL) at 0° C was added DIPEA (0.294-1.07 mmol, 5 eq). The resulting solution was stirred for 10 min and acetic anhydride (0.0588-0.214 mmol, 1 eq) was slowly added over 5 min. The reaction mixture was allowed to warm to room temperature, stirred for 16h and quenched by the addition of water (5 mL per mmol). The resulting mixture was concentrated under reduced pressure and purified by reversed-phase preparative HPLC to yield the desired product.

**General procedure B: coupling VHL to diacid**

To a solution of diacid (0.642-1.07 mmol, 3-5 eq) and HATU (0.600-0.964 mmol, 2.8-4.5 eq) in dry DMF (2-17 mL) was added DIPEA (6.42 mmol, 30 eq). The resulting solution was stirred for 10 min and VHL ligand 1 HCl salt (0.214 mmol, 1 eq) dissolved in dry DMF (3 mL) was slowly added over 5 min. The reaction mixture was then stirred for 16h and quenched by the addition of water (5 mL per mmol). The resulting mixture was concentrated under reduced pressure and purified by reversed-phase preparative HPLC to yield the desired product.

**General procedure C: coupling FC-NH$_2$ to linker-VHL**

To a solution of linker-VHL (0.0646 mmol, 1.1 eq) and HATU (0.0764 mmol, 1.3 eq) in dry DMF (4 mL) was added DIPEA (0.587 mmol, 10 eq). The resulting solution was stirred for 10 min and FC-NH$_2$ (0.0588 mmol, 1 eq) dissolved in dry DMF (1 mL) was added. The reaction mixture was then stirred for 16h and quenched by the addition of water (5 mL per mmol). The resulting mixture was concentrated under reduced pressure and purified by reversed-phase preparative HPLC to yield the desired product.

**Detailed synthetic procedures and characterization**

**FC-NH$_2$** - ((2*S*,3*R*,4*S*,5*S*,6*R*)-2-(((1*S*,4*R*,5*R*,6*R*,6a*S*,9*S*,10a*R*,*E*)-3-((*S*)-1-aminopropan-2-yl)-1,5-dihydroxy-9-(methoxymethyl)-6,10a-dimethyl-1,2,4,5,6,6a,7,8,9,10a-decahydrodicyclopenta[*a*,*d*][8]annulen-4-yl)oxy)-6-(((2-methylbut-3-en-2-yl)oxy)methyl)tetrahydro-2*H*-pyran-3,4,5-triol)

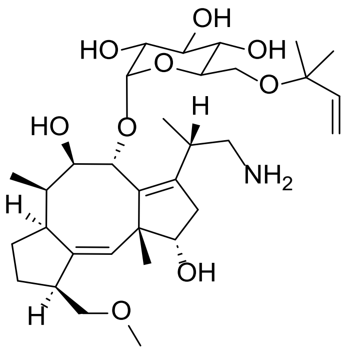

**FC-NH₂** was semi-synthesized from **FC-A** (obtained as a metabolite of *Phomopsis amygdali*) in accordance with the previous report, and isolated as white solid (469 mg, 0.735 mmol, 38% yield over deacetylation, tosylation, azide installation, and azide reduction) that matched the reported spectral data.[3] **LC-MS (ESI)**: $t_R$ = 3.03 min, *m/z* = 637.41 calcd. for [$C_{32}H_{53}NO_9$ +acetonitrile H]$^+$, found 637.75. **HRMS (ESI-TOF)**: *m/z* = 596.3799 calcd. for [$C_{32}H_{53}NO_9$ + H]$^+$, found 596.3805 [$\delta$ 1.0 ppm].

**FC-NAc** - (*N*-((*S*)-2-((1*S*,4*R*,5*R*,6*R*,6a*S*,9*S*,10a*R*,*E*)-1,5-dihydroxy-9-(methoxymethyl)-6,10a-dimethyl-4-(((2*S*,3R,4*S*,5*S*,6*R*)-3,4,5-trihydroxy-6-(((2-methylbut-3-en-2-yl)oxy)methyl)tetrahydro-2*H*-pyran-2-yl)oxy)-1,2,4,5,6,6a,7,8,9,10a-decahydrodicyclopenta[*a*,*d*][8]annulen-3-yl)propyl)acetamide)

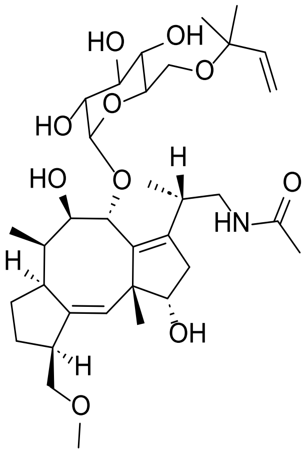

**FC-NAc** was synthesized via general procedure **A** using **FC-NH₂** (35.0 mg, 58.8 µmol, 1 eq), DIPEA (51.2 µL, 0.294 mmol, 5 eq) and acetic anhydride (5.55 µL, 58.8 µmol, 1 eq) in dry chloroform (3 mL). The crude product was purified by reversed-phase preparative HPLC (30-35% acetonitrile (+0.1% FA) in water (+0.1% FA) gradient) to afford **FC-NAc** as a white solid (28.6 mg, 44.9 µmol, 76.4% yield) that matched the reported spectral data.[3] **LC-MS (ESI)**: $t_R$ = 3.63 min, *m/z* = 638.39 calcd. for [$C_{34}H_{55}NO_{10}$ + H]$^+$, found 638.58. **HRMS (ESI-TOF)**: *m/z* = 638.3904 calcd. for [$C_{34}H_{55}NO_{10}$ + H]$^+$, found 638.3909 [$\delta$ 0.8 ppm]. **$^1$H NMR** (399 MHz, DMSO) $\delta$ 7.73 – 7.65 (m, 1H), 5.83 – 5.68 (m, 2H), 5.30 (s, 1H), 5.09 (dd, *J* = 10.9, 1.1 Hz, 2H), 4.86 – 4.70 (m, 5H), 3.76 – 3.62 (m, 3H), 3.61 – 3.54 (m, 1H), 3.49 – 3.40 (m, 2H), 3.28 – 3.13 (m, 8H), 3.07 – 2.91 (m, 3H), 2.69 – 2.60 (m, 2H), 2.13 – 2.04 (m, 1H), 1.92 – 1.77 (m, 6H), 1.51 – 1.37 (m, 3H), 1.17 (s, 6H), 1.13 (s, 3H), 0.92 (d, *J* = 5.9 Hz, 3H), 0.72 (d, *J* = 7.1 Hz, 3H). **$^{13}$C NMR** (100 MHz, DMSO) $\delta$ 171.3, 169.6, 144.1, 140.8, 139.4, 137.1, 127.0, 113.7, 102.3, 80.9, 77.9, 77.6, 75.3, 74.6, 73.1, 72.2, 70.6, 62.5, 57.9, 52.3, 48.2, 42.9, 41.0, 35.4, 35.2, 33.5, 27.1, 25.9, 25.7, 23.3, 22.6, 16.1, 9.2.

**2-methylbut-3-en-2-yl ether cleaved FC-NAc** - *N*-((*S*)-2-((1*S*,4*R*,5*R*,6*R*,6a*S*,9*S*,10a*R*,*E*)-1,5-dihydroxy-9-(methoxymethyl)-6,10a-dimethyl-4-(((2*S*,3*R*,4*S*,5*S*,6*R*)-3,4,5-trihydroxy-6-(hydroxymethyl)tetrahydro-2*H*-pyran-2-yl)oxy)-1,2,4,5,6,6a,7,8,9,10a-decahydrodicyclopenta[*a*,*d*][8]annulen-3-yl)propyl)acetamide

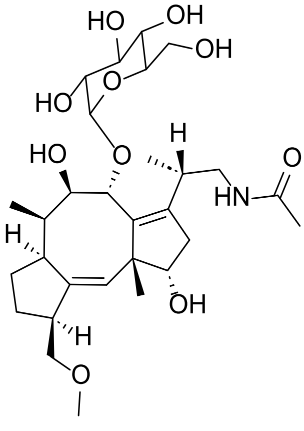

To a solution of **FC-NAc** (9.85 mg , 15.4 µmol, 1 eq) in dry DCM (1 mL) at 0° C was added TFA (59.1 µL, 772 µmol, 50 eq). The reaction mixture was stirred for 20 min at 0° C and quenched with 0.5 mL saturated $NaHCO_3$ solution. The resulting mixture was concentrated under reduced pressure and the crude material was purified by reversed-phase preparative HPLC (20-25% acetonitrile (+0.1% FA) in water (+0.1% FA) gradient) to afford **2-methylbut-3-en-2-yl ether cleaved FC-NAc** as a white solid (5.56 mg, 9.76 µmol, 63.2% yield). **LC-MS (ESI)**: $t_R$ = 3.01 min, *m/z* = 570.33 calcd. for $[C_{29}H_{47}NO_{10} + H]^+$, found 570.33. **HRMS (ESI-TOF)**: *m/z* = 570.3278 calcd. for $[C_{29}H_{47}NO_{10} + H]^+$, found 570.3267 [*δ* -1.9 ppm]. **$^1$H NMR** (399 MHz, MeOD) *δ* 5.37 (s, 1H), 4.93 (d, *J* = 3.9 Hz, 1H), 4.55 (s, 1H), 3.96 – 3.84 (m, 2H), 3.78 – 3.57 (m, 5H), 3.42 (dd, *J* = 9.7, 3.8 Hz, 1H), 3.34 – 3.29 (m, 6H), 3.26 – 3.14 (m, 3H), 2.79 – 2.70 (m, 2H), 2.25 (dd, *J* = 14.5, 6.8 Hz, 1H), 2.07 – 1.88 (m, 6H), 1.61 – 1.46 (m, 3H), 1.20 (s, 3H), 0.99 (d, *J* = 1.9 Hz, 3H), 0.81 (d, *J* = 7.2 Hz, 3H) ppm. **$^{13}$C NMR** (100 MHz, MeOD) *δ* 173.4, 143.0, 142.8, 139.1, 127.9, 103.7, 82.7, 79.7, 79.1, 77.1, 75.0, 74.3, 73.8, 71.5, 62.2, 58.8, 54.3, 49.8, 45.0, 43.0, 42.7, 36.7, 36.6, 35.0, 28.4, 24.3, 22.7, 16.2, 10.0 ppm. Experimental procedure adapted from Ohkanda *et al.*[4]

**VH032** - ((2*R*,4*S*)-1-((*R*)-2-acetamido-3,3-dimethylbutanoyl)-4-hydroxy-*N*-(4-(4-methylthiazol-5-yl)benzyl)pyrrolidine-2-carboxamide)

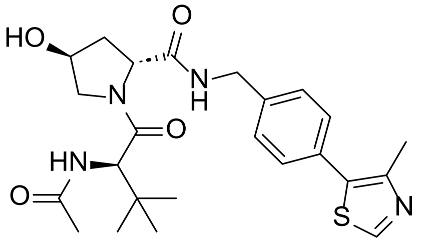

**VH032** was synthesized via general procedure **A** using VHL ligand 1 HCl salt (100 mg, 0.214 mmol, 1 eq), DIPEA (186 µL, 1.07 mmol, 5 eq) and acetic anhydride (20.2 µL, 0.214 mmol, 1 eq) in dry chloroform (5 mL). The crude product was purified by reversed-phase preparative HPLC (30-40% acetonitrile (+0.1% FA) in water (+0.1% FA) gradient) to afford **VH032** as a white solid (94.5 mg, 0.200 mmol, 93.4% yield) that matched the reported spectral data.[5] **LC-MS (ESI)**: $t_R$ = 3.42 min, *m/z* = 473.22 calcd. for $[C_{24}H_{32}N_4O_4S + H]^+$, found 473.17. **HRMS (ESI-TOF)**: *m/z* = 473.2223 calcd. for $[C_{24}H_{32}N_4O_4S + H]^+$, found 473.2219 [*δ* -0.8 ppm]. **$^1$H NMR** (399 MHz, $CDCl_3$) *δ* 8.68 (s, 1H), 7.41 – 7.27 (m, 5H), 6.16 (d, *J* = 8.7 Hz, 1H), 4.70 (t, *J* = 7.9 Hz, 1H), 4.59 – 4.47 (m, 3H), 4.33 (dd, *J* = 14.9, 5.2 Hz, 1H), 4.07 (d, *J* = 11.3 Hz, 1H), 3.61 (dd, *J* = 11.3, 3.7 Hz, 1H), 3.32 (s, 1H), 2.59 – 2.49 (m, 4H), 2.16 – 2.08 (m, 1H), 1.98 (s, 3H), 0.93 (s, 9H). **$^{13}$C NMR** (100 MHz, $CDCl_3$) *δ* 172.0, 170.9, 170.8, 150.5, 148.6, 138.2, 131.7, 131.2, 129.7, 128.3, 70.2, 58.6, 57.8, 56.9, 43.4, 36.0, 35.0, 26.5, 23.2, 16.2.

**CV1p** ((2*S*,4*R*)-1-((*S*)-2-(3-(3-(((*S*)-2-((1*S*,4*R*,5*R*,6*R*,6a*S*,9*S*,10a*R*,*E*)-1,5-dihydroxy-9-(methoxymethyl)-6,10a-dimethyl-4-(((2*S*,3*R*,4*S*,5*S*,6*R*)-3,4,5-trihydroxy-6-(((2-methylbut-3-en-2-yl)oxy)methyl)tetrahydro-2*H*-pyran-2-yl)oxy)-1,2,4,5,6,6a,7,8,9,10a-decahydrodicyclopenta[*a*,*d*][8]annulen-3-yl)propyl)amino)-3-oxopropoxy)propanamido)-3,3-dimethylbutanoyl)-4-hydroxy-*N*-(4-(4-methylthiazol-5-yl)benzyl)pyrrolidine-2-carboxamide)

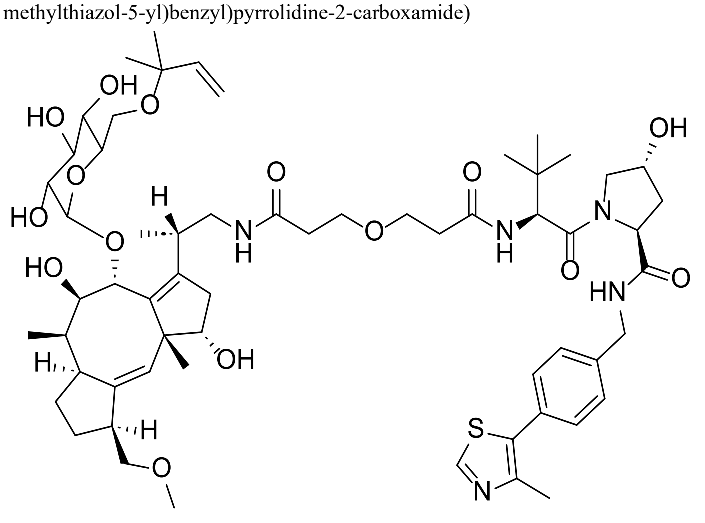

The intermediate **VHL1p** was synthesized via general procedure **B** using 3,3'-oxydipropionic acid (104 mg, 0.642 mmol, 3 eq), HATU (228 mg, 0.600 mmol, 2.8 eq) DIPEA (1.12 mL, 6.42 mmol, 30 eq) and VHL ligand 1 HCl salt (100 mg, 0.214 mmol, 1 eq) in dry DMF (5 mL). The crude product was purified by reversed-phase preparative HPLC (25-30% acetonitrile (+0.1% FA) in water (+0.1% FA) gradient) to afford intermediate **VHL1p** as a white solid (42.0 mg, 0.073 mmol, 34.2% yield). **LC-MS (ESI)**: $t_R$ = 3.41 min, *m/z* = 575.25 calcd. for $[C_{28}H_{38}N_4O_7S + H]^+$, found 575.92. **HRMS (ESI-TOF)**: *m/z* = 575.2540 calcd. for $[C_{28}H_{38}N_4O_7S + H]^+$, found 575.2531 [$\delta$ -1.6 ppm].

**CV1p** was synthesized via general procedure **C** using **VHL1p** (37.1 mg, 0.0646 mmol, 1.1 eq), HATU (29.0 mg, 0.0764 mmol, 1.3 eq), DIPEA (102 µL, 0.587 mmol, 10 eq) and **FC-NH$_2$** (35 mg, 0.0588 mmol, 1 eq) in dry DMF (5 mL). The crude product was purified by reversed-phase preparative HPLC (40-45% acetonitrile (+0.1% FA) in water (+0.1% FA) gradient) to afford **CV1p** as a white solid (6.08 mg, 5.28 µmol, 9.0% yield), 13% 2-methylbut-3-en-2-yl ether cleaved. **LC-MS (ESI)**: $t_R$ = 3.72 min, *m/z* = 1084.55 calcd. for $[C_{55}H_{81}N_5O_{15}S + H]^+$, found 1084.42; $t_R$ = 4.17 min, *m/z* = 1152.62 calcd. for $[C_{60}H_{89}N_5O_{15}S + H]^+$, found 1152.58. **HRMS (ESI-TOF)**: *m/z* = 542.7803 calcd. for $[C_{55}H_{81}N_5O_{15}S + 2H]^{2+}$, found 542.7787 [$\delta$ -2.9 ppm]; *m/z* = 576.8116 calcd. for $[C_{60}H_{89}N_5O_{15}S + 2H]^{2+}$, found 576.8120 [$\delta$ 0.7 ppm]. **$^1$H NMR** (399 MHz, DMSO) $\delta$ 9.03 (s, 1H), 8.56 (t, *J* = 6.1 Hz, 1H), 7.93 (d, *J* = 9.3 Hz, 1H), 7.77 – 7.64 (m, 1H), 7.43 – 7.37 (m, 4H), 5.77 (dd, *J* = 17.7, 10.8 Hz, 1H), 5.32 – 5.27 (m, 1H), 5.11 (dd, *J* = 17.7, 1.5 Hz, 1H), 5.07 (dd, *J* = 10.8, 1.5 Hz, 1H), 4.74 – 4.73 (m, 1H), 4.55 (d, *J* = 9.4 Hz, 1H), 4.44 – 4.33 (m, 3H), 4.22 (dd, *J* = 15.9, 5.3 Hz, 1H), 3.68 – 3.42 (m, 12H), 3.25 – 2.97 (m, 11H), 2.68 – 2.59 (m, 2H), 2.45 (s, 3H), 2.41 – 2.31 (m, 3H), 2.11 – 2.01 (m, 2H), 1.93 – 1.80 (m, 4H), 1.47 – 1.37 (m, 3H), 1.29 – 0.97 (m, 10H), 0.94 – 0.88 (m, 12H), 0.72 (d, *J* = 7.0 Hz, 3H) ppm. **$^{13}$C NMR** (100 MHz, DMSO) $\delta$ 171.9, 170.4, 169.9, 169.5, 151.6, 147.4, 144.1, 140.8, 139.6, 139.4, 137.0, 131.3, 129.5, 128.6, 127.4, 127.1, 113.6, 102.2, 80.8, 77.8, 77.6, 75.3, 74.6, 73.0, 72.1, 72.1, 70.5, 68.9, 66.6, 62.5, 58.7, 57.9, 56.4, 56.3, 52.3, 48.2, 42.7, 41.6, 41.0, 40.9, 37.9, 36.0, 35.5, 35.4, 35.3, 35.2, 33.5, 27.1, 26.3, 25.8, 25.7, 23.3, 15.8, 15.8, 9.2 ppm.

**CV2p** ((2*S*,4*R*)-1-((2*S*,19*S*)-2-(tert-butyl)-19-((1*S*,4*R*,5*R*,6*R*,6a*S*,9*S*,10a*R*,*E*)-1,5-dihydroxy-9-(methoxymethyl)-6,10a-dimethyl-4-(((2*S*,3*R*,4*S*,5*S*,6*R*)-3,4,5-trihydroxy-6-(((2-methylbut-3-en-2-yl)oxy)methyl)tetrahydro-2*H*-pyran-2-yl)oxy)-1,2,4,5,6,6a,7,8,9,10a-decahydrodicyclopenta[*a*,*d*][8]annulen-3-yl)-4,16-dioxo-7,10,13-trioxa-3,17-diazaicosanoyl)-4-hydroxy-*N*-(4-(4-methylthiazol-5-yl)benzyl)pyrrolidine-2-carboxamide)

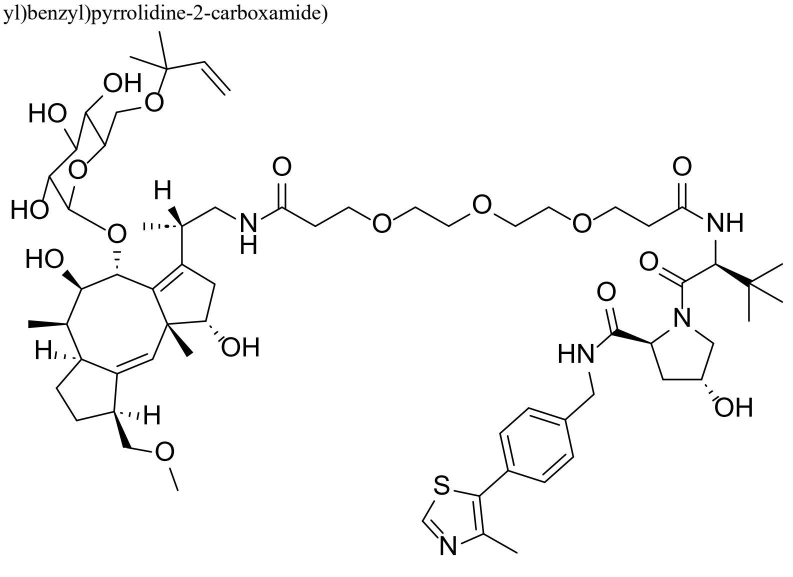

The intermediate **VHL2p** was synthesized via general procedure **B** using 3,3'-((oxybis(ethane-2,1-diyl))bis(oxy))dipropionic acid (161 mg, 0.642 mmol, 3 eq), HATU (228 mg, 0.600 mmol, 2.8 eq) DIPEA (1.12 mL, 6.42 mmol, 30 eq) and VHL ligand 1 HCl salt (100 mg, 0.214 mmol, 1 eq) in dry DMF (5 mL). The crude product was purified by reversed-phase preparative HPLC (35-40% acetonitrile (+0.1% FA) in water (+0.1% FA) gradient) to afford intermediate **VHL2p** as a white solid (86.6 mg, 0.131 mmol, 61.0% yield).
**LC-MS (ESI)**: $t_R$ = 3.50 min, *m/z* = 663.31 calcd. for $[C_{32}H_{46}N_4O_9S + H]^+$, found 663.25. **HRMS (ESI-TOF)**: *m/z* = 663.3064 calcd. for $[C_{32}H_{46}N_4O_9S + H]^+$, found 663.3063 [$\delta$ -0.2 ppm].

**CV2p** was synthesized via general procedure **C** using **VHL2p** (42.8 mg, 0.0646 mmol, 1.1 eq), HATU (29.0 mg, 0.0764 mmol, 1.3 eq), DIPEA (102 µL, 0.587 mmol, 10 eq) and **FC-NH₂** (35 mg, 0.0588 mmol, 1 eq) in dry DMF (5 mL). The crude product was purified by reversed-phase preparative HPLC (45-50% acetonitrile (+0.1% FA) in water (+0.1% FA) gradient) to afford **CV2p** as a white solid (11.9 mg, 9.58 µmol, 16.3% yield), 5% 2-methylbut-3-en-2-yl ether cleaved. **LC-MS (ESI)**: $t_R$ = 3.72 min, *m/z* = 1172.61 calcd. for $[C_{59}H_{89}N_5O_{17}S + H]^+$, found 1172.33; $t_R$ = 4.15 min, *m/z* = 1240.67 calcd. for $[C_{64}H_{97}N_5O_{17}S + H]^+$, found 1240.50. **HRMS (ESI-TOF)**: *m/z* = 586.8065 calcd. for $[C_{59}H_{89}N_5O_{17}S + 2H]^{2+}$, found 586.8062 [$\delta$ -0.5 ppm]; *m/z* = 620.8378 calcd. for $[C_{64}H_{97}N_5O_{17}S + 2H]^{2+}$, found 620.8384 [$\delta$ 1.0 ppm]. **$^1$H NMR** (399 MHz, $CD_3CN$) $\delta$ 9.25 (s, 1H), 7.50 – 7.40 (m, 5H), 7.03 (d, *J* = 8.8 Hz, 1H), 6.99 – 6.92 (m, 1H), 5.81 (dd, *J* = 17.7, 10.8 Hz, 1H), 5.32 – 5.29 (m, 1H), 5.14 (dd, *J* = 17.7, 1.4 Hz, 1H), 5.08 (dd, *J* = 10.8, 1.4 Hz, 1H), 4.87 (d, *J* = 3.8 Hz, 1H), 4.58 – 4.42 (m, 4H), 4.30 (dd, *J* = 15.8, 5.6 Hz, 1H), 3.81 – 3.54 (m, 21H), 3.38 – 3.24 (m, 9H), 3.10 – 3.05 (m, 2H), 2.81 – 2.67 (m, 4H), 2.50 – 2.39 (m, 4H), 2.20 – 2.02 (m, 3H), 1.90 – 1.85 (m, 2H), 1.52 – 1.46 (m, 3H), 1.23 – 1.17 (m, 10H), 1.02 – 0.93 (m, 12H), 0.78 (d, *J* = 7.1 Hz, 3H) ppm. **$^{13}$C NMR** (100 MHz, $CD_3CN$) $\delta$ 173.0, 172.7, 172.2, 171.8, 154.3, 145.2, 144.9, 143.1, 141.9, 141.9, 138.7, 135.3, 130.1, 128.9, 127.3, 114.2, 103.2, 82.0, 79.4, 79.1, 76.5, 76.0, 74.4, 73.7, 73.4, 72.1, 70.9, 70.8, 70.6, 67.9, 67.7, 63.6, 60.2, 58.6, 58.2, 57.7, 54.2, 49.5, 44.8, 43.1, 42.4, 42.2, 38.4, 37.1, 37.0, 36.7, 36.4, 36.2, 34.8, 28.3, 26.7, 26.3, 24.3, 16.6, 14.6, 10.0 ppm.

**CV3p** ($N^1$-((*S*)-2-((1*S*,4*R*,5*R*,6*R*,6a*S*,9*S*,10a*R*,*E*)-1,5-dihydroxy-9-(methoxymethyl)-6,10a-dimethyl-4-(((2*S*,3*R*,4*S*,5*S*,6*R*)-3,4,5-trihydroxy-6-(((2-methylbut-3-en-2-yl)oxy)methyl)tetrahydro-2*H*-pyran-2-yl)oxy)-1,2,4,5,6,6a,7,8,9,10a-decahydrodicyclopenta[*a*,*d*][8]annulen-3-yl)propyl)-$N^{19}$-((*S*)-1-((2*S*,4*R*)-4-hydroxy-2-((4-(4-methylthiazol-5-yl)benzyl)carbamoyl)pyrrolidin-1-yl)-3,3-dimethyl-1-oxobutan-2-yl)-4,7,10,13,16-pentaoxanonadecanediamide)

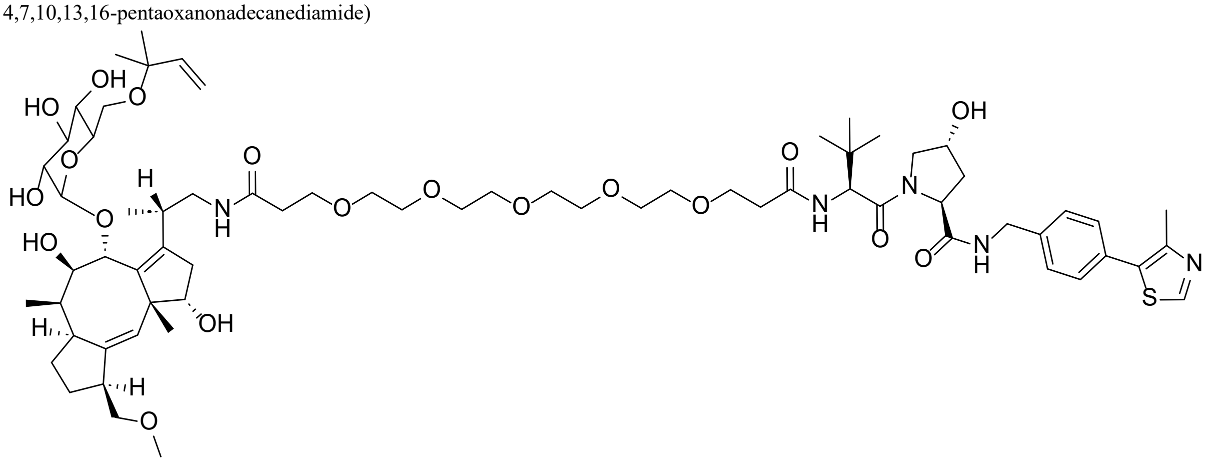

The intermediate **VHL3p** was synthesized via general procedure **B** using 4,7,10,13,16-pentaoxanonadecanedioic acid (217 mg, 0.642 mmol, 3 eq), HATU (228 mg, 0.600 mmol, 2.8 eq) DIPEA (1.12 mL, 6.42 mmol, 30 eq),and VHL ligand 1 HCl salt (100 mg, 0.214 mmol, 1 eq) in dry DMF (5 mL). The crude product was purified by reversed-phase preparative HPLC (35-40% acetonitrile (+0.1% FA) in water (+0.1% FA) gradient) to afford intermediate **VHL3p** as a white solid (90.5 mg, 0.121 mmol, 56.3% yield). **LC-MS (ESI)**: $t_R$ = 3.55 min, *m/z* = 751.36 calcd. for $[C_{36}H_{54}N_4O_{11}S + H]^+$, found 751.42. **HRMS (ESI-TOF)**: *m/z* = 751.3588 calcd. for $[C_{36}H_{54}N_4O_{11}S + H]^+$, found 751.3586 [$\delta$ -0.3 ppm].

**CV3p** was synthesized via general procedure **C** using **VHL3p** (48.5 mg, 0.0646 mmol, 1.1 eq), HATU (29.0 mg, 0.0764 mmol, 1.3 eq), DIPEA (102 µL, 0.587 mmol, 10 eq) and **FC-NH₂** (35 mg, 0.0588 mmol, 1 eq) in dry DMF (5 mL). The crude product was purified by reversed-phase preparative HPLC (45-50% acetonitrile (+0.1% FA) in water (+0.1% FA) gradient) to afford **CV3p** as a white solid (22.2 mg, 16.7 µmol, 28.4% yield), 6% 2-methylbut-3-en-2-yl ether cleaved. **LC-MS (ESI)**: $t_R$ = 3.74 min, *m/z* = 1260.66 calcd. for $[C_{63}H_{97}N_5O_{19}S + H]^+$, found 1260.42; $t_R$ = 4.16 min, *m/z* = 1328.72 calcd. for $[C_{68}H_{105}N_5O_{19}S + H]^+$, found 1328.58. **HRMS (ESI-TOF)**: *m/z* = 630.8328 calcd. for $[C_{63}H_{97}N_5O_{19}S + 2H]^{2+}$, found 630.8316 [$\delta$ -1.9 ppm]; *m/z* = 664.8641 calcd. for $[C_{68}H_{105}N_5O_{19}S + 2H]^{2+}$, found 664.8644 [$\delta$ 0.5 ppm]. **$^1$H NMR** (399 MHz, $CD_3CN$) $\delta$ 9.24 (s, 1H), 7.50 – 7.43 (m, 4H), 7.33 (t, *J* = 6.2 Hz, 1H), 7.02 (d, *J* = 9.0 Hz, 1H), 6.98 – 6.92 (m, 1H), 5.81 (dd, *J* = 17.7, 10.8 Hz, 1H), 5.31 (s, 1H), 5.14 (dd, *J* = 17.7, 1.4 Hz, 1H), 5.08 (dd, *J* = 10.8, 1.4 Hz, 1H), 4.87 (d, *J* = 4.0 Hz, 1H), 4.57 – 4.42 (m, 4H), 4.29 (dd, *J* = 15.8, 5.5 Hz, 1H), 3.85 – 3.53 (m, 31H), 3.40 – 3.24 (m, 8H), 3.09 – 3.05 (m, 2H), 2.84 – 2.70 (m, 3H), 2.52 – 2.38 (m, 4H), 2.24 – 2.03 (m, 3H), 1.91 – 1.84 (m, 2H), 1.53 – 1.48 (m, 3H), 1.23 – 1.16 (m, 10H), 1.02 – 0.92 (m, 12H), 0.79 (d, *J* = 7.2 Hz, 3H) ppm. **$^{13}$C NMR** (100 MHz, $CD_3CN$) $\delta$ 172.9, 172.7, 172.1, 171.7, 154.2, 145.2, 145.2, 143.1, 141.9, 141.7, 138.7, 135.1, 130.1, 129.1, 128.9, 127.3, 114.2, 103.2, 82.0, 79.3, 79.1, 76.5, 76.0, 74.5, 73.6, 73.4, 72.1, 71.0, 70.9, 70.8, 70.7, 67.9, 67.8, 63.6, 60.2, 58.6, 58.1, 57.6, 54.2, 49.5, 44.8, 43.1, 42.4, 42.2, 38.4, 37.2, 37.0, 36.7, 36.4, 36.2, 34.7, 28.3, 26.8, 26.3, 24.3, 16.6, 14.8, 10.0 ppm.

**CV4p** ($N^1$-((*S*)-2-((1*S*,4*R*,5*R*,6*R*,6a*S*,9*S*,10a*R*,*E*)-1,5-dihydroxy-9-(methoxymethyl)-6,10a-dimethyl-4-(((2*S*,3*R*,4*S*,5*S*,6*R*)-3,4,5-trihydroxy-6-(((2-methylbut-3-en-2-yl)oxy)methyl)tetrahydro-2*H*-pyran-2-yl)oxy)-1,2,4,5,6,6a,7,8,9,10a-decahydrodicyclopenta[*a*,*d*][8]annulen-3-yl)propyl)-$N^{25}$-((*S*)-1-((2*S*,4*R*)-4-hydroxy-2-((4-(4-methylthiazol-5-yl)benzyl)carbamoyl)pyrrolidin-1-yl)-3,3-dimethyl-1-oxobutan-2-yl)-4,7,10,13,16,19,22-heptaoxapentacosanediamide)

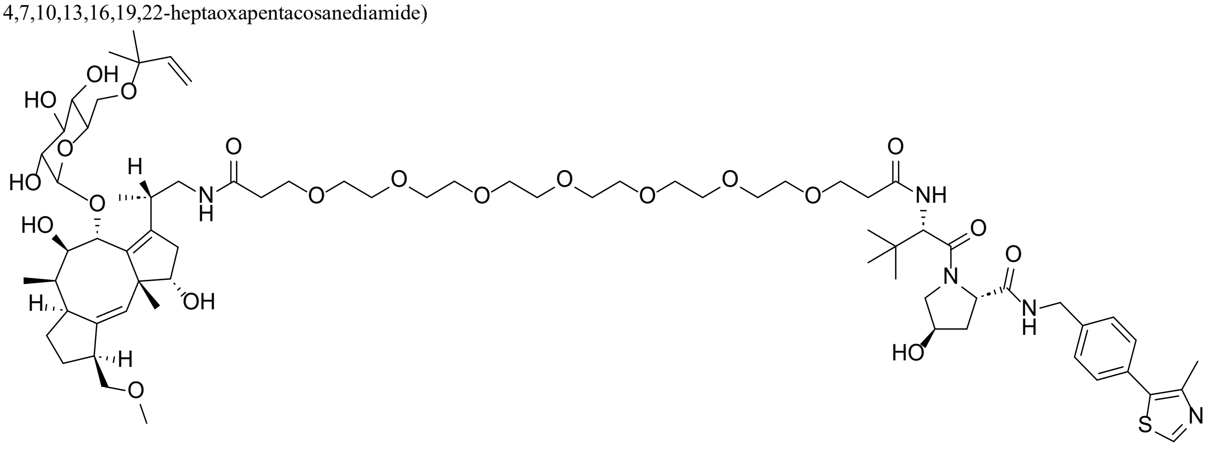

The intermediate **VHL4p** was synthesized via general procedure **B** using 4,7,10,13,16,19,22-heptaoxapentacosanedioic acid (274 mg, 0.642 mmol, 3 eq), HATU (228 mg, 0.600 mmol, 2.8 eq) DIPEA (1.12 mL, 6.42 mmol, 30 eq) and VHL ligand 1 HCl salt (100 mg, 0.214 mmol, 1 eq) in dry DMF (5 mL). The crude product was purified by reversed-phase preparative HPLC (35-40% acetonitrile (+0.1% FA) in water (+0.1% FA) gradient) to afford intermediate **VHL4p** as a white solid (58.0 mg, 0.0692 mmol, 32.3% yield). **LC-MS (ESI)**: $t_R$ = 3.59 min, *m/z* = 839.41 calcd. for $[C_{40}H_{62}N_4O_{13}S + H]^+$, found 839.50. **HRMS (ESI-TOF)**: *m/z* = 839.4113 calcd. for $[C_{40}H_{62}N_4O_{13}S + H]^+$, found 839.4113 [$\delta$ 0.0 ppm].

**CV4p** was synthesized via general procedure **C** using **VHL4p** (54.2 mg, 0.0646 mmol, 1.1 eq), HATU (29.0 mg, 0.0764 mmol, 1.3 eq), DIPEA (102 µL, 0.587 mmol, 10 eq) and **FC-NH₂** (35 mg, 0.0588 mmol, 1 eq) in dry DMF (5 mL). The crude product was purified by reversed-phase preparative HPLC (45-50% acetonitrile (+0.1% FA) in water (+0.1% FA) gradient) to afford **CV4p** as a white solid (26.1 mg, 18.5 µmol, 31.4% yield), 5% 2-methylbut-3-en-2-yl ether cleaved. **LC-MS (ESI)**: $t_R$ = 3.77 min, *m/z* = 1348.71 calcd. for $[C_{67}H_{105}N_5O_{21}S + H]^+$, found 1348.42; $t_R$ = 4.16 min, *m/z* = 1416.77 calcd. for $[C_{72}H_{113}N_5O_{21}S + H]^+$, found 1416.58. **HRMS (ESI-TOF)**: *m/z* = 674.8589 calcd. for $[C_{67}H_{105}N_5O_{21}S + 2H]^{2+}$, found 674.8579 [$\delta$ -1.5 ppm]; *m/z* = 708.8903 calcd. for $[C_{72}H_{113}N_5O_{21}S + 2H]^{2+}$, found 708.8897 [$\delta$ -0.8 ppm]. **$^{1}$H NMR** (399 MHz, $CD_3CN$) $\delta$ 9.28 (s, 1H), 7.51 – 7.44 (m, 4H), 7.31 (t, *J* = 6.2 Hz, 1H), 7.01 (d, *J* = 9.0 Hz, 1H), 6.97 – 6.89 (m, 1H), 5.81 (dd, *J* = 17.7, 10.8 Hz, 1H), 5.32 – 5.29 (m, 1H), 5.14 (dd, *J* = 17.7, 1.4 Hz, 1H), 5.08 (dd, *J* = 10.8, 1.4 Hz, 1H), 4.87 (d, *J* = 3.8 Hz, 1H), 4.59 – 4.41 (m, 4H), 4.28 (dd, *J* = 15.8, 5.5 Hz, 1H), 3.82 – 3.52 (m, 39H), 3.42 – 3.21 (m, 8H), 3.10 – 3.04 (m, 2H), 2.80 – 2.72 (m, 3H), 2.52 – 2.42 (m, 4H), 2.22 – 2.03 (m, 3H), 1.92 – 1.85 (m, 2H), 1.54 – 1.47 (m, 3H), 1.24 – 1.15 (m, 10H), 1.02 – 0.94 (m, 12H), 0.79 (d, *J* = 7.2 Hz, 3H) ppm. **$^{13}$C NMR** (100 MHz, $CD_3CN$) $\delta$ 173.0, 172.6, 172.0, 171.7, 154.4, 145.2, 145.1, 143.1, 141.9, 141.8, 138.8, 135.1, 130.1, 129.0, 128.9, 127.3, 114.2, 103.2, 82.0, 79.3, 79.0, 76.5, 76.0, 74.4, 73.6, 73.4, 72.1, 71.0, 71.0, 71.0, 70.9, 70.9, 70.8, 70.6, 67.9, 67.8, 63.6, 60.2, 58.6, 58.1, 57.6, 54.2, 49.5, 44.8, 43.1, 42.4, 42.2, 38.4, 37.2, 37.0, 36.7, 36.4, 36.2, 34.7, 28.3, 26.7, 26.3, 26.3, 24.3, 16.6, 14.7, 10.0 ppm.

**CV0a** ($N^1$-((*S*)-2-((1*S*,4*R*,5*R*,6*R*,6a*S*,9*S*,10a*R*,*E*)-1,5-dihydroxy-9-(methoxymethyl)-6,10a-dimethyl-4-(((2*S*,3*R*,4*S*,5*S*,6*R*)-3,4,5-trihydroxy-6-(((2-methylbut-3-en-2-yl)oxy)methyl)tetrahydro-2*H*-pyran-2-yl)oxy)-1,2,4,5,6,6a,7,8,9,10a-decahydrodicyclopenta[*a*,*d*][8]annulen-3-yl)propyl)-$N^4$-((*S*)-1-((2*S*,4*R*)-4-hydroxy-2-((4-(4-methylthiazol-5-yl)benzyl)carbamoyl)pyrrolidin-1-yl)-3,3-dimethyl-1-oxobutan-2-yl)succinamide)

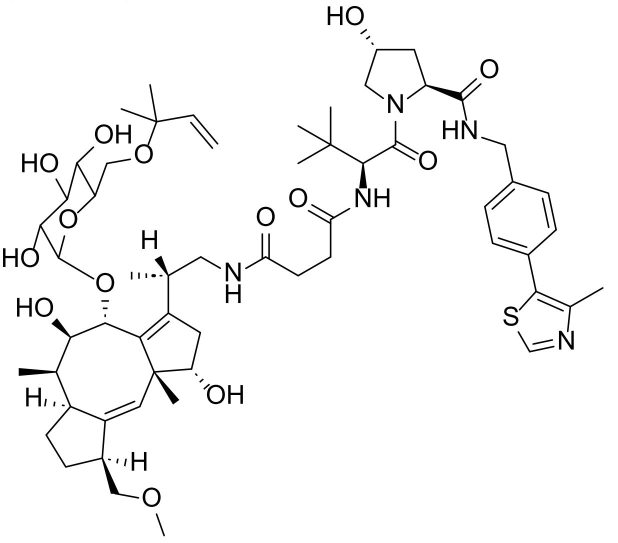

To a solution of VHL ligand 1 HCl salt (100mg, 0.214 mmol, 1 eq) and DIPEA (186 µL, 1.07 mmol, 5 eq) in dry chloroform (5 mL) was added succinic anhydride (25.7 mg, 0.257 mmol, 1.2 eq). The resulting solution was stirred for 3 h and subsequently concentrated under reduced pressure. The crude product was purified by reversed-phase preparative HPLC (30-35% acetonitrile (+0.1% FA) in water (+0.1% FA) gradient) to afford intermediate **VHL0a** as a white solid (77.5 mg, 0.146 mmol, 68.2% yield). **LC-MS (ESI)**: $t_R$ = 3.38 min, *m/z* = 531.23 calcd. for $[C_{26}H_{34}N_4O_6S + H]^+$, found 531.25. **HRMS (ESI-TOF)**: *m/z* = 531.2277 calcd. for $[C_{26}H_{34}N_4O_6S + H]^+$, found 531.2272 [$\delta$ -0.9 ppm].

**CV0a** was synthesized via general procedure **C** using **VHL0a** (34.3 mg, 0.0646 mmol, 1.1 eq), HATU (29.0 mg, 0.0764 mmol, 1.3 eq), DIPEA (102 µL, 0.587 mmol, 10 eq) and **FC-NH$_2$** (35 mg, 0.0588 mmol, 1 eq) in dry DMF (5 mL). The crude product was purified by reversed-phase preparative HPLC (40-45% acetonitrile (+0.1% FA) in water (+0.1% FA) gradient) to afford **CV0a** as a white solid (8.46 mg, 7.64 µmol, 13.0% yield), 43% 2-methylbut-3-en-2-yl ether cleaved. **LC-MS (ESI)**: $t_R$ = 3.73 min, *m/z* = 1040.53 calcd. for $[C_{53}H_{77}N_5O_{14}S + H]^+$, found 1040.33; $t_R$ = 4.19 min, *m/z* = 1108.59 calcd. for $[C_{58}H_{85}N_5O_{14}S + H]^+$, found 1108.33. **HRMS (ESI-TOF)**: *m/z* = 520.7672 calcd. for $[C_{53}H_{77}N_5O_{14}S + 2H]^{2+}$, found 520.7668 [$\delta$ -0.8 ppm], *m/z* = 554.7985 calcd. for $[C_{58}H_{85}N_5O_{14}S + 2H]^{2+}$, found 554.7987 [$\delta$ 0.4 ppm]. **$^1$H NMR** (399 MHz, $CD_3CN$) $\delta$ 9.50 (s, 1H), 7.50 – 7.44 (m, 4H), 7.01 – 6.97 (m, 1H), 5.82 (dd, *J* = 17.7, 10.8 Hz, 1H), 5.33 – 5.26 (m, 1H), 5.14 (dd, *J* = 17.7, 1.4 Hz, 1H), 5.08 (dd, *J* = 10.8, 1.4 Hz, 1H), 4.93 – 4.84 (m, 1H), 4.60 – 4.29 (m, 5H), 4.00 – 3.91 (m, 1H), 3.84 – 3.62 (m, 6H), 3.49 – 3.37 (m, 3H), 3.30 – 3.23 (m, 5H), 3.18 – 3.08 (m, 2H), 2.87 – 2.75 (m, 1H), 2.54 – 2.47 (m, 9H), 2.22 – 2.13 (m, 2H), 2.11 – 1.98 (m, 2H), 1.93 – 1.81 (m, 3H), 1.74 – 1.62 (m, 1H), 1.54 – 1.42 (m, 3H), 1.29 – 1.10 (m, 8H), 0.99 (d, *J* = 8.4 Hz, 12H), 0.78 (d, *J* = 2.6 Hz, 3H) ppm. **$^{13}$C NMR** (100 MHz, $CD_3CN$) $\delta$ 173.7, 173.6, 172.7, 171.6, 155.2, 144.8, 142.8, 142.2, 141.6, 138.4, 136.5, 129.8, 128.6, 127.2, 127.0, 113.8, 102.7, 81.5, 78.8, 78.6, 76.2, 75.7, 74.2, 73.0, 72.8, 72.1, 70.7, 63.0, 59.9, 58.6, 58.3, 57.5, 53.9, 49.1, 44.8, 42.7, 42.0, 41.8, 38.0, 36.2, 36.0, 35.5, 33.9, 30.7, 30.6, 27.9, 26.4, 25.9, 25.9, 23.9, 16.0, 13.2, 9.6 ppm.

**CV1a** ($N^1$-((*S*)-2-((1*S*,4*R*,5*R*,6*R*,6a*S*,9*S*,10a*R*,*E*)-1,5-dihydroxy-9-(methoxymethyl)-6,10a-dimethyl-4-(((2*S*,3*R*,4*S*,5*S*,6*R*)-3,4,5-trihydroxy-6-(((2-methylbut-3-en-2-yl)oxy)methyl)tetrahydro-2*H*-pyran-2-yl)oxy)-1,2,4,5,6,6a,7,8,9,10a-decahydrodicyclopenta[*a*,*d*][8]annulen-3-yl)propyl)-$N^7$-((*S*)-1-((2*S*,4*R*)-4-hydroxy-2-((4-(4-methylthiazol-5-yl)benzyl)carbamoyl)pyrrolidin-1-yl)-3,3-dimethyl-1-oxobutan-2-yl)heptanediamide)

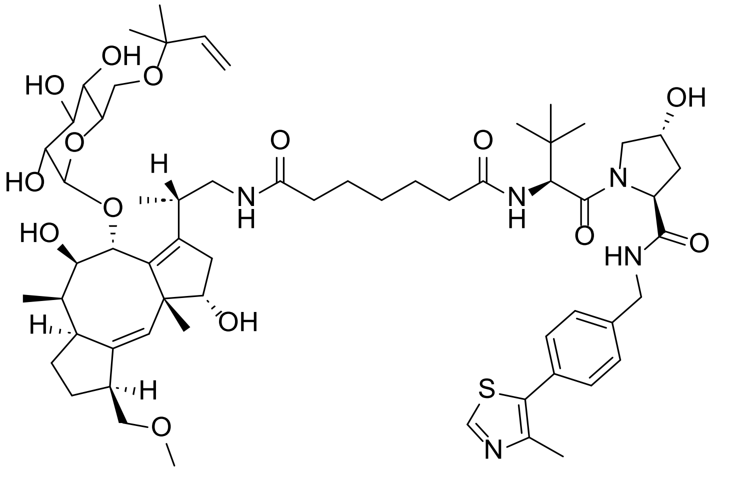

The intermediate **VHL1a** was synthesized via general procedure **B** using heptanedioic acid (171 mg, 1.07 mmol, 5 eq), HATU (366 mg, 0.964 mmol, 4.5 eq) DIPEA (1.12 mL, 6.42 mmol, 30 eq) and VHL ligand 1 HCl salt (100 mg, 0.214 mmol, 1 eq) in dry DMF (10 mL). The crude product was purified by reversed-phase preparative HPLC (30-35% acetonitrile (+0.1% FA) in water (+0.1% FA) gradient) to afford intermediate **VHL1a** as a white solid (50.5 mg, 0.0882 mmol, 41.2% yield). **LC-MS (ESI)**: $t_R$ = 3.58 min, *m/z* = 573.27 calcd. for $[C_{29}H_{40}N_4O_6S + H]^+$, found 573.42. **HRMS (ESI-TOF)**: *m/z* = 573.2747 calcd. for $[C_{29}H_{40}N_4O_6S + H]^+$, found 573.2733 [$\delta$ -2.4 ppm].

**CV1a** was synthesized via general procedure **C** using **VHL1a** (37.0 mg, 0.0646 mmol, 1.1 eq), HATU (29.0 mg, 0.0764 mmol, 1.3 eq), DIPEA (102 µL, 0.587 mmol, 10 eq) and **FC-NH$_2$** (35 mg, 0.0588 mmol, 1 eq) in dry DMF (5 mL). The crude product was purified by reversed-phase preparative HPLC (40-45% acetonitrile (+0.1% FA) in water (+0.1% FA) gradient) to afford **CV1a** as a white solid (8.21 mg, 7.14 µmol, 12.2% yield), 7% 2-methylbut-3-en-2-yl ether cleaved. **LC-MS (ESI)**: $t_R$ = 3.79 min, *m/z* = 1082.57 calcd. for $[C_{56}H_{83}N_5O_{14}S + H]^+$, found 1082.42; $t_R$ = 4.24 min, *m/z* = 1150.64 calcd. for $[C_{61}H_{91}N_5O_{14}S + H]^+$, found 1150.58. **HRMS (ESI-TOF)**: *m/z* = 541.7907 calcd. for $[C_{56}H_{83}N_5O_{14}S + 2H]^{2+}$, found 541.7891 [$\delta$ -3.0 ppm]; *m/z* = 575.8220 calcd. for $[C_{61}H_{91}N_5O_{14}S + 2H]^{2+}$, found 575.8218 [$\delta$ -0.3 ppm]. **$^1$H NMR** (399 MHz, DMSO) $\delta$ 9.01 (s, 1H), 8.56 (t, *J* = 6.1 Hz, 1H), 7.83 (d, *J* = 9.3 Hz, 1H), 7.63 – 7.56 (m, 1H), 7.43 – 7.37 (m, 4H), 5.76 (dd, *J* = 17.7, 10.8 Hz, 1H), 5.34 – 5.26 (m, 1H), 5.11 (dd, *J* = 17.8, 1.5 Hz, 1H), 5.06 (dd, *J* = 10.8, 1.4 Hz, 1H), 4.73 (d, *J* = 3.8 Hz, 1H), 4.54 (d, *J* = 9.3 Hz, 1H), 4.46 – 4.40 (m, 2H), 4.36 – 4.33 (m, 1H), 4.21 (dd, *J* = 16.1, 5.3 Hz, 1H), 3.45 – 3.42 (m, 2H), 3.31 – 3.10 (m, 10H), 3.06 – 2.95 (m, 3H), 2.66 – 2.59 (m, 2H), 2.45 – 2.44 (m, 3H), 2.28 – 2.21 (m, 1H), 2.15 – 1.98 (m, 6H), 1.93 – 1.80 (m, 4H), 1.55 – 1.35 (m, 8H), 1.32 – 1.00 (m, 13H), 0.94 – 0.87 (m, 12H), 0.72 (d, *J* = 7.0 Hz, 3H) ppm. **$^{13}$C NMR** (100 MHz, DMSO) $\delta$ 172.6, 172.1, 172.0, 169.7, 151.6, 147.5, 144.2, 140.9, 139.6, 139.5, 137.1, 131.3, 129.5, 128.7, 127.4, 127.1, 113.7, 102.2, 80.8, 77.9, 77.6, 75.3, 74.6, 73.1, 72.2, 72.1, 70.5, 68.9, 62.5, 58.7, 57.9, 56.4, 56.3, 52.4, 48.2, 42.7, 41.7, 41.0, 38.0, 35.4, 35.3, 35.2, 35.2, 34.9, 33.5, 28.5, 27.1, 26.4, 25.9, 25.7, 25.3, 25.2, 23.3, 16.0, 15.8, 9.2 ppm.

**CV2a** (*N*[1]-((*S*)-2-((1*S*,4*R*,5*R*,6*R*,6a*S*,9*S*,10a*R*,*E*)-1,5-dihydroxy-9-(methoxymethyl)-6,10a-dimethyl-4-(((2*S*,3*R*,4*S*,5*S*,6*R*)-3,4,5-trihydroxy-6-(((2-methylbut-3-en-2-yl)oxy)methyl)tetrahydro-2*H*-pyran-2-yl)oxy)-1,2,4,5,6,6a,7,8,9,10a-decahydrodicyclopenta[*a*,*d*][8]annulen-3-yl)propyl)-*N*[13]-((*S*)-1-((2*S*,4*R*)-4-hydroxy-2-((4-(4-methylthiazol-5-yl)benzyl)carbamoyl)pyrrolidin-1-yl)-3,3-dimethyl-1-oxobutan-2-yl)tridecanediamide)

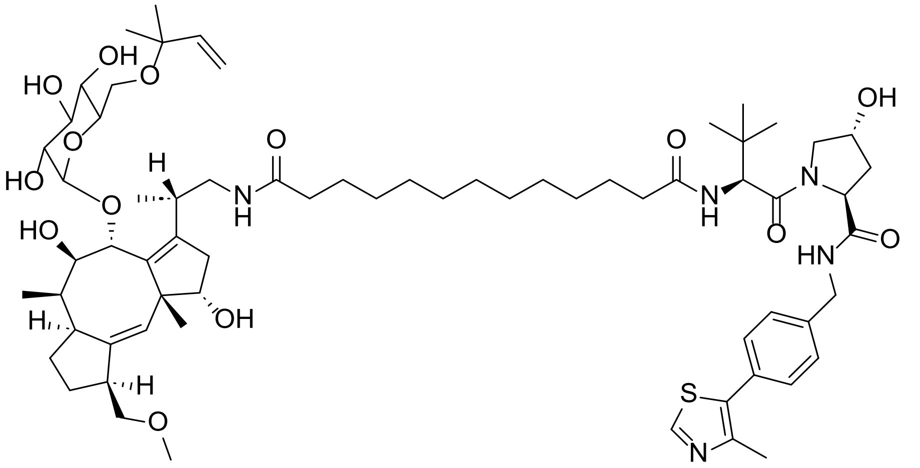

The intermediate **VHL2a** was synthesized via general procedure **B** using tridecanedioic acid (262 mg, 1.07 mmol, 5 eq), HATU (366 mg, 0.964 mmol, 4.5 eq) DIPEA (1.12 mL, 6.42 mmol, 30 eq) and VHL ligand 1 HCl salt (100 mg, 0.214 mmol, 1 eq) in dry DMF (15 mL). The crude product was purified by reversed-phase preparative HPLC (45-50% acetonitrile (+0.1% FA) in water (+0.1% FA) gradient) to afford intermediate **VHL2a** as a white solid (54.2 mg, 0.0825 mmol, 38.5% yield). **LC-MS (ESI)**: $t_R$ = 4.44 min, *m/z* = 657.37 calcd. for $[C_{35}H_{52}N_4O_6S + H]^+$, found 657.50. **HRMS (ESI-TOF)**: *m/z* = 657.3686 calcd. for $[C_{35}H_{52}N_4O_6S + H]^+$, found 657.3676 [$\delta$ -1.5 ppm].

**CV2a** was synthesized via general procedure **C** using **VHL2a** (42.5 mg, 0.0646 mmol, 1.1 eq), HATU (29.0 mg, 0.0764 mmol, 1.3 eq), DIPEA (102 µL, 0.587 mmol, 10 eq) and **FC-NH$_2$** (35 mg, 0.0588 mmol, 1 eq) in dry DMF (5 mL). The crude product was purified by reversed-phase preparative HPLC (50-55% acetonitrile (+0.1% FA) in water (+0.1% FA) gradient) to afford **CV2a** as a white solid (6.75 mg, 5.46 µmol, 9.30% yield), 14% 2-methylbut-3-en-2-yl ether cleaved. **LC-MS (ESI)**: $t_R$ = 4.26 min, *m/z* = 1166.67 calcd. for $[C_{62}H_{95}N_5O_{14}S + H]^+$, found 1166.58; $t_R$ = 4.69 min, *m/z* = 1234.73 calcd. for $[C_{67}H_{103}N_5O_{14}S + H]^+$, found 1234.67. **HRMS (ESI-TOF)**: *m/z* = 583.8376 calcd. for $[C_{62}H_{95}N_5O_{14}S + 2H]^{2+}$, found 583.8370 [$\delta$ -1.0 ppm], *m/z* = 617.8690 calcd. for $[C_{67}H_{103}N_5O_{14}S + 2H]^{2+}$, found 617.8689 [$\delta$ -0.2 ppm]. **$^1$H NMR** (399 MHz, DMSO) $\delta$ 9.02 (s, 1H), 8.55 (t, *J* = 6.1 Hz, 1H), 7.82 (d, *J* = 9.3 Hz, 1H), 7.59 – 7.51 (m, 1H), 7.43 – 7.37 (m, 4H), 5.77 (dd, *J* = 17.7, 10.8 Hz, 1H), 5.30 (s, 1H), 5.11 (dd, *J* = 17.7, 1.5 Hz, 1H), 5.06 (dd, *J* = 10.8, 1.4 Hz, 1H), 4.73 (d, *J* = 3.8 Hz, 1H), 4.54 (d, *J* = 9.4 Hz, 1H), 4.41 – 4.35 (m, 3H), 4.23 (dd, *J* = 16.1, 5.3 Hz, 1H), 3.72 – 3.58 (m, 6H), 3.51 – 3.37 (m, 3H), 3.27 – 3.13 (m, 8H), 3.09 – 2.96 (m, 3H), 2.68 – 2.59 (m, 2H), 2.46 – 2.42 (m, 3H), 2.30 – 1.96 (m, 7H), 1.93 – 1.78 (m, 4H), 1.51 – 1.40 (m, 7H), 1.25 – 1.13 (m, 21H), 0.94 – 0.86 (m, 12H), 0.72 (d, *J* = 7.0 Hz, 3H) ppm. **$^{13}$C NMR** (100 MHz, DMSO) $\delta$ 172.5, 172.1, 171.9, 169.7, 151.6, 147.5, 144.1, 140.9, 139.6, 139.5, 137.1, 131.3, 129.5, 128.6, 127.4, 127.0, 113.6, 102.2, 80.8, 77.8, 77.6, 75.3, 74.5, 73.1, 72.2, 72.0, 70.5, 68.8, 62.5, 58.7, 57.9, 56.3, 56.3, 52.4, 48.2, 42.7, 41.6, 41.0, 37.9, 35.4, 35.2, 35.2, 34.9, 33.4, 29.0, 29.0, 28.8, 28.8, 28.7, 27.1, 26.4, 25.8, 25.7, 25.4, 25.4, 23.3, 16.0, 15.8, 9.2 ppm.

**CV3a** ($N^1$-((*S*)-2-((1*S*,4*R*,5*R*,6*R*,6a*S*,9*S*,10a*R*,*E*)-1,5-dihydroxy-9-(methoxymethyl)-6,10a-dimethyl-4-(((2*S*,3*R*,4*S*,5*S*,6*R*)-3,4,5-trihydroxy-6-(((2-methylbut-3-en-2-yl)oxy)methyl)tetrahydro-2*H*-pyran-2-yl)oxy)-1,2,4,5,6,6a,7,8,9,10a-decahydrodicyclopenta[*a*,*d*][8]annulen-3-yl)propyl)-$N^{19}$-((*S*)-1-((2*S*,4*R*)-4-hydroxy-2-((4-(4-methylthiazol-5-yl)benzyl)carbamoyl)pyrrolidin-1-yl)-3,3-dimethyl-1-oxobutan-2-yl)nonadecanediamide)

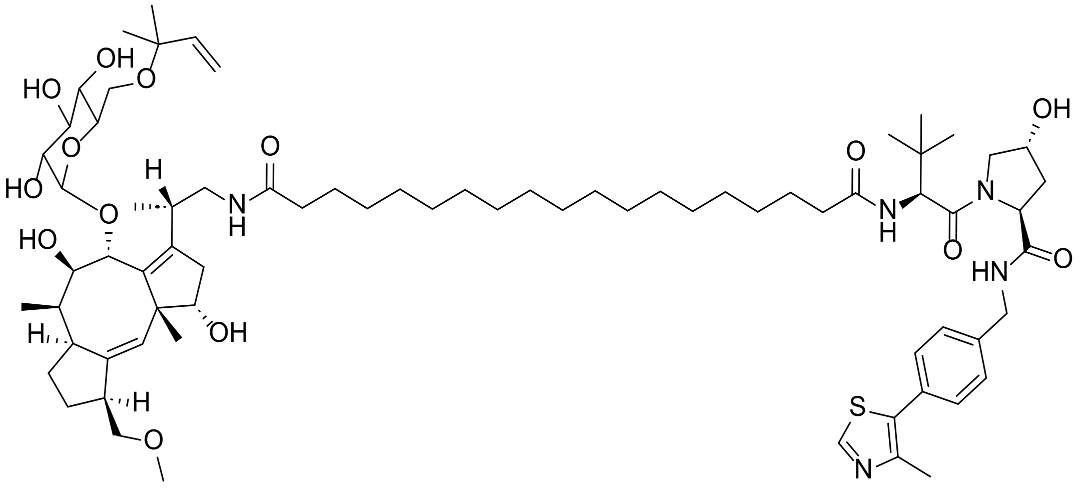

The intermediate **VHL3a** was synthesized via general procedure **B** using nonadecanedioic acid (352 mg, 1.07 mmol, 5 eq), HATU (366 mg, 0.964 mmol, 4.5 eq) DIPEA (1.12 mL, 6.42 mmol, 30 eq) and VHL ligand 1 HCl salt (100 mg, 0.214 mmol, 1 eq) in dry DMF (20 mL). The crude product was purified by reversed-phase preparative HPLC (85-90% acetonitrile (+0.1% FA) in water (+0.1% FA) gradient) to afford intermediate **VHL3a** as a white solid (87.4 mg, 0.118 mmol, 55.1% yield). **LC-MS (ESI)**: $t_R$ = 7.22 min, *m/z* = 741.46 calcd. for $[C_{41}H_{64}N_4O_6S + H]^+$, found 740.92. **HRMS (ESI-TOF)**: *m/z* = 741.4625 calcd. for $[C_{41}H_{64}N_4O_6S + H]^+$, found 741.4627 [$\delta$ 0.3 ppm].

**CV3a** was synthesized via general procedure **C** using **VHL3a** (47.9 mg, 0.0646 mmol, 1.1 eq), HATU (29.0 mg, 0.0764 mmol, 1.3 eq), DIPEA (102 µL, 0.587 mmol, 10 eq) and **FC-NH$_2$** (35 mg, 0.0588 mmol, 1 eq) in dry DMF (5 mL). The crude product was purified by reversed-phase preparative HPLC (70-75% acetonitrile (+0.1% FA) in water (+0.1% FA) gradient) to afford **CV3a** as a white solid (6.82 mg, 5.17 µmol, 8.80% yield), 20% 2-methylbut-3-en-2-yl ether cleaved. **LC-MS (ESI)**: $t_R$ = 4.99 min, *m/z* = 1250.76 calcd. for $[C_{68}H_{107}N_5O_{14}S + H]^+$, found 1250.67; $t_R$ = 5.44 min, *m/z* = 1318.82 calcd. for $[C_{73}H_{115}N_5O_{14}S + H]^+$, found 1318.83. **HRMS (ESI-TOF)**: *m/z* = 625.8846 calcd. for $[C_{68}H_{107}N_5O_{14}S + 2H]^{2+}$, found 625.8867 [$\delta$ 3.4 ppm]; *m/z* = 659.9159 calcd. for $[C_{73}H_{115}N_5O_{14}S + 2H]^{2+}$, found 659.9147 [$\delta$ -1.8 ppm]. **$^1$H NMR** (399 MHz, DMSO) $\delta$ 9.05 (s, 1H), 8.55 (t, *J* = 6.1 Hz, 1H), 7.83 (d, *J* = 9.4 Hz, 1H), 7.59 – 7.52 (m, 1H), 7.43 – 7.38 (m, 4H), 5.77 (dd, *J* = 17.7, 10.8 Hz, 1H), 5.30 (s, 1H), 5.13 (dd, *J* = 17.7, 1.5 Hz, 1H), 5.05 (dd, *J* = 10.7, 1.4 Hz, 1H), 4.75 – 4.73 (m, 1H), 4.53 (d, *J* = 2.9 Hz, 1H), 4.44 – 4.34 (m, 3H), 4.21 (dd, *J* = 15.9, 5.3 Hz, 1H), 3.72 – 3.59 (m, 6H), 3.49 – 3.38 (m, 3H), 3.25 – 3.15 (m, 8H), 3.07 – 2.95 (m, 3H), 2.65 – 2.60 (m, 2H), 2.45 (s, 3H), 2.24 – 2.03 (m, 7H), 1.90 – 1.80 (m, 4H), 1.48 – 1.40 (m, 7H), 1.23 – 1.17 (m, 33H), 0.94 – 0.90 (m, 12H), 0.72 (d, *J* = 7.1 Hz, 3H) ppm. **$^{13}$C NMR** (100 MHz, DMSO) $\delta$ 172.5, 172.1, 171.9, 169.7, 151.7, 147.3, 144.1, 140.9, 139.6, 139.5, 137.1, 131.4, 129.4, 128.6, 127.4, 127.0, 113.6, 102.2, 80.8, 77.9, 77.6, 75.3, 74.5, 73.1, 72.2, 72.0, 70.5, 68.8, 62.5, 58.7, 57.9, 56.3, 52.4, 48.2, 42.7, 41.6, 41.0, 37.9, 35.3, 35.2, 34.8, 33.4, 29.1, 29.0, 29.0, 29.0, 28.8, 28.7, 28.6, 27.1, 26.4, 25.8, 25.7, 25.4, 25.3, 23.3, 16.0, 15.7, 9.2 ppm.

**(*S*-OH)-CV2a** (*N*[1]-((*S*)-2-((1*S*,4*R*,5*R*,6*R*,6a*S*,9*S*,10a*R*,*E*)-1,5-dihydroxy-9-(methoxymethyl)-6,10a-dimethyl-4-(((2*S*,3*R*,4*S*,5*S*,6*R*)-3,4,5-trihydroxy-6-(((2-methylbut-3-en-2-yl)oxy)methyl)tetrahydro-2*H*-pyran-2-yl)oxy)-1,2,4,5,6,6a,7,8,9,10a-decahydrodicyclopenta[*a*,*d*][8]annulen-3-yl)propyl)-*N*[13]-((*S*)-1-((2*S*,4*S*)-4-hydroxy-2-((4-(4-methylthiazol-5-yl)benzyl)carbamoyl)pyrrolidin-1-yl)-3,3-dimethyl-1-oxobutan-2-yl)tridecanediamide)

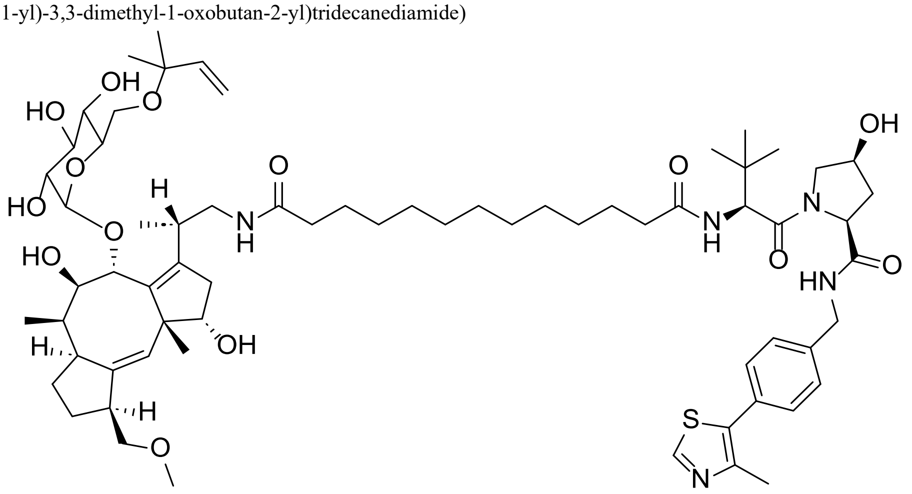

The intermediate **VHL(*S*-OH)3a** was synthesized via general procedure **B** using tridecanedioic acid (262 mg, 1.07 mmol, 5 eq), HATU (366 mg, 0.964 mmol, 4.5 eq) DIPEA (1.12 mL, 6.42 mmol, 30 eq) and VHL(*S*-OH) ligand 1 HCl salt (100 mg, 0.214 mmol, 1 eq) in dry DMF (15 mL). The crude product was purified by reversed-phase preparative HPLC (45-50% acetonitrile (+0.1% FA) in water (+0.1% FA) gradient) to afford intermediate **VHL(*S*-OH)3a** as a white solid (49.2 mg, 0.0749 mmol, 35.0% yield). **LC-MS (ESI)**: $t_R$ = 4.46 min, *m/z* = 657.37 calcd. for $[C_{35}H_{52}N_4O_6S + H]^+$, found 657.58. **HRMS (ESI-TOF)**: *m/z* = 657.3686 calcd. for $[C_{35}H_{52}N_4O_6S + H]^+$, found 657.3685 [$\delta$ -0.2 ppm].

**(*S*-OH)-CV2a** was synthesized via general procedure **C** using **VHL(*S*-OH)3a** (42.5 mg, 0.0646 mmol, 1.1 eq), HATU (29.0 mg, 0.0764 mmol, 1.3 eq), DIPEA (102 µL, 0.587 mmol, 10 eq) and **FC-NH₂** (35 mg, 0.0588 mmol, 1 eq) in dry DMF (5 mL). The crude product was purified by reversed-phase preparative HPLC (50-55% acetonitrile (+0.1% FA) in water (+0.1% FA) gradient) to afford **(*S*-OH)-CV2a** as a white solid (5.83 mg, 4.72 µmol, 8.03% yield), no 2-methylbut-3-en-2-yl ether cleavage. **LC-MS (ESI)**: $t_R$ = 4.72 min, *m/z* = 1234.73 calcd. for $[C_{67}H_{103}N_5O_{14}S + H]^+$, found 1234.92. **HRMS (ESI-TOF)**: *m/z* = 617.8690 calcd. for $[C_{67}H_{103}N_5O_{14}S + 2H]^{2+}$, found 617.8692 [$\delta$ 0.3 ppm]. **$^1$H NMR** (399 MHz, DMSO) $\delta$ 9.02 (s, 1H), 8.62 (t, *J* = 6.1 Hz, 1H), 7.83 (d, *J* = 8.8 Hz, 1H), 7.59 – 7.53 (m, 1H), 7.42 – 7.37 (m, 4H), 5.77 (dd, *J* = 17.7, 10.8 Hz, 1H), 5.31 – 5.28 (m, 1H), 5.11 (dd, *J* = 17.7, 1.5 Hz, 1H), 5.06 (dd, *J* = 10.8, 1.5 Hz, 1H), 4.73 (d, *J* = 3.8 Hz, 1H), 4.46 – 4.44 (m, 1H), 4.37 – 4.36 (m, 1H), 4.29 – 4.26 (m, 1H), 4.24 – 4.18 (m, 2H), 3.93 (dd, *J* = 10.2, 5.7 Hz, 1H), 3.78 – 3.48 (m, 5H), 3.48 – 3.41 (m, 3H), 3.26 – 3.15 (m, 8H), 3.06 – 2.95 (m, 3H), 2.67 – 2.60 (m, 2H), 2.45 (s, 3H), 2.35 – 2.04 (m, 7H), 1.88 – 1.71 (m, 4H), 1.51 – 1.40 (m, 7H), 1.24 – 1.13 (m, 21H), 0.96 – 0.88 (m, 12H), 0.72 (d, *J* = 7.0 Hz, 3H) ppm. **$^{13}$C NMR** (100 MHz, DMSO) $\delta$ 172.5, 172.5, 172.4, 170.0, 151.6, 147.6, 144.1, 140.9, 139.5, 139.3, 137.1, 131.2, 129.6, 128.7, 127.4, 127.0, 113.6, 102.2, 80.8, 77.9, 77.6, 75.3, 74.6, 73.1, 72.2, 72.1, 70.5, 69.1, 62.5, 58.5, 57.9, 56.6, 55.6, 52.4, 48.2, 42.7, 41.8, 41.0, 36.9, 35.4, 35.2, 34.8, 34.7, 33.4, 29.0, 28.9, 28.8, 28.8, 28.7, 27.1, 26.4, 25.8, 25.7, 25.4, 25.4, 23.3, 16.0, 15.9, 9.2 ppm.

#### NMR spectra

$^{1}$H NMR spectrum of **FC-NAc**

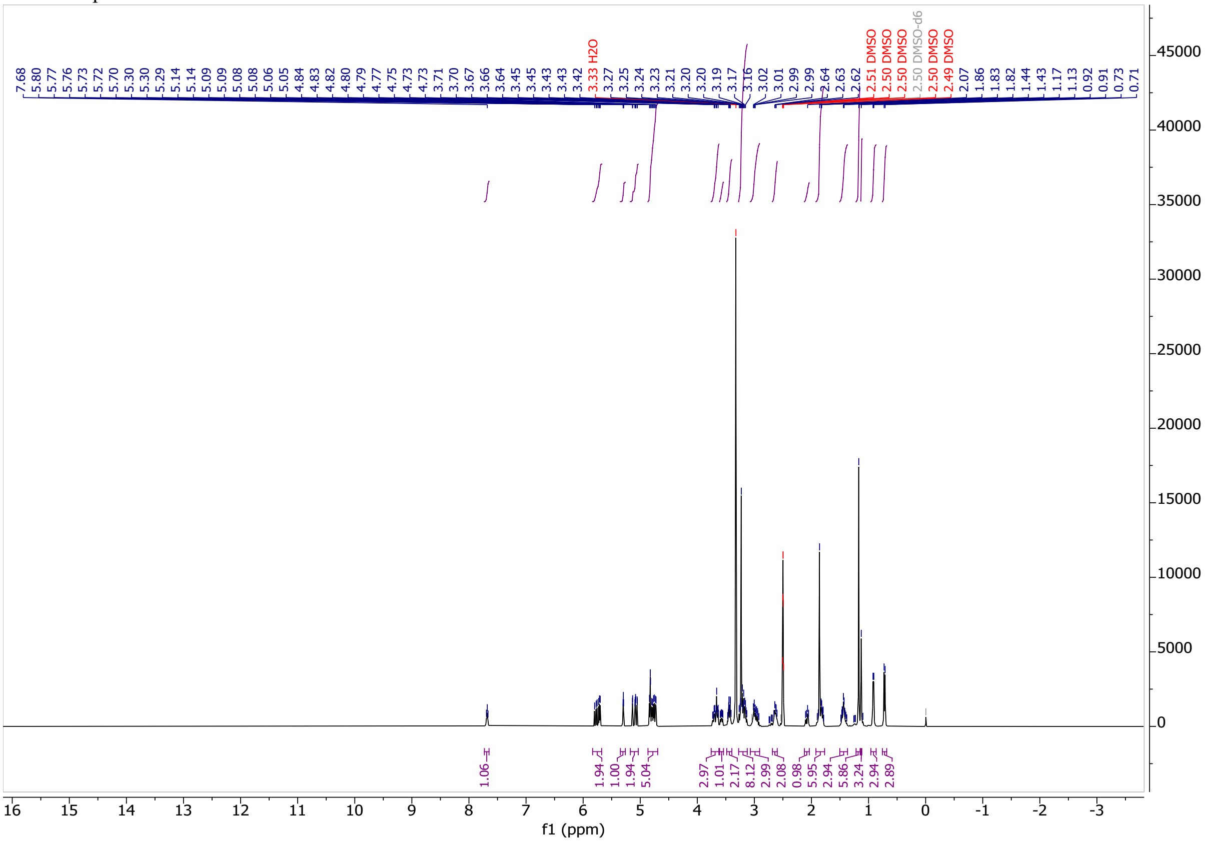

$^{13}$C NMR spectrum of **FC-NAc**

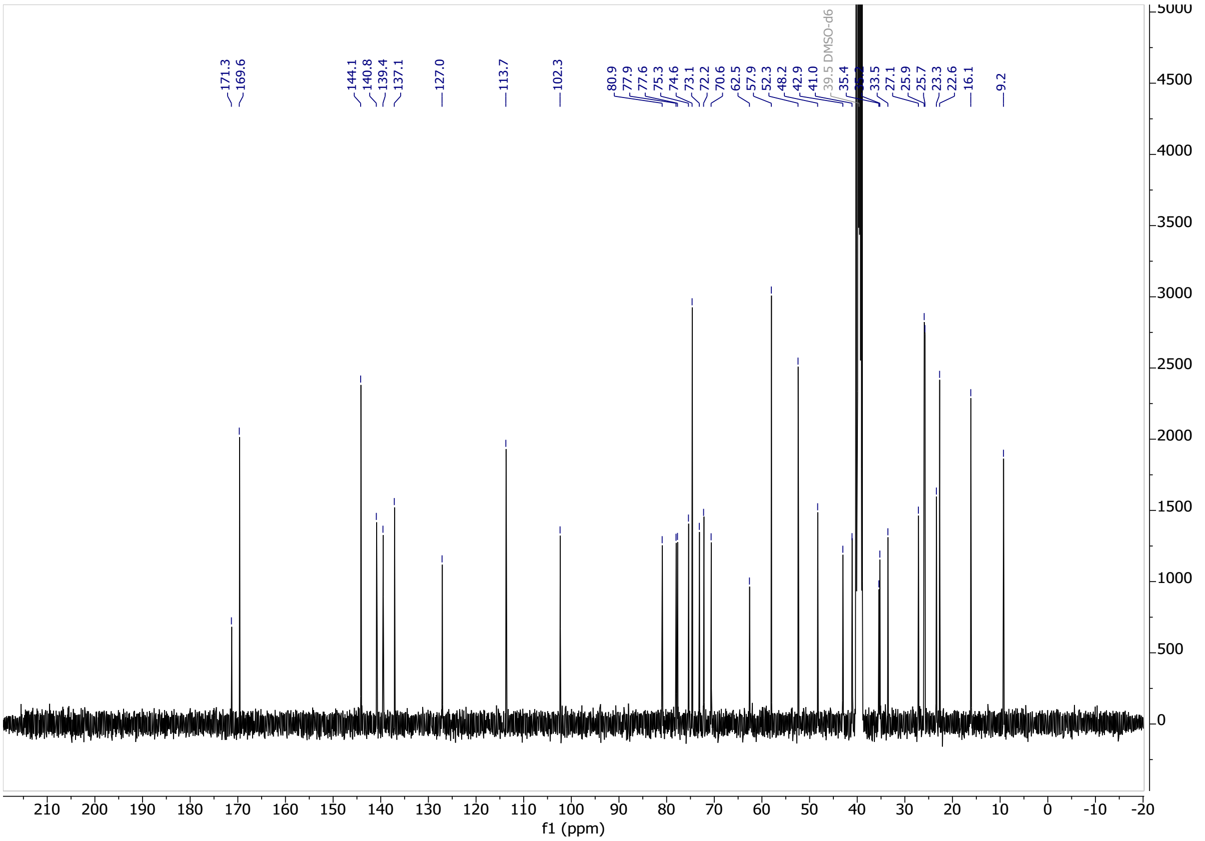

[1]H NMR spectrum of **VH032**

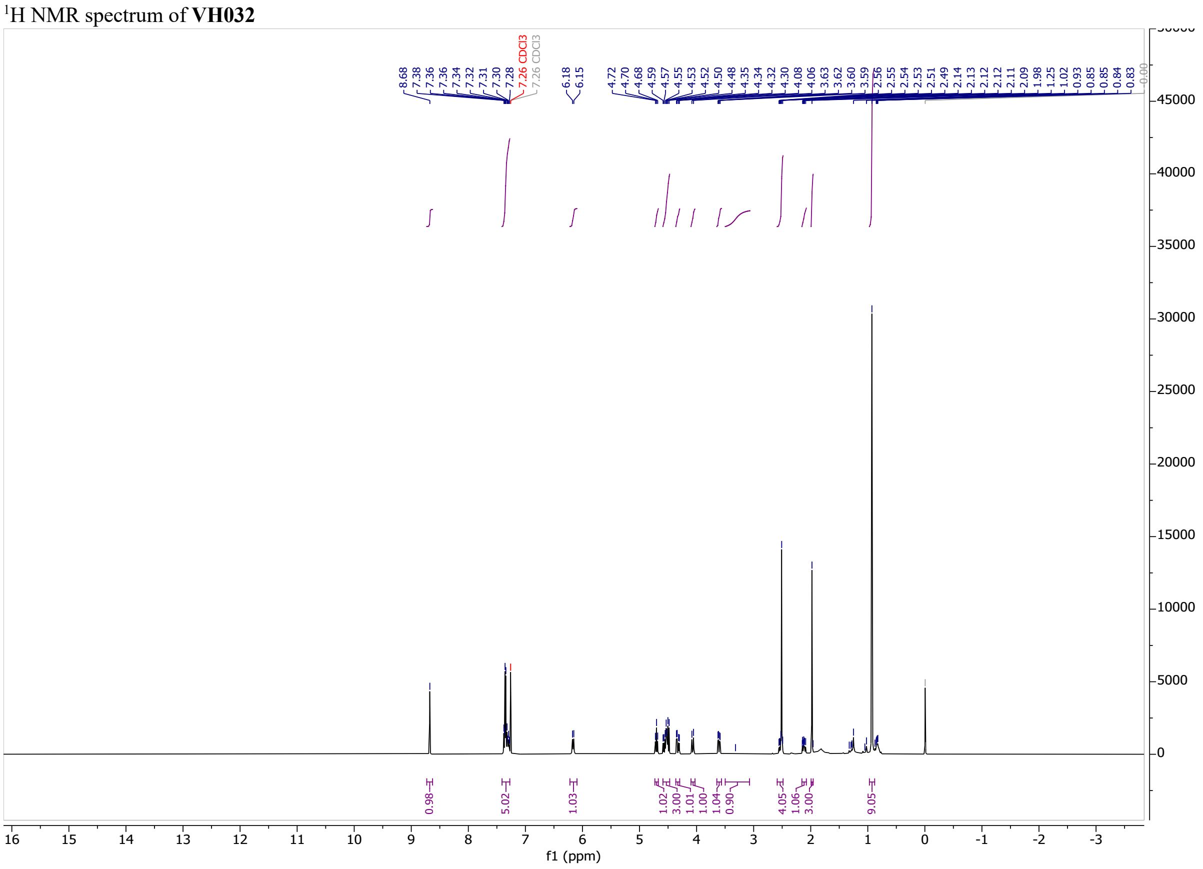

[13]C NMR spectrum of **VH032**

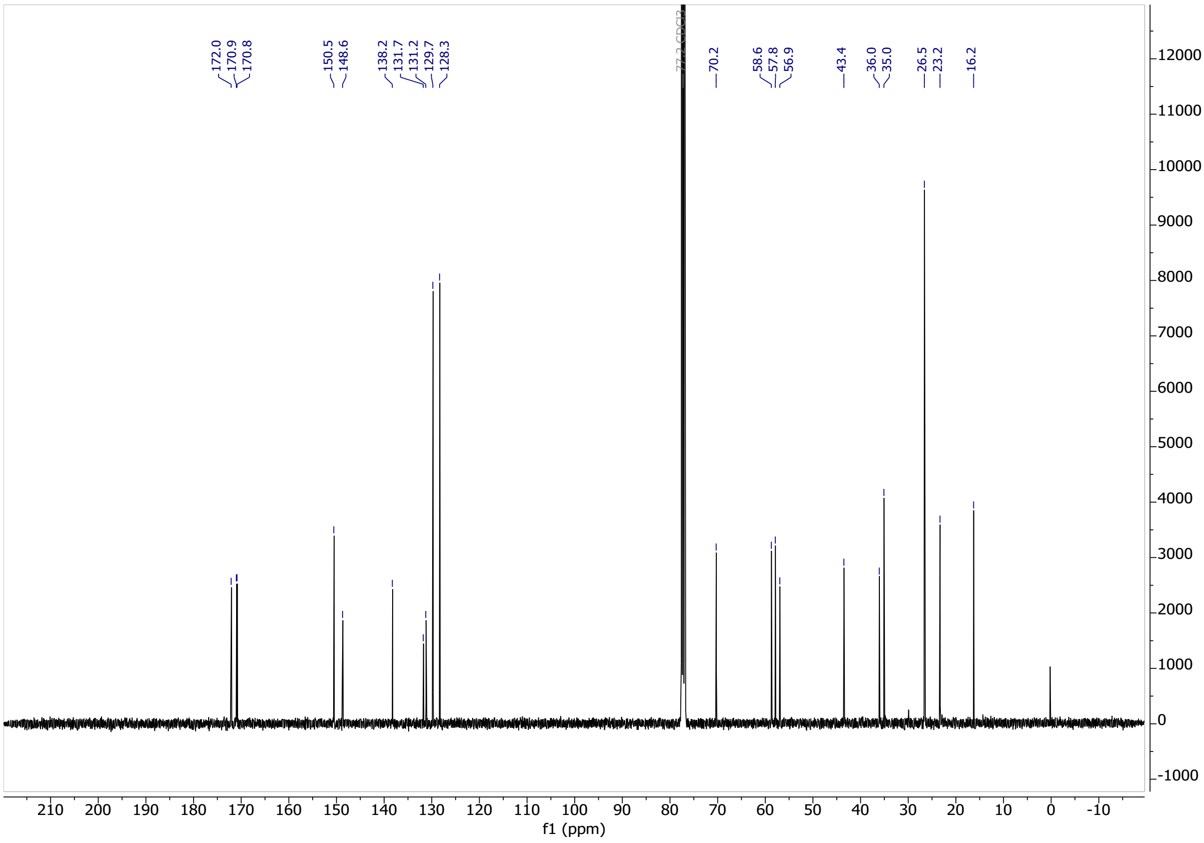

[1]H NMR spectrum of **2-methylbut-3-en-2-yl ether cleaved FC-NAc**

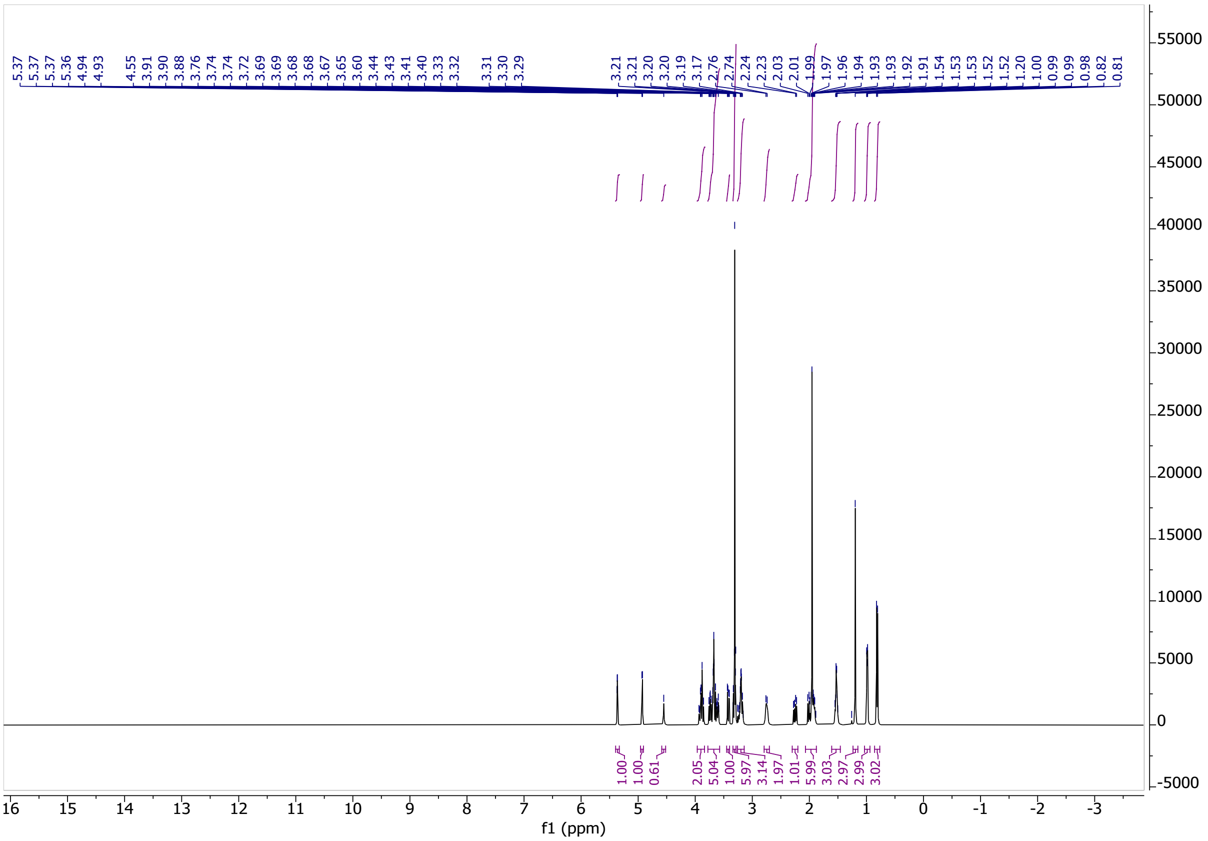

[13]C NMR spectrum of **2-methylbut-3-en-2-yl ether cleaved FC-NAc**

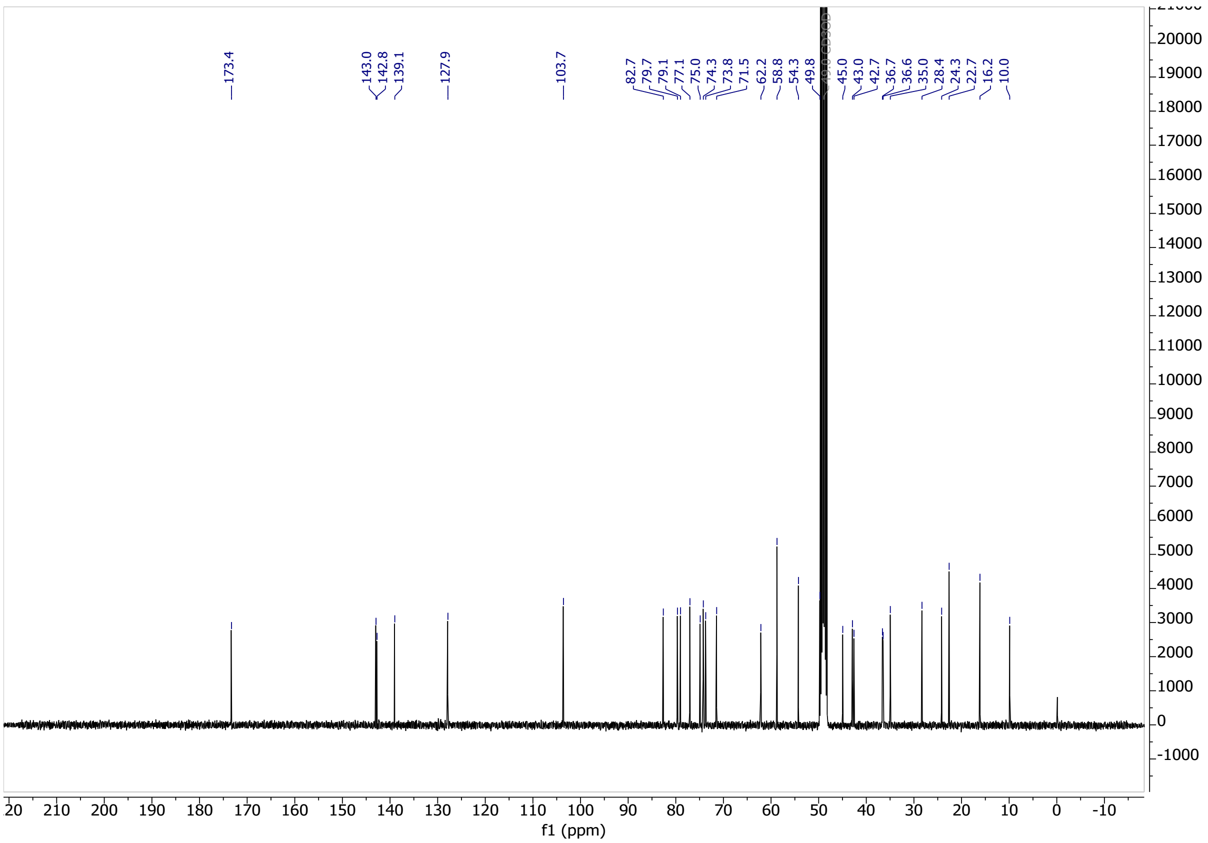

[1]H NMR spectrum of **CV1p**

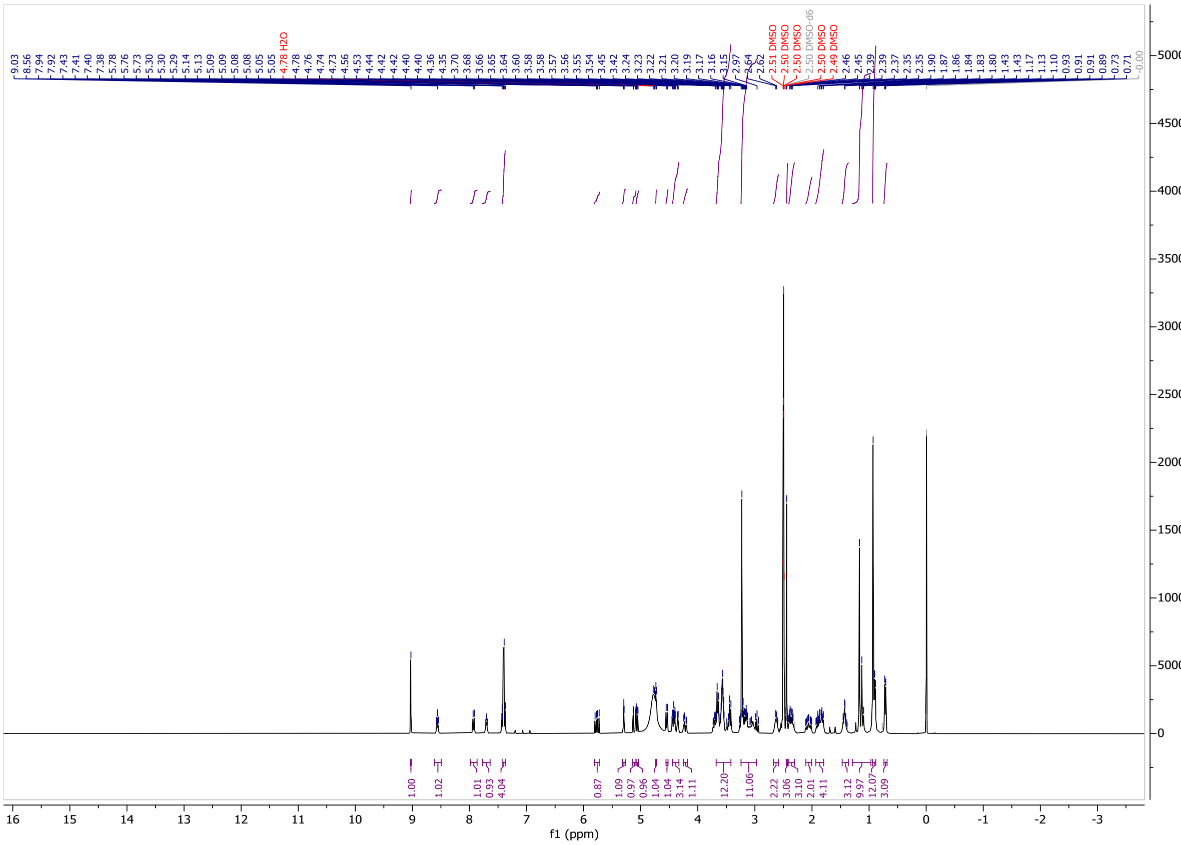

[13]C NMR spectrum of **CV1p**

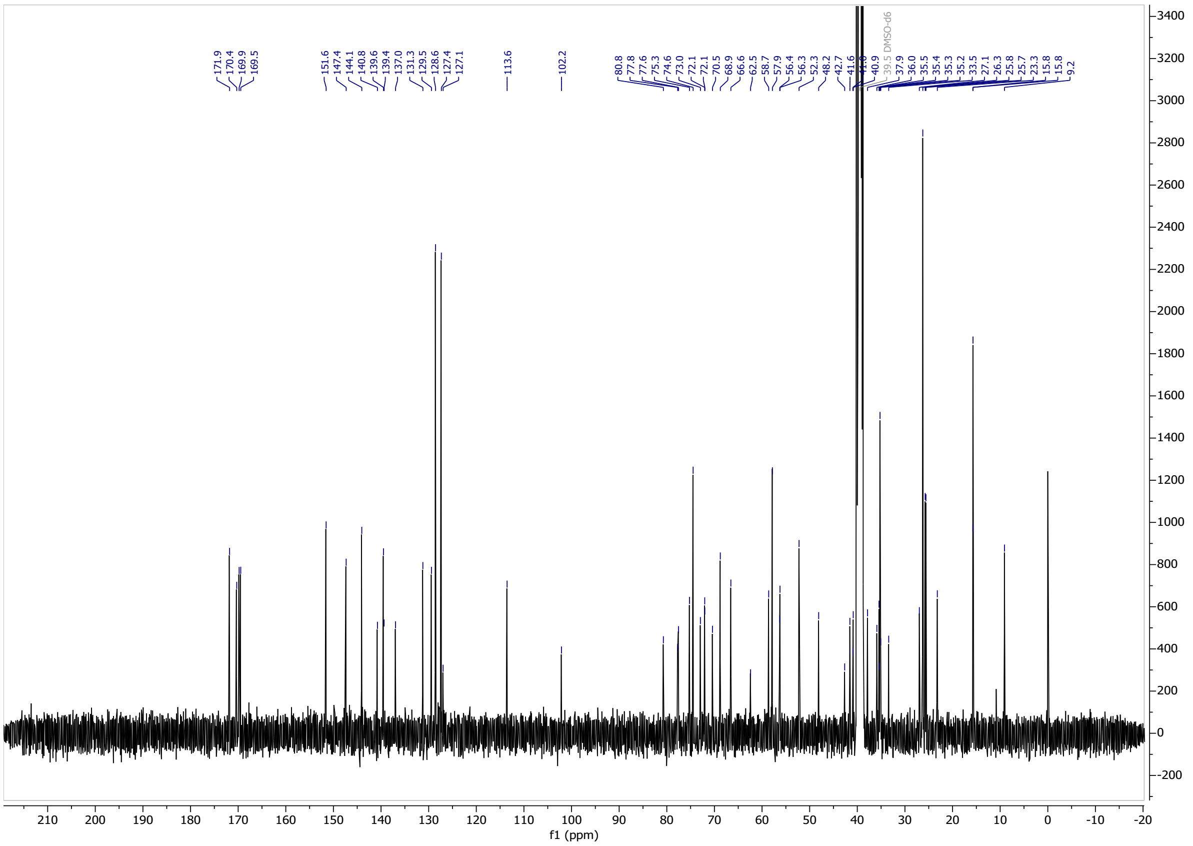

$^{1}H$ NMR spectrum of **CV2p**

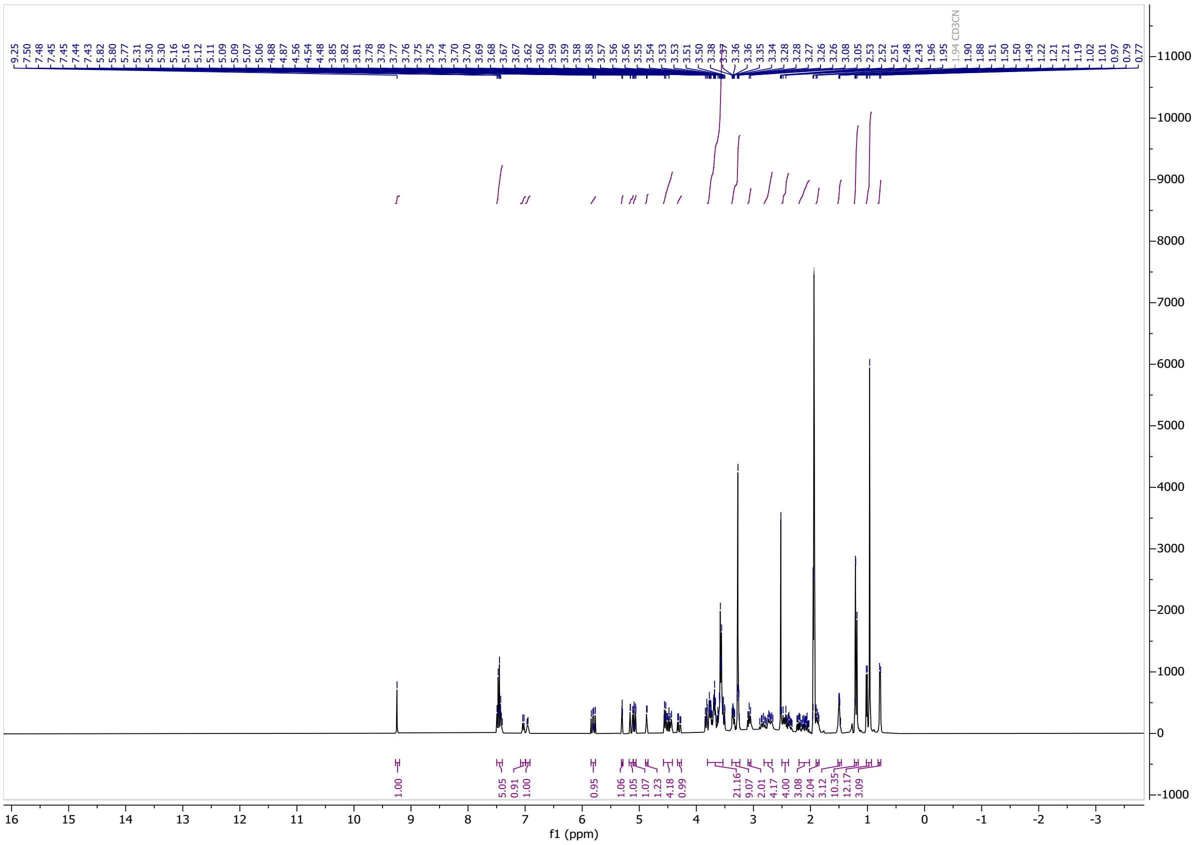

$^{13}C$ NMR spectrum of **CV2p**

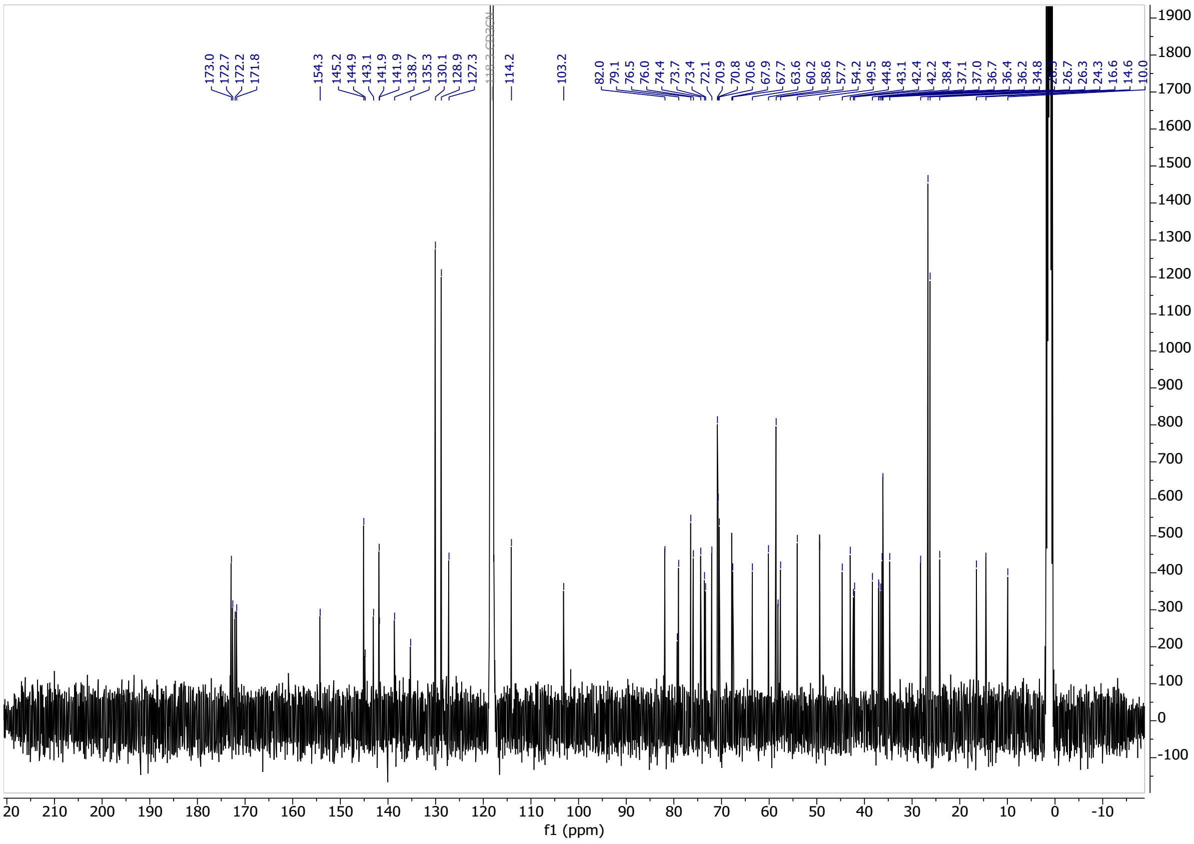

$^{1}$H NMR spectrum of **CV3p**

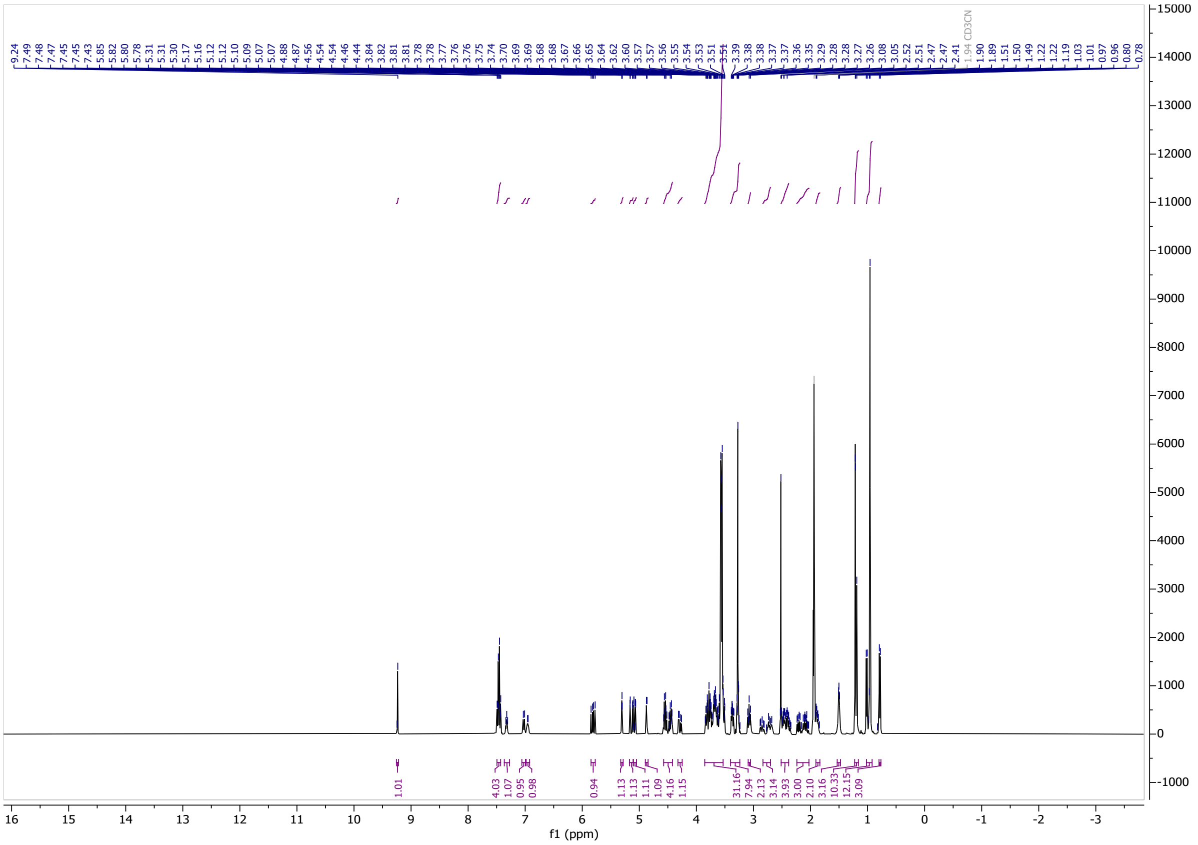

$^{13}$C NMR spectrum of **CV3p**

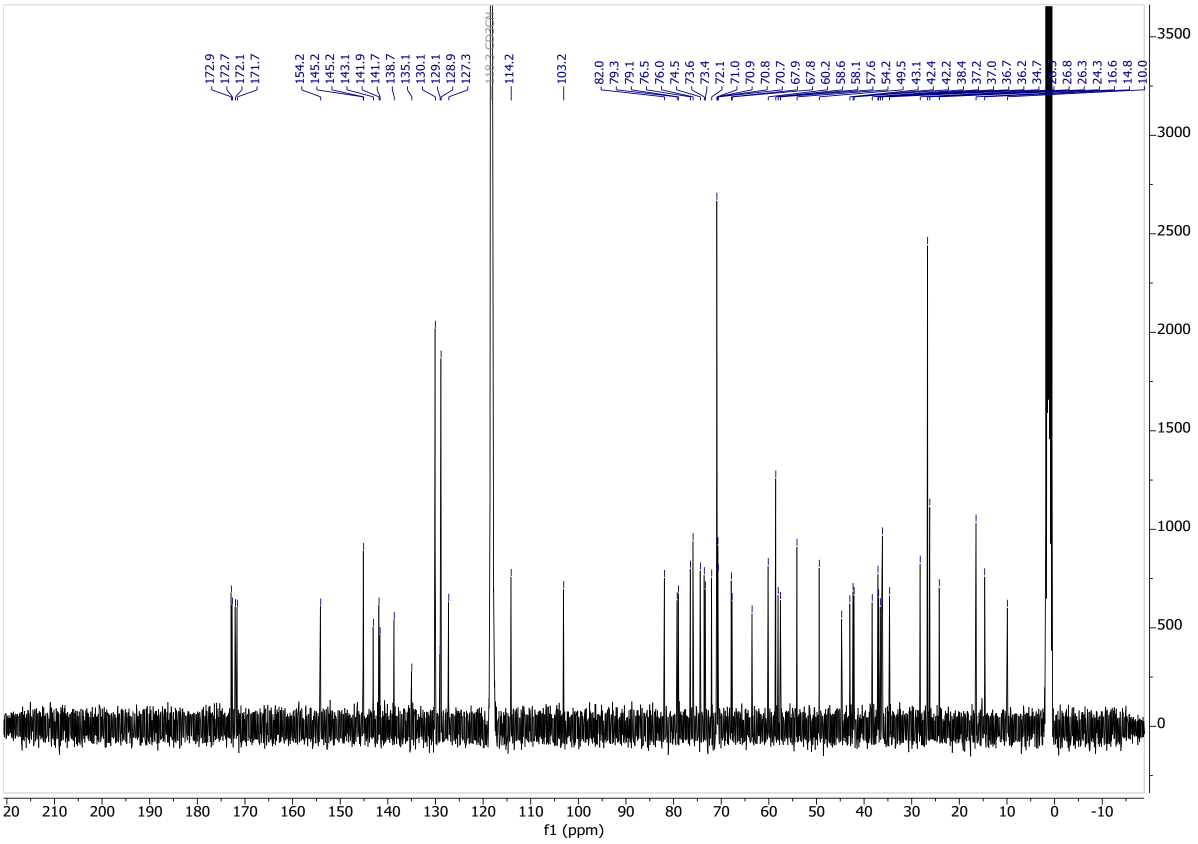

[1]H NMR spectrum of **CV4p**

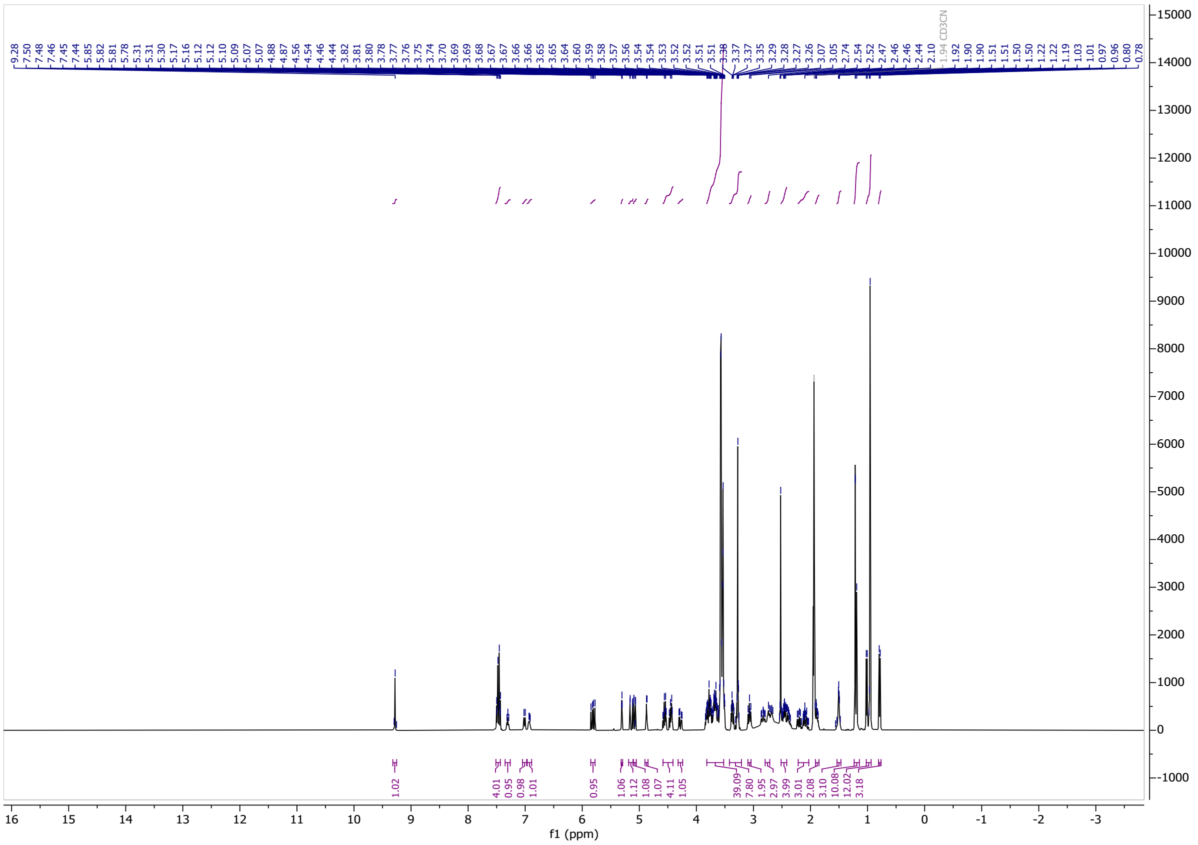

[13]C NMR spectrum of **CV4p**

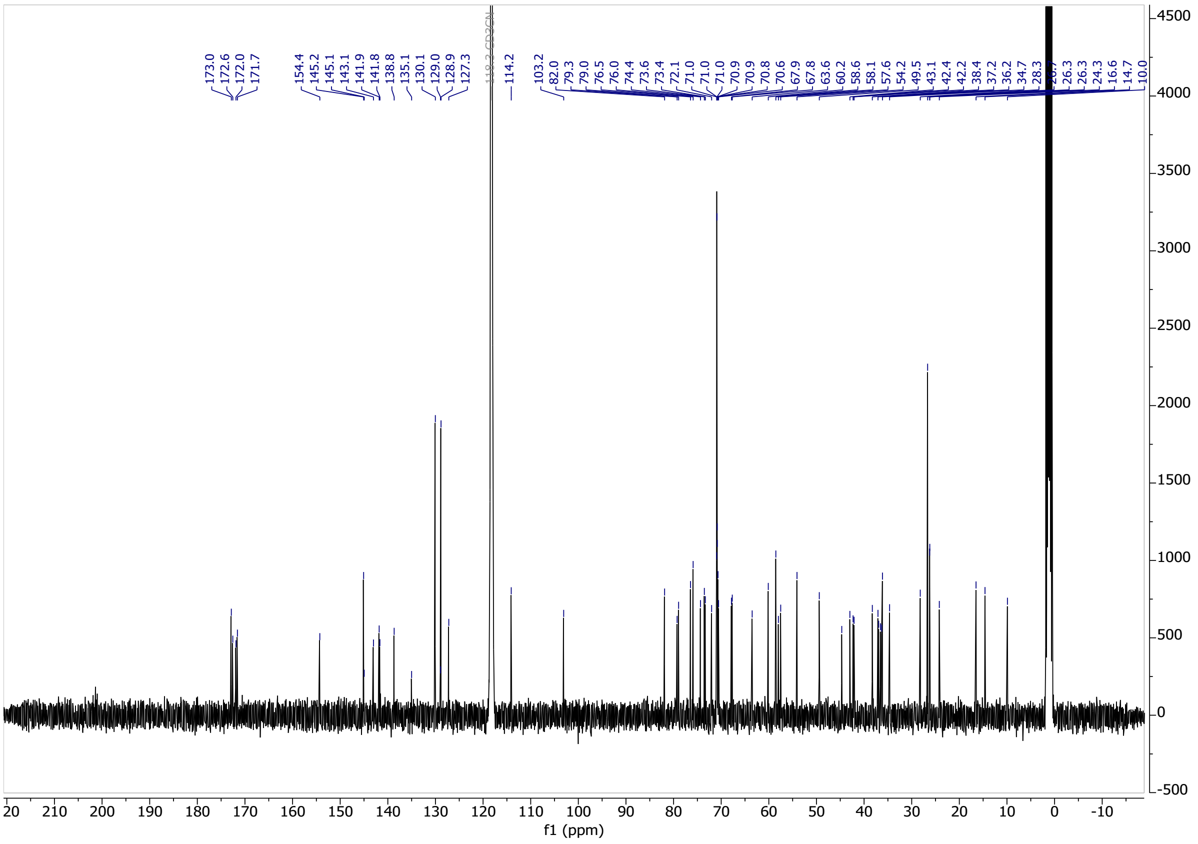

[1]H NMR spectrum of **CV0a**

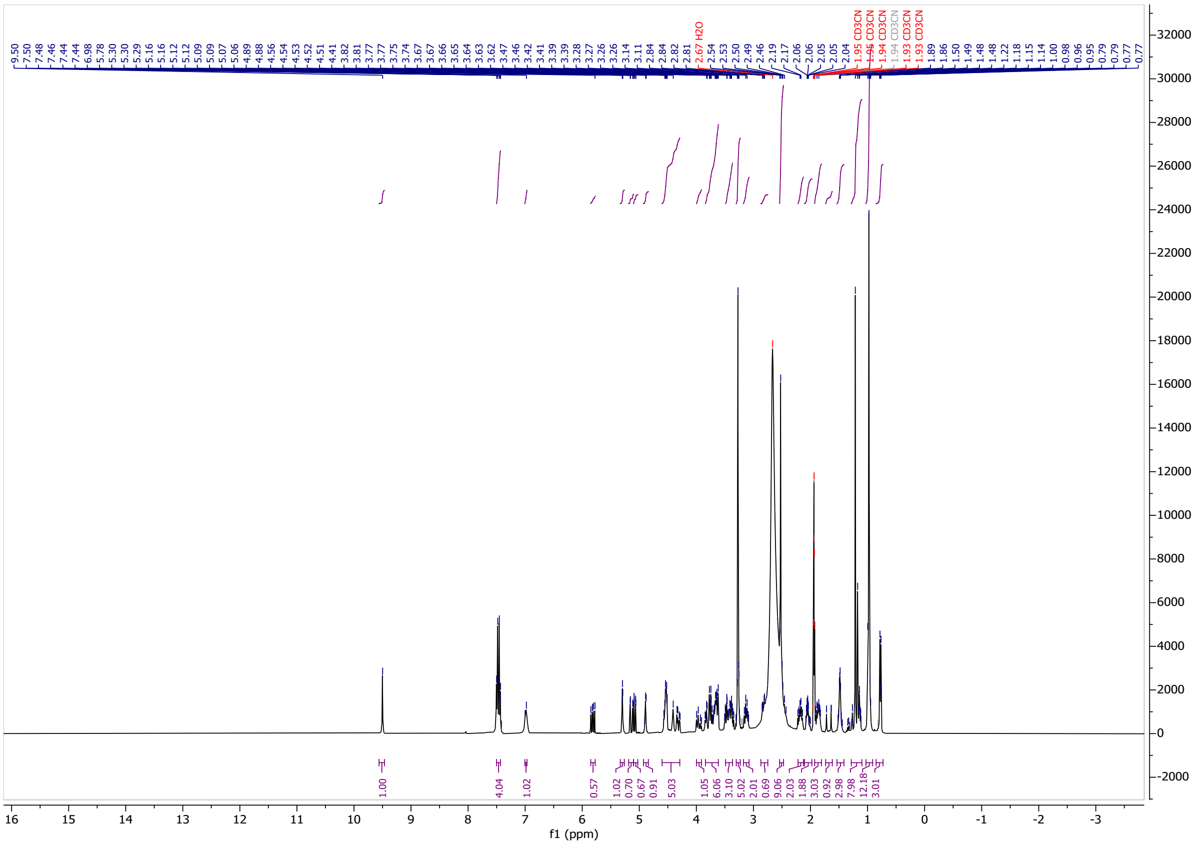

[13]C NMR spectrum of **CV0a**

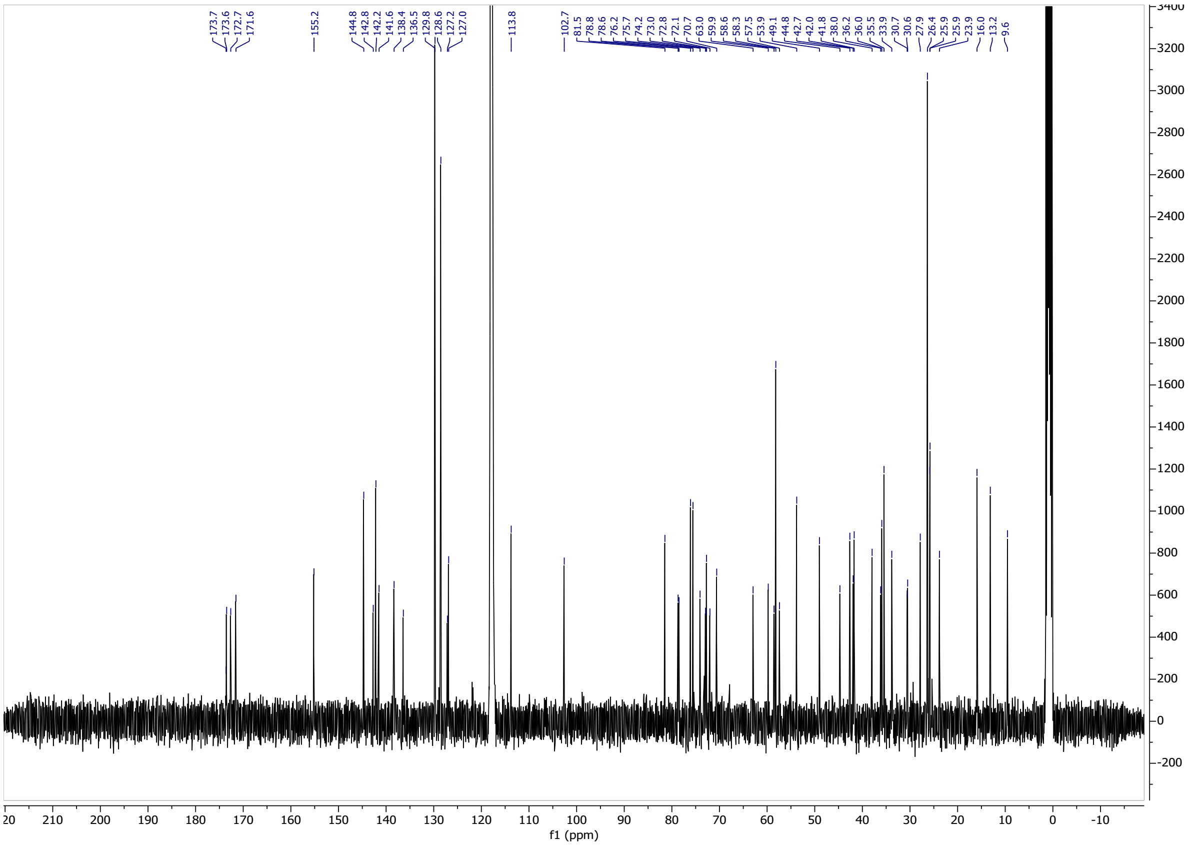

$^{1}$H NMR spectrum of **CV1a**

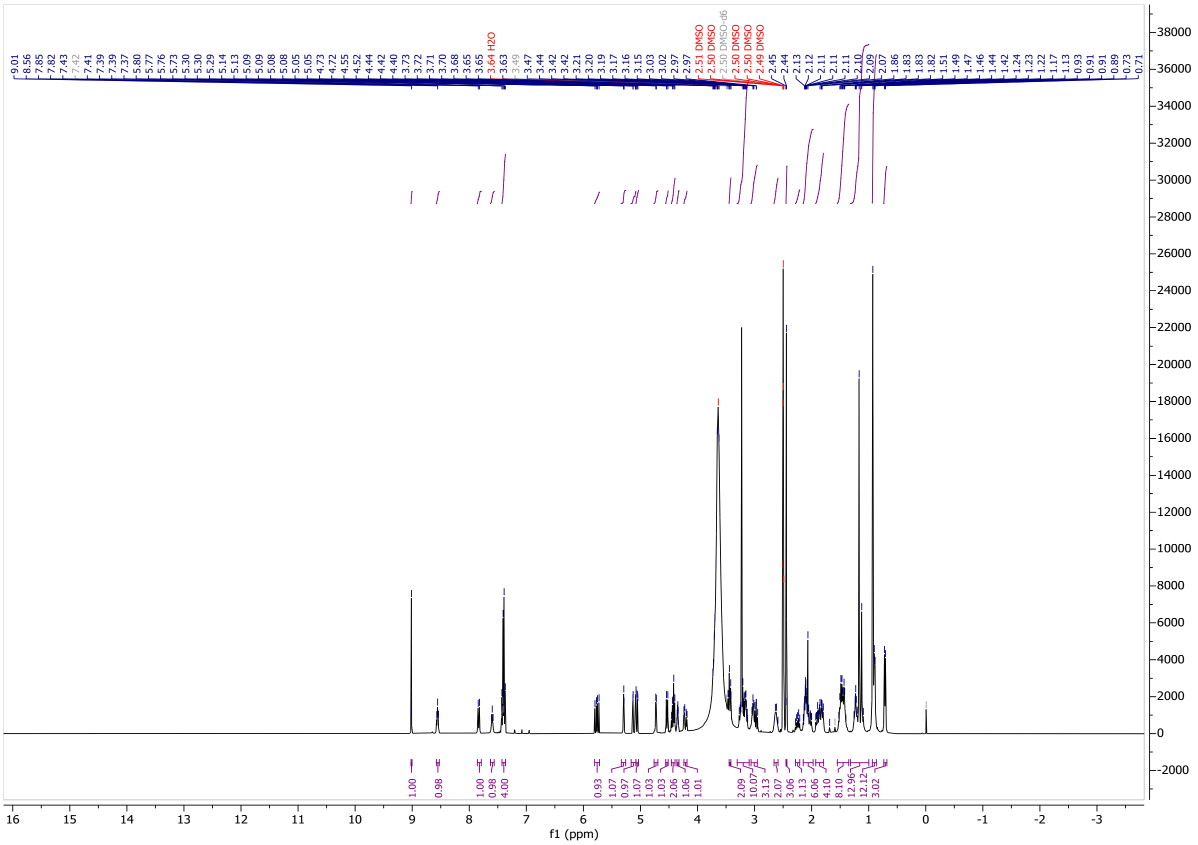

$^{13}$C NMR spectrum of **CV1a**

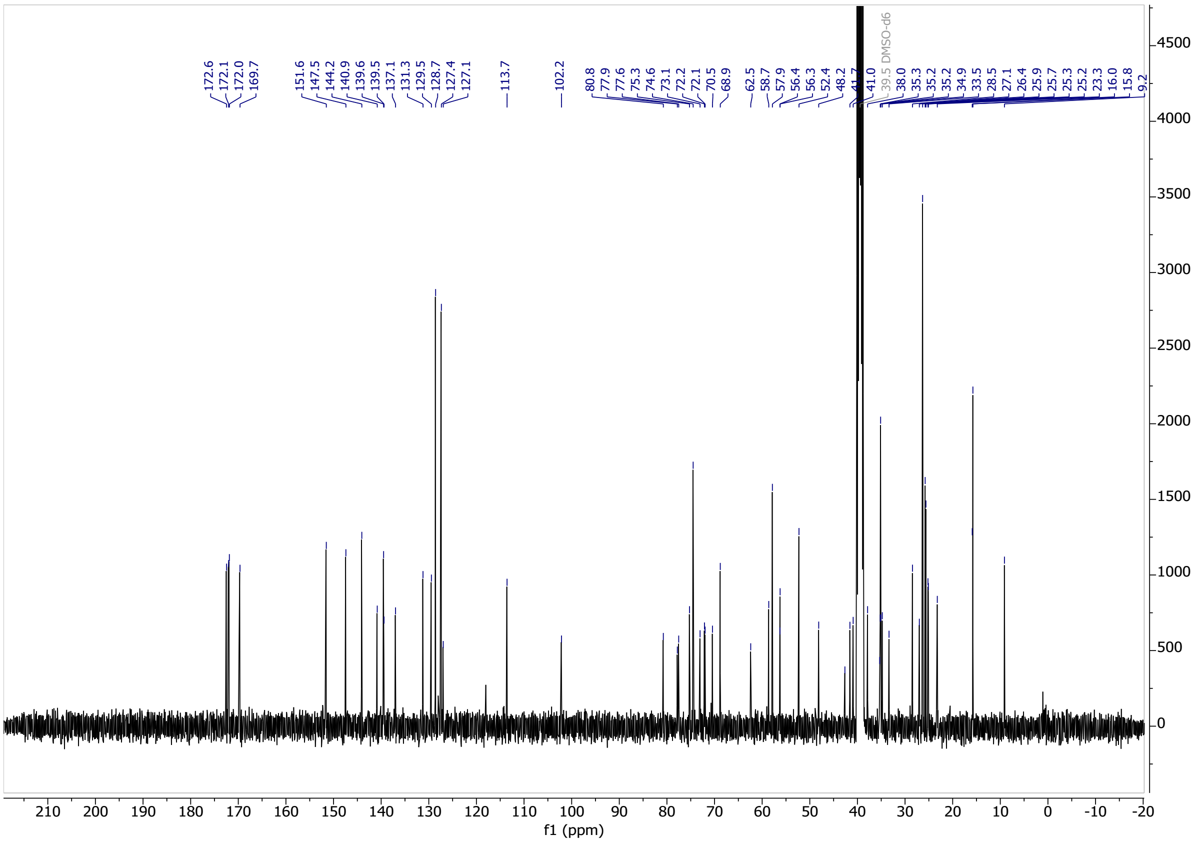

[1]H NMR spectrum of **CV2a**

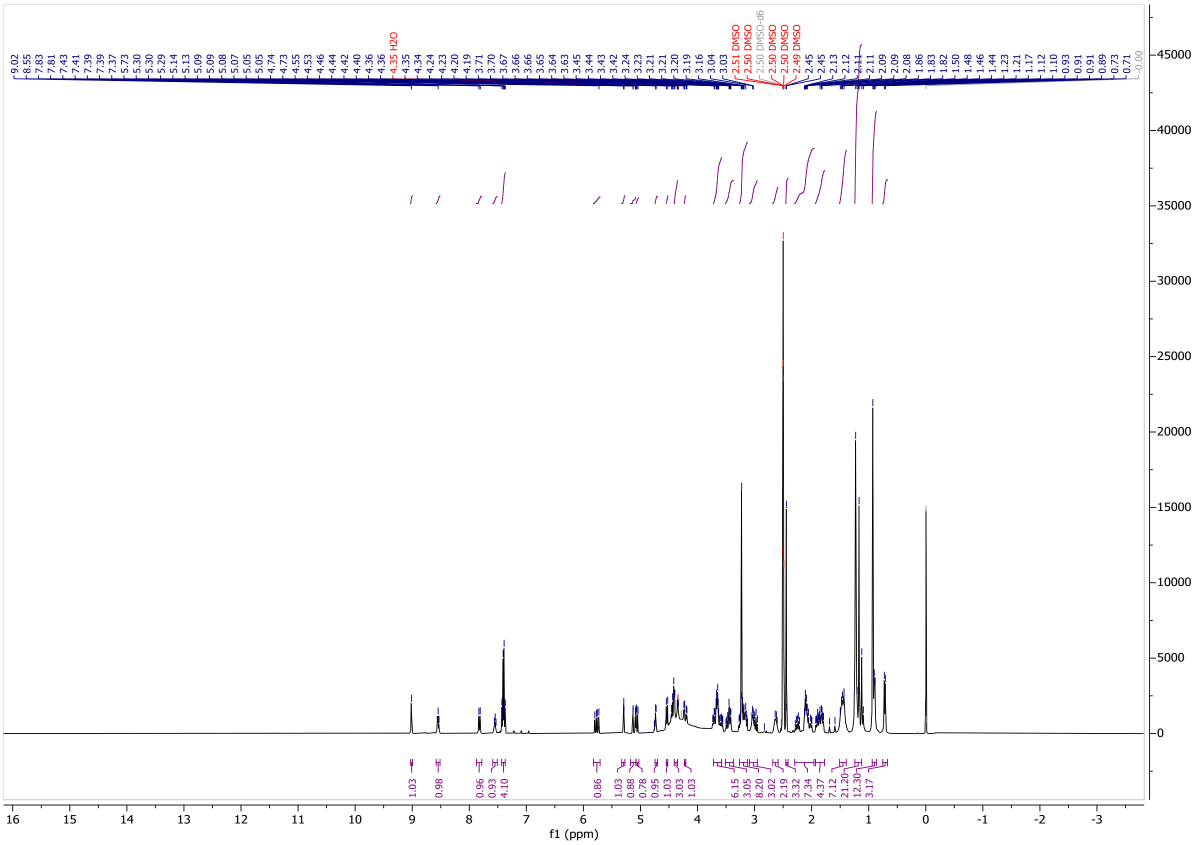

[13]C NMR spectrum of **CV2a**

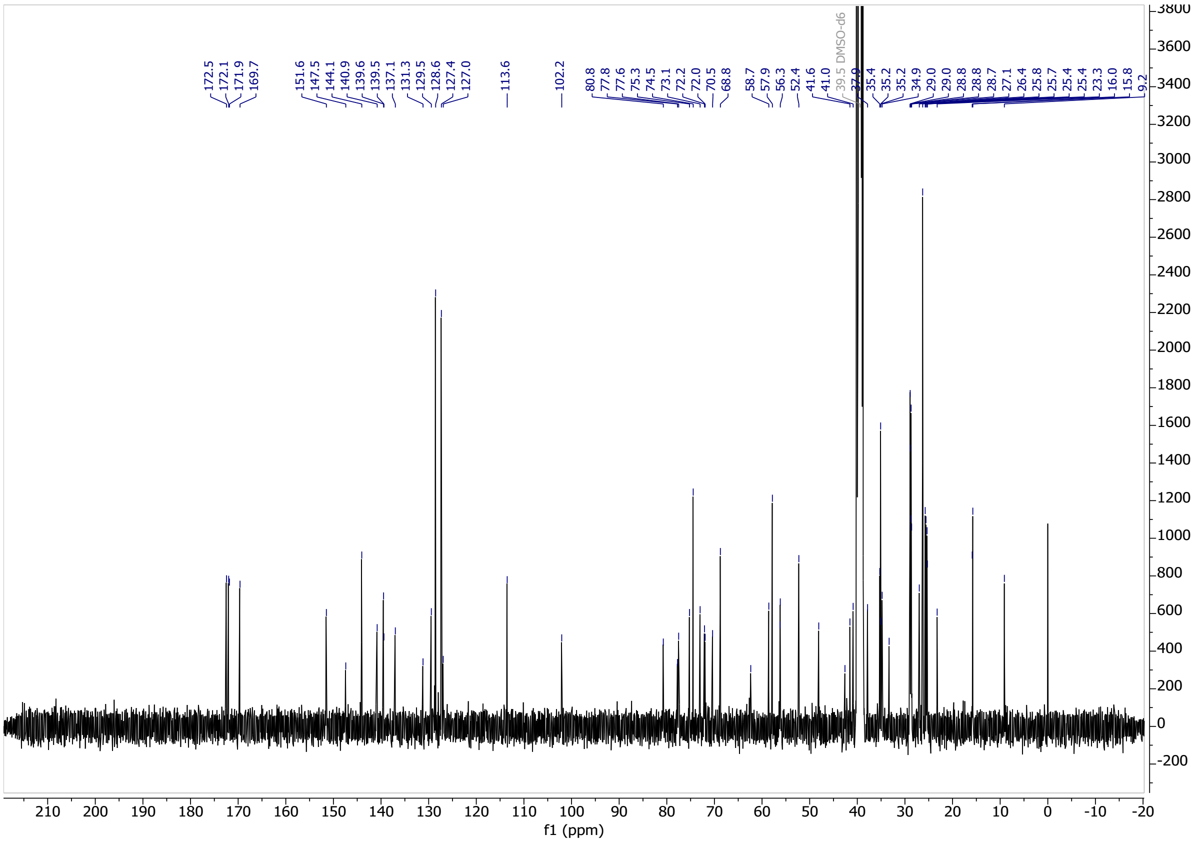

[1]H NMR spectrum of **CV3a**

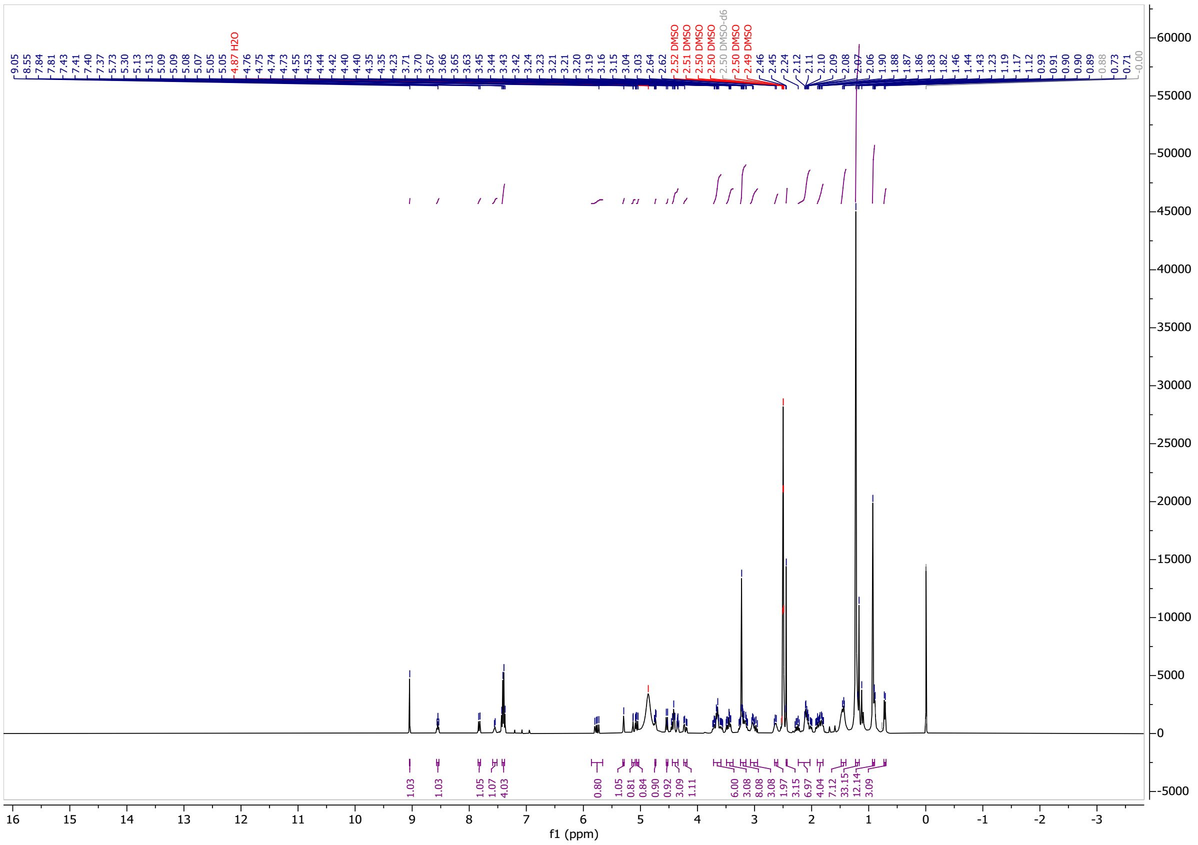

[13]C NMR spectrum of **CV3a**

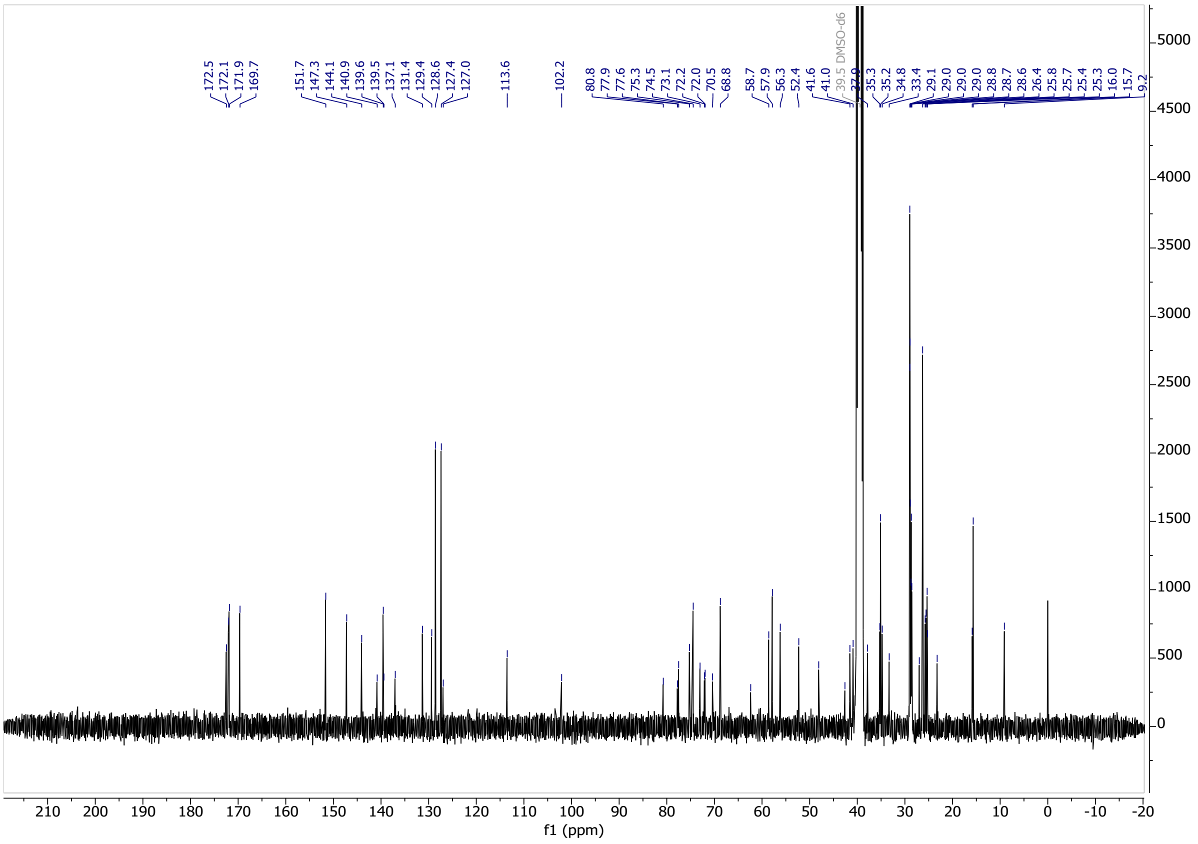

[1]H NMR spectrum of **(*S*-OH)-CV2a**

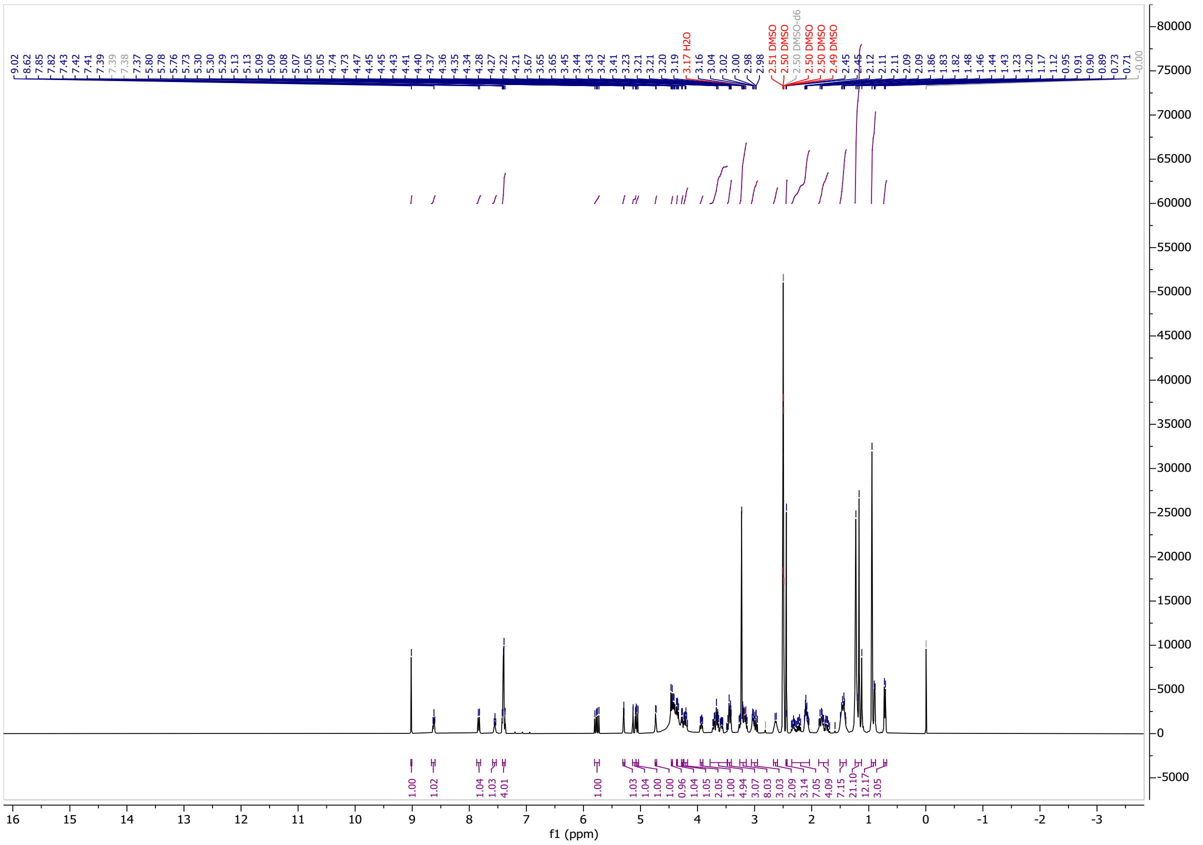

[13]C NMR spectrum of **(*S*-OH)-CV2a**

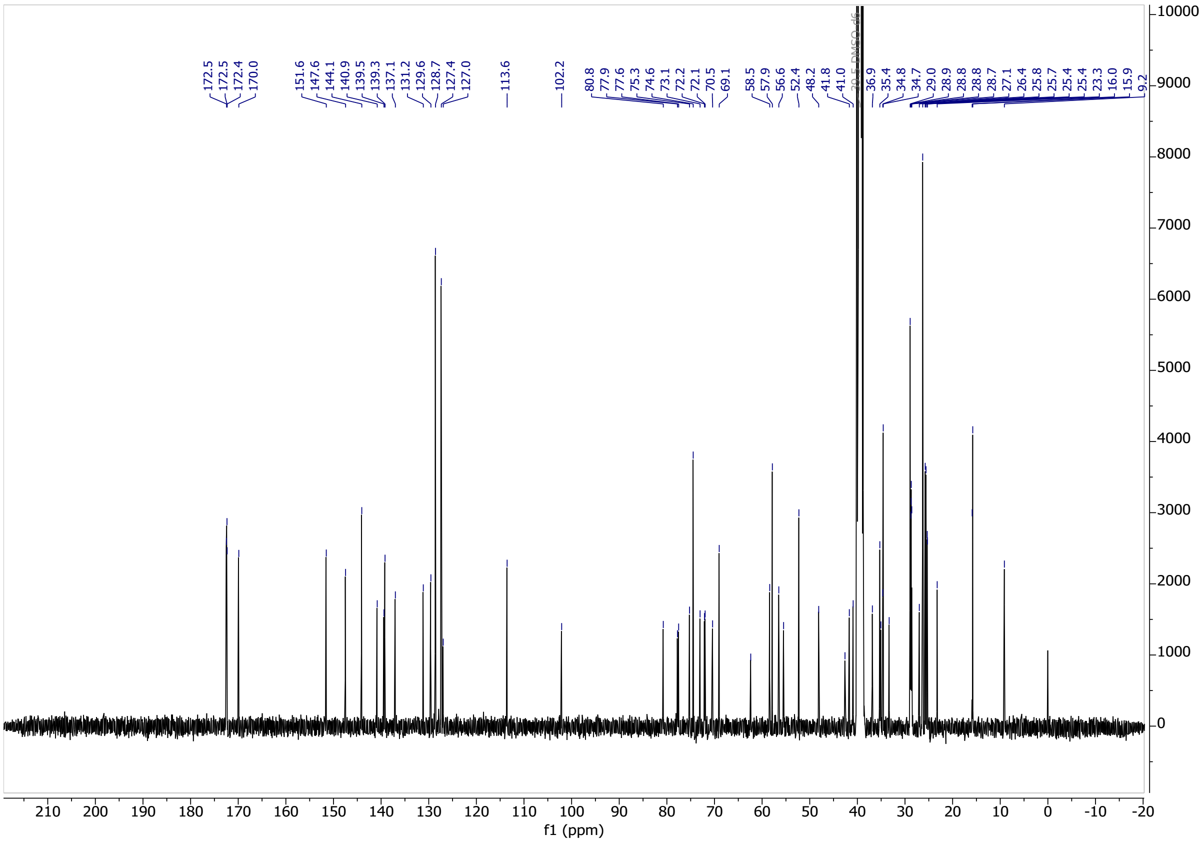

**HPLC chromatograms**

HPLC chromatogram of **FC-NH$_2$**

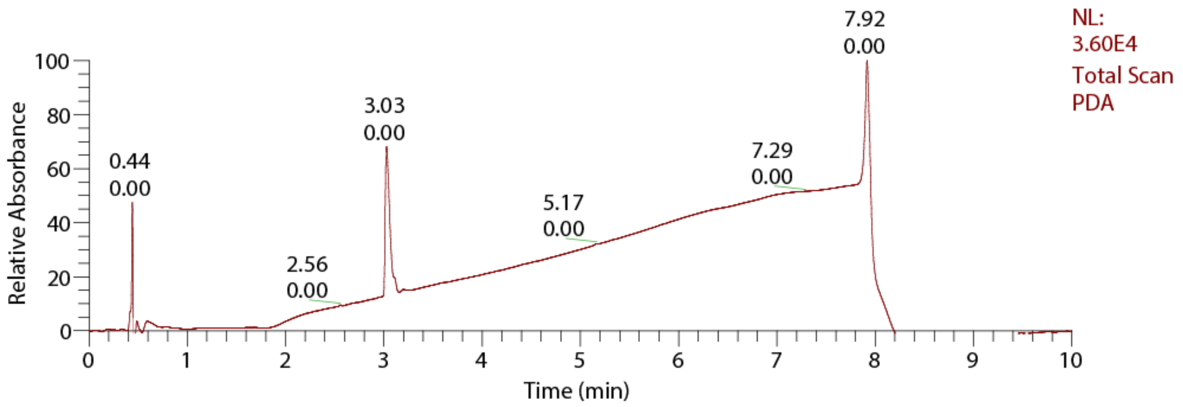

HPLC chromatogram of **FC-NAc**

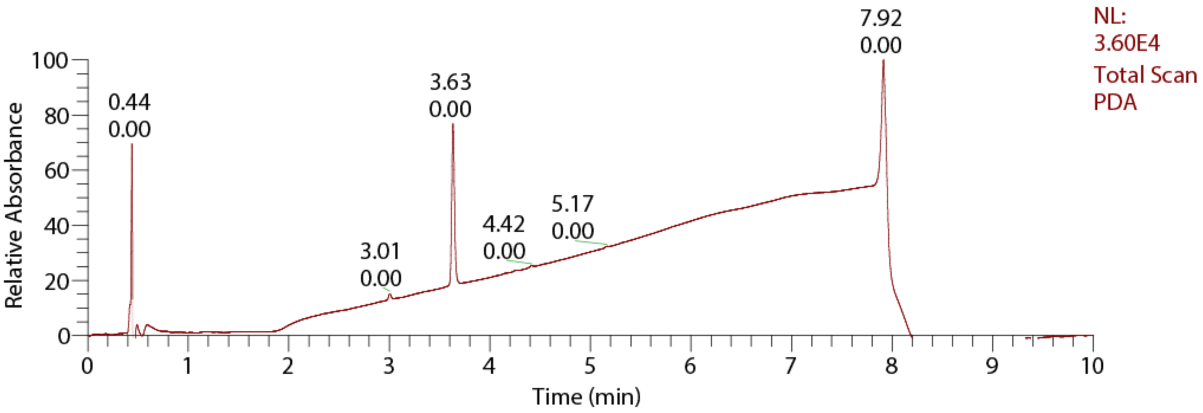

HPLC chromatogram of **VH032**

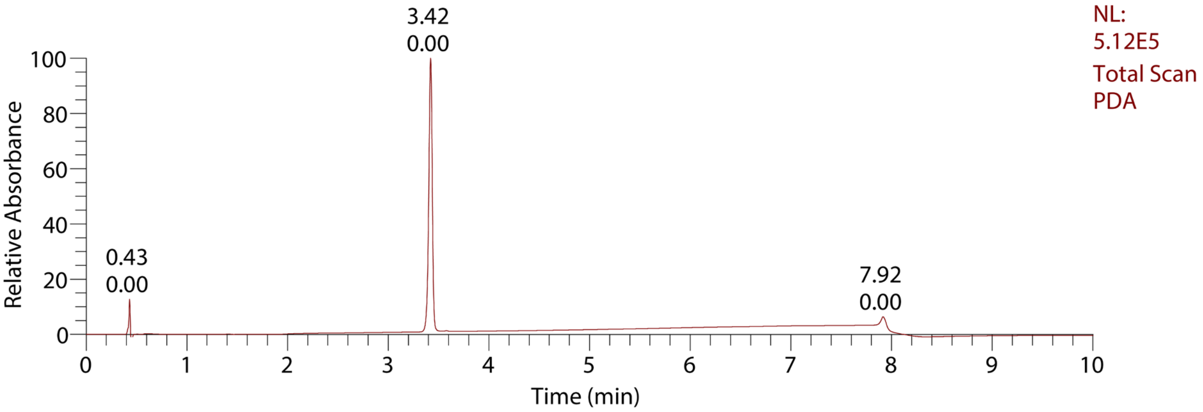

HPLC chromatogram of **2-methylbut-3-en-2-yl ether cleaved FC-NAc**

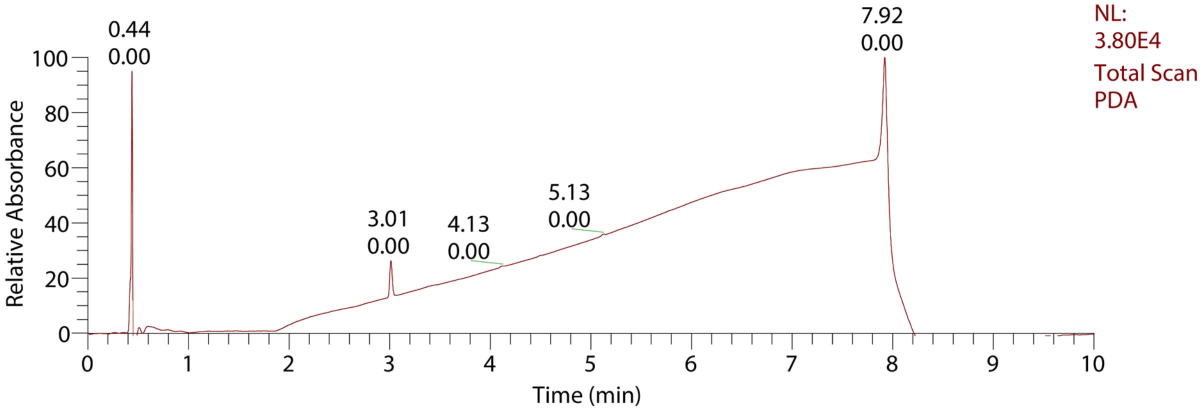

HPLC chromatogram of **CV1p**

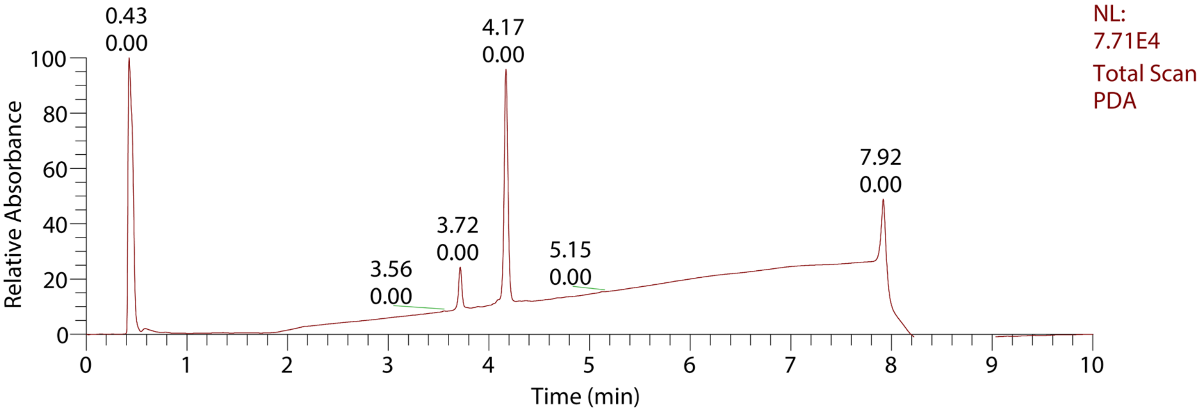

HPLC chromatogram of **CV2p**

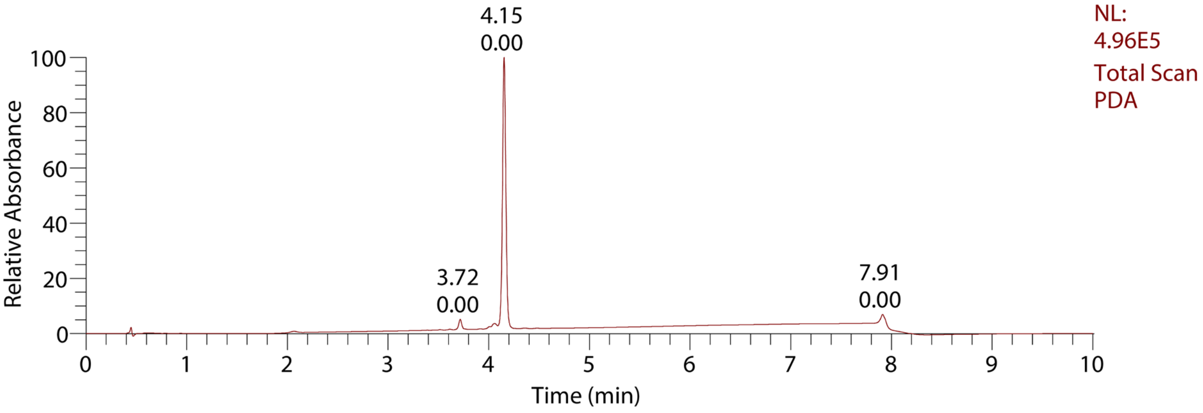

HPLC chromatogram of **CV3p**

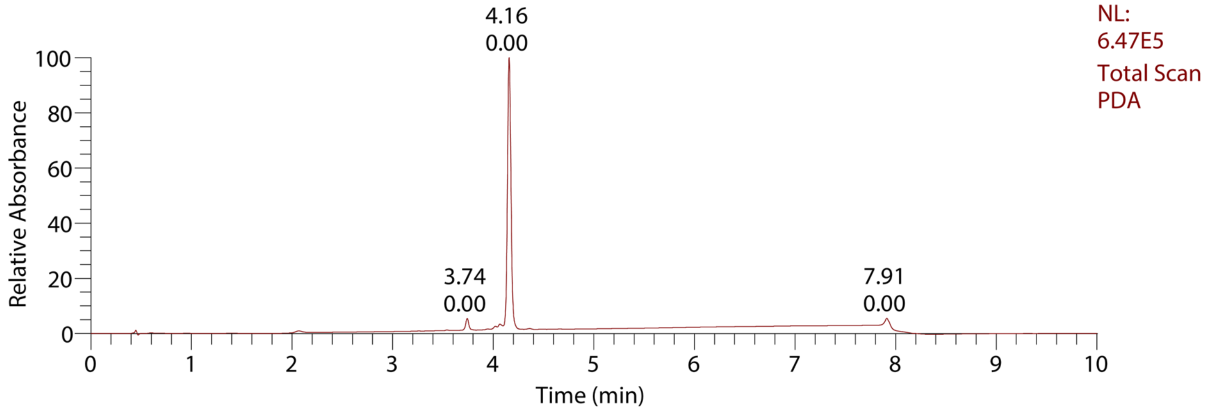

HPLC chromatogram of **CV4p**

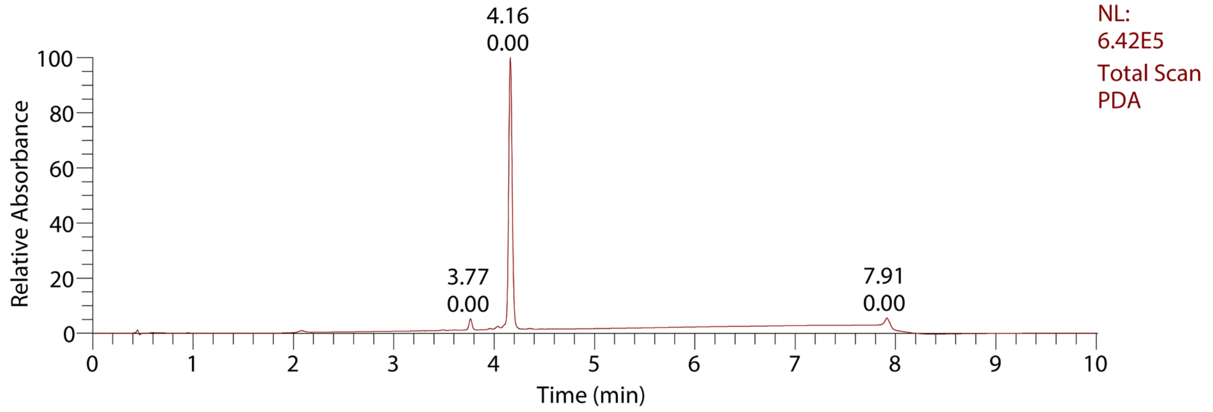

HPLC chromatogram of **CV0a**

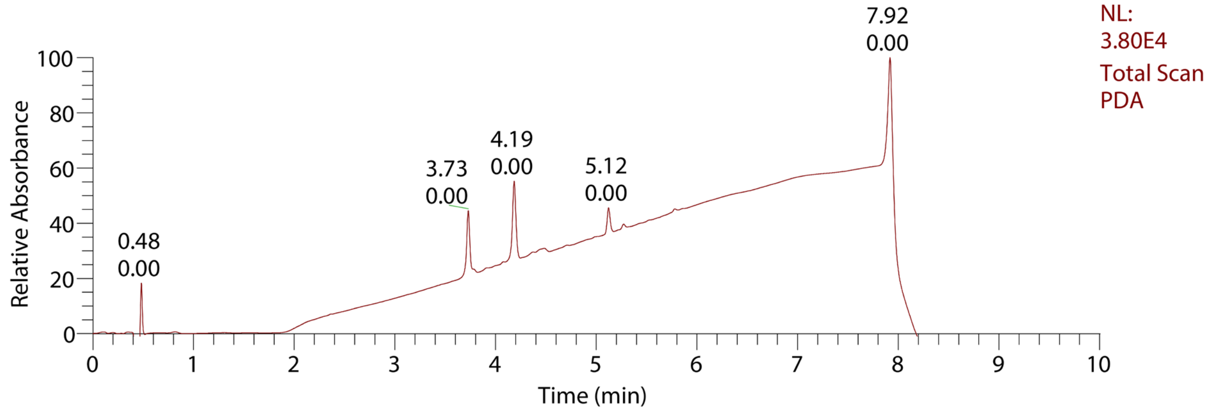

HPLC chromatogram of **CV1a**

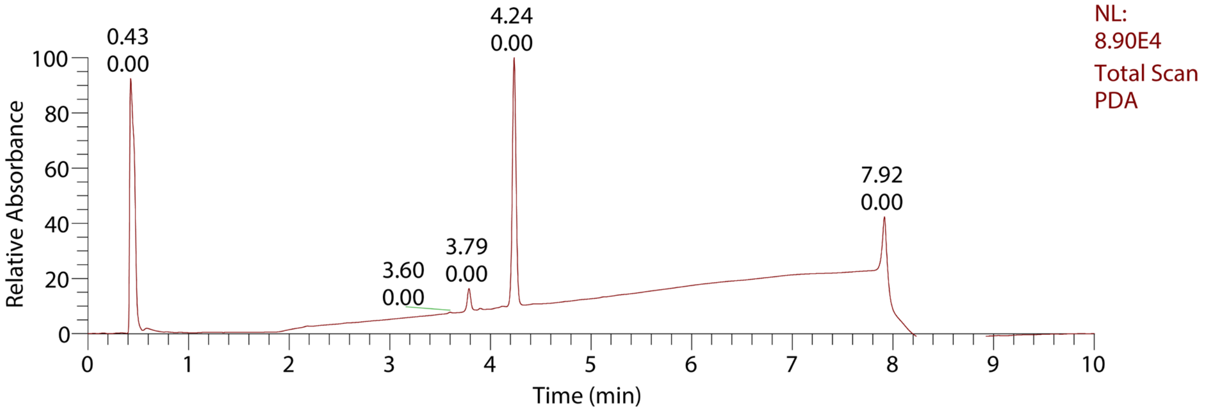

HPLC chromatogram of **CV2a**

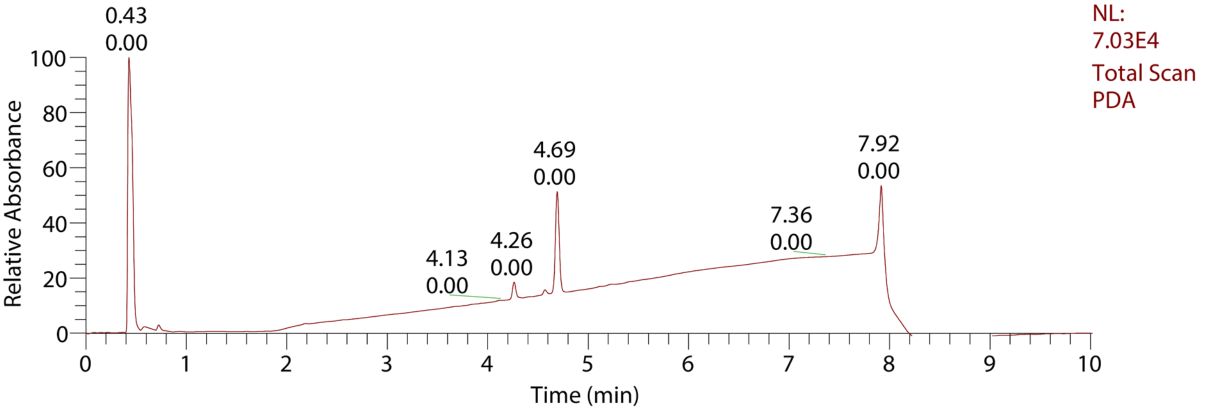

HPLC chromatogram of **CV3a**

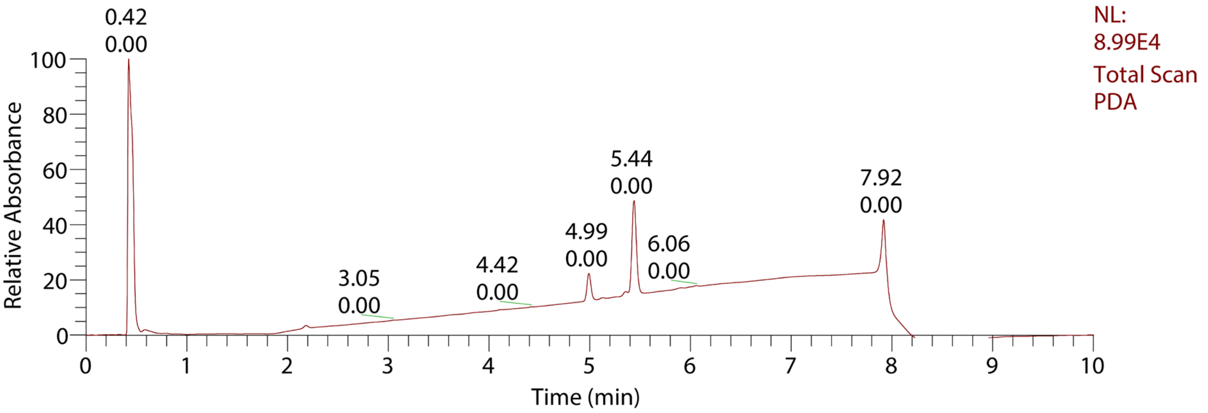

HPLC chromatogram of **(*S*-OH)-CV2a**

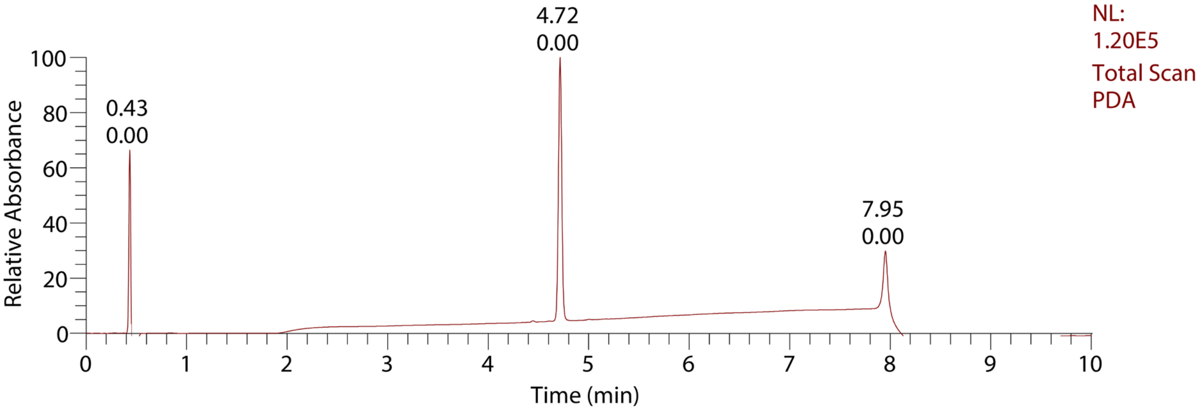
